# Noninvasive Diagnostic Ultrasound-Guided Focused Ultrasound Enables Selective, Reversible Inhibition of Peripheral Nociceptive Fibers and Prevents Acute Pain

**DOI:** 10.64898/2026.05.21.727004

**Authors:** Namrata Sangwan, Lidia Mergelian, Michael Klukinov, Morteza Mohammadjavadi, Riya Navani, Cholawat Pacharinsak, Kim Butts Pauly, Jose G. Vilches-Moure, David C. Yeomans, Thomas A. Anderson

## Abstract

**Background:** Acute postoperative pain remains a major clinical therapeutic challenge. Current peripheral nerve blockade (PNB) techniques are effective for some patients but are limited by invasiveness, short duration, reliance on highly trained providers, and off-target motor and sensory effects. Focused ultrasound (FUS) is a novel neuromodulatory technology with the potential to achieve noninvasive, selective, reversible, and prolonged inhibition of peripheral nociceptive fibers to prevent and treat acute pain. We hypothesized that noninvasive transcutaneous targeting of the rat sciatic nerve using co-aligned diagnostic ultrasound (dUS) and FUS transducers could produce selective and reversible inhibition of nociceptive pain behaviors while preserving motor and non-nociceptive sensory functions.

**Methods:** In an *in vivo* rat hindpaw incisional (HPI) pain model, using a novel, transcutaneous dUS-guided FUS system, the sciatic nerve was located with dUS, and FUS energy was applied to it just prior to hindpaw incision. FUS parameters were iteratively adjusted to achieve reversible, selective inhibition of nociceptive behaviors without changing motor and non-pain sensory behaviors. Animals were randomized into six groups: No Intervention (Control), HPI Only (Disease Control), Sham FUS, FUS Only, FUS+HPI (Intervention), and LA+HPI (Positive Control). Primary outcomes were changes in nociceptive sensory functions, assessed by thermal and mechanical sensitivity. Secondary outcomes were changes in non-nociceptive sensory and motor functions, assessed by hindpaw flexion and extension reflexes.

**Results:** Compared with the HPI Only group, the FUS+HPI group demonstrated (1) significant attenuation of hindpaw thermal hypersensitivity from day 0 - week 5.0 and week 8.0 - 16.0 (p < 0.05–0.001); (2) significant attenuation of mechanical hypersensitivity from day 0 until week 4.0 (p < 0.05–0.001); (3) no significant attenuation of flexion; and (4) no significant attenuation of extension.

**Conclusions:** Transcutaneous dUS-guided FUS enables selective, reversible inhibition of Aδ and C nociceptive fiber mediated behaviors while sparing Aα motor and Aβ sensory behaviors. FUS-induced PNB prevented both acute and persistent pain behaviors. These findings support FUS as a promising noninvasive peripheral nerve blockade strategy for acute pain management.

## Introduction

Acute pain management remains a significant clinical challenge across perioperative, inpatient, and emergency care settings. Moderate to severe acute pain is highly prevalent and frequently suboptimally controlled despite a multitude of available therapies.^1–3^ Inadequate management of acute pain is associated with increased morbidity including an increased risk of developing chronic pain.^1,3^ Acute pain arises in response to tissue injury or disease processes; postoperative and post-traumatic pain often persists for days to weeks.^4,5^ Mechanistically, acute pain results from injury-induced activation and sensitization of peripheral nociceptors, with signal transmission primarily mediated by unmyelinated C fibers, which transmit dull, pain at rest, and thinly myelinated Aδ fibers, which convey sharp, pain with movement.^4,6^

Pharmacologic management of acute pain relies heavily on systemic medications, including nonsteroidal anti-inflammatory drugs, acetaminophen, and μ-opioid receptor agonists; however, systemic analgesics frequently provide incomplete analgesia and are limited by dose-dependent adverse effects.^7,8^ Within the peripheral nervous system, opioids preferentially inhibit C fiber–mediated nociceptive transmission over that of Aδ fibers, resulting in less effective control of movement-associated pain.^6,9^ Opioid administration is additionally associated with significant short-and long-term complications, including respiratory depression, opioid-induced hyperalgesia, prolonged postoperative use, and dependence.^10,11^

Regional anesthetic techniques such as traditional needle-and local anesthetic-based peripheral nerve blocks (PNBs) improve analgesia but remain constrained by their invasiveness, limited duration, the need for highly specialized training to conduct, and local anesthetics’s lack of fiber selectivity.^12–14^ Local anesthetics nonselectively inhibit nociceptive, motor, and non-nociceptive sensory fibers, which impairs motor function, may delay rehabilitation and increase the risk of falls, and reduces patient satisfaction.^13,15^ Single-shot PNBs typically provide analgesia for less than 24 hours, whereas continuous PNBs are generally maintained for ≤4 days because of infection-related concerns, despite postoperative pain frequently persisting for substantially longer periods.^12,16^ Further, continuous catheter-based techniques increase procedural complexity and carry risks including infection and systemic toxicity.^12,16^

Focused ultrasound (FUS) has emerged as a noninvasive, image-guided modality capable of delivering spatially confined acoustic energy to targeted tissues with high spatiotemporal precision.^17,18^ At high intensities, FUS induces localized thermal ablation and is clinically utilized in conditions including prostate cancer and uterine fibroids.^19,20^ Within the central nervous system, MRI-guided FUS enables stereotactic targeting and has demonstrated clinical efficacy in movement disorders including essential tremor and Parkinson’s disease.^21–23^ At sub-ablative intensities, FUS exerts neuromodulatory effects through combined mechanical, biophysical, and thermal mechanisms, including acoustic radiation force, membrane deformation, localized tissue heating, and modulation of mechanosensitive ion channels.^24,25^ These effects alter neuronal excitability through changes in ion channel gating and membrane conductance.^25,26^ In central nervous system applications, low-intensity FUS has been shown to reversibly modulate neuronal circuits and transiently open the blood–brain barrier.^27,28^

Extending beyond the central nervous system, FUS-mediated neuromodulation of the peripheral nervous system has been demonstrated in preclinical models, where acoustic exposure alters compound action potentials, conduction velocity, and neuronal excitability.^29–33^ In rodent models of peripheral nerve injury and neuropathic pain, focused ultrasound has been shown to increase nociceptive thresholds and modulate electrophysiological responses following application to peripheral nerves including the sciatic and common peroneal nerves.^34–36^ These effects occur in an intensity-dependent manner and may preferentially affect smaller-diameter nociceptive fibers.^24,25^

Prior *ex vivo* electrophysiological studies from our group demonstrated dose-dependent focused ultrasound–induced changes to some action potential parameters similar, but not identical to, changes induced by increasing concentrations of local anesthetics, bupivacaine and ropivacaine.^38^ Additionally, *in vivo* work from our laboratory demonstrated that after invasive FUS application to the sciatic nerve, animals had reversible attenuation of hindpaw mechanical hypersensitivity, thermal hypersensitivity, flexion, and extension from 0.5 to 9 weeks and reversible nerve structural change.^37,39^

In the present study, we investigated whether diagnostic ultrasound (dUS)-guided FUS application to the sciatic nerve, administered prior to injury, can produce preferential and reversible inhibition of nociceptive (Aδ and C) fiber activity while preserving motor (Aα) and non-pain sensory (Aβ) fiber functions. Using an *in vivo* rat hindpaw incision model of pain, we evaluated the behavioral consequences of FUS application to assess its potential as a noninvasive strategy for acute pain prevention.

## Methods

### Measured Outcomes

The primary aim of this study was to evaluate the presence and duration of Aδ and C fiber inhibition following FUS application to the rat sciatic nerve in animals undergoing hindpaw incision (HPI), compared with HPI Only controls, using behavioral assays of mechanical and thermal hyperalgesia over a 16.0-week period. Secondary aims were to determine (1) the presence and duration of Aα and Aβ fiber inhibition using hindpaw extension and flexion testing, and (2) histopathological changes in the sciatic nerve following FUS application, compared with controls. To reduce possible variability in outcomes, all interventions and assessments were performed on the right hindlimb.

### *In Vivo* Study Design and Study Arms Allocation

Initially, FUS parameters (time, peak-to-peak pressure, duty cycle, waveform, pulse duration) were iteratively adjusted to maximize nociceptive (Aδ and C) fiber inhibition and minimize motor (Aα) and non-pain sensory fiber inhibition (Aβ), as determined by the behavioral testing, described below. Animals were tested as described below for 1-2 weeks to determine if nociceptive and/or motor and non-pain sensory fibers were inhibited. If all fibers were inhibited, FUS parameters were adjusted to decrease total energy delivery; if no fibers were inhibited, FUS parameters were adjusted to increase total energy delivery. Once a FUS parameter set was determined that primarily inhibited nociceptive fibers with minimal to no inhibition of motor and non-pain sensory fibers, adult male and female Sprague–Dawley rats (150–350 g; Charles River Laboratories, Wilmington, Massachusetts, USA) were randomly allocated to six experimental groups, with randomization performed to ensure balanced allocation of male and female animals within each group (n = 12 per group; 6 males and 6 females each): No Intervention (Control), HPI Only (Disease Control), Sham FUS, FUS Only, FUS+HPI (Intervention), and LA+HPI (Positive Control) (Figure 1). In the Sham FUS group, all procedures were performed identically to the FUS Only and FUS+HPI groups, except that no acoustic energy was delivered. In the Intervention group, acoustic energy was delivered to the sciatic nerve immediately prior to the hindpaw incision. In the LA+HPI group, local anesthetic was injected around the sciatic nerve immediately prior to the hindpaw incision.

**Figure 1.**
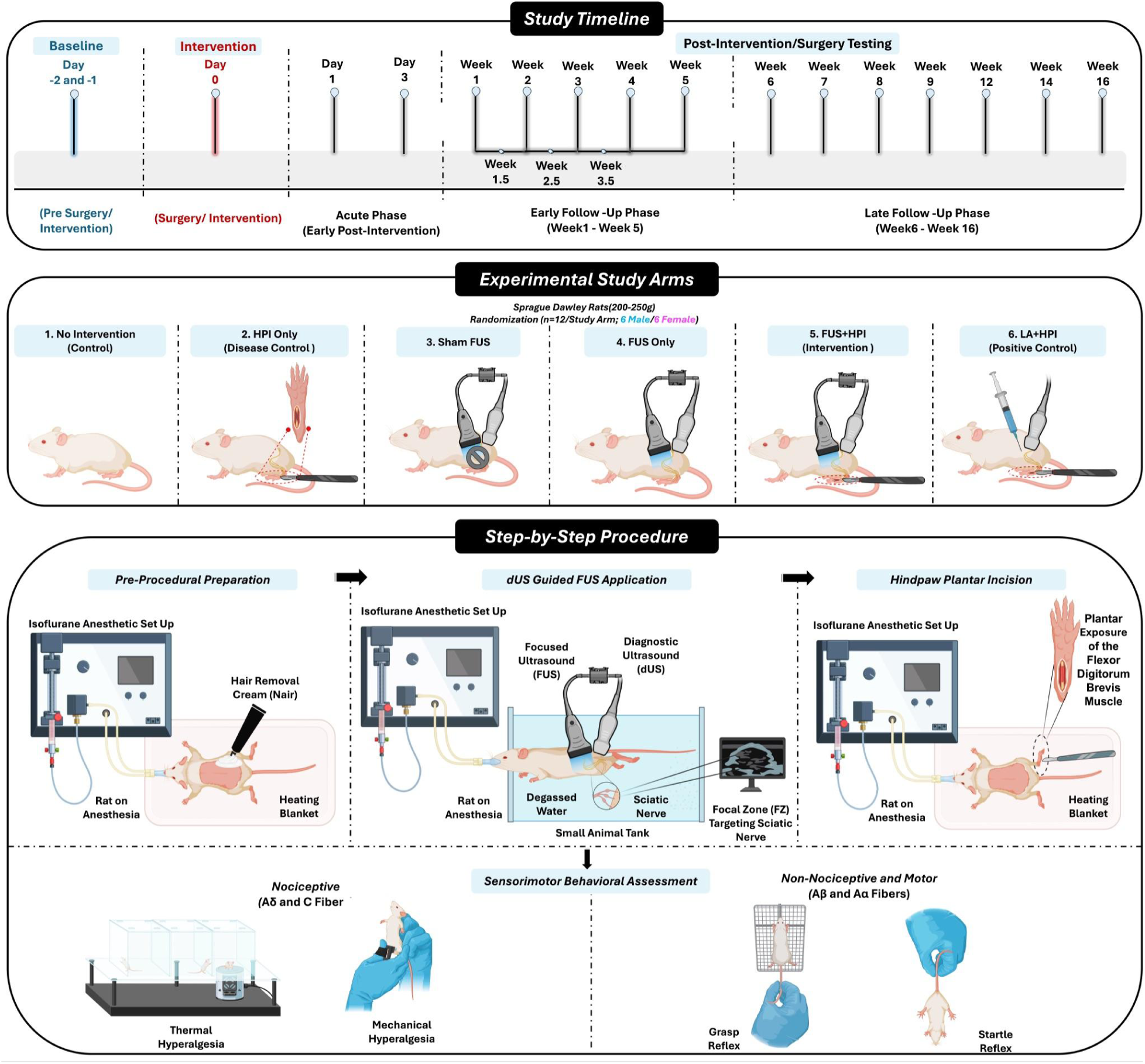
Representative illustration of study design and allocation of animals across varied study arms. **(A)** Baseline behavioral testing followed by FUS, LA, or HPI according to study arm; assessments conducted from 2 h post-intervention (day 0) through week 16. **(B)** Adult Sprague Dawley rats (150–350 g; n = 72) randomized into six study arms (n = 12 per arm; including male and female across each study arm): No Intervention (Control), HPI Only (Disease Control), Sham FUS, FUS Only, FUS+HPI (Intervention), and LA+HPI (Positive Control). **(C)** Stepwise experimental procedure showing animal preparation and positioning in the acoustic coupling setup, dUS-guided localization of the sciatic nerve, targeted FUS application, followed by hindpaw incision surgery according to study allocation, and subsequent post-intervention neurobehavioral testing. FUS, focused ultrasound; dUS, diagnostic ultrasound; LA, local anesthetic; HPI, hindpaw incision.

### Transcutaneous Focused Ultrasound (FUS) Application to Sciatic Nerve

To facilitate acoustic coupling, under inhaled general anesthesia (2% isoflurane), hair over the posterior thigh region (overlying the biceps femoris muscle) was removed, and animals were positioned in a temperature-controlled (37 °C) degassed water bath. The FUS (1.5 MHz, Sonele Inc., Markham, ON, Canada) and dUS (50 MHz, Capistrano Lab, San Clemente, CA, USA) transducers were co-aligned using a needle hydrophone (HNA 0400, ONDA Corporation, Sunnyvale, CA, USA) (Figure 2A); and the location of the-3dB focal zone was marked on the dUS screen (Figure 2B). The co-aligned dUS-FUS transducers were positioned over the posterior thigh (Figure 2C), and the sciatic nerve was visualized approximately 1 cm proximal to its trifurcation. The FUS focal zone (FZ), corresponding to the region of maximal acoustic energy, was precisely aligned with the nerve (Figure 2D), and acoustic energy was delivered.

**Figure 2.**
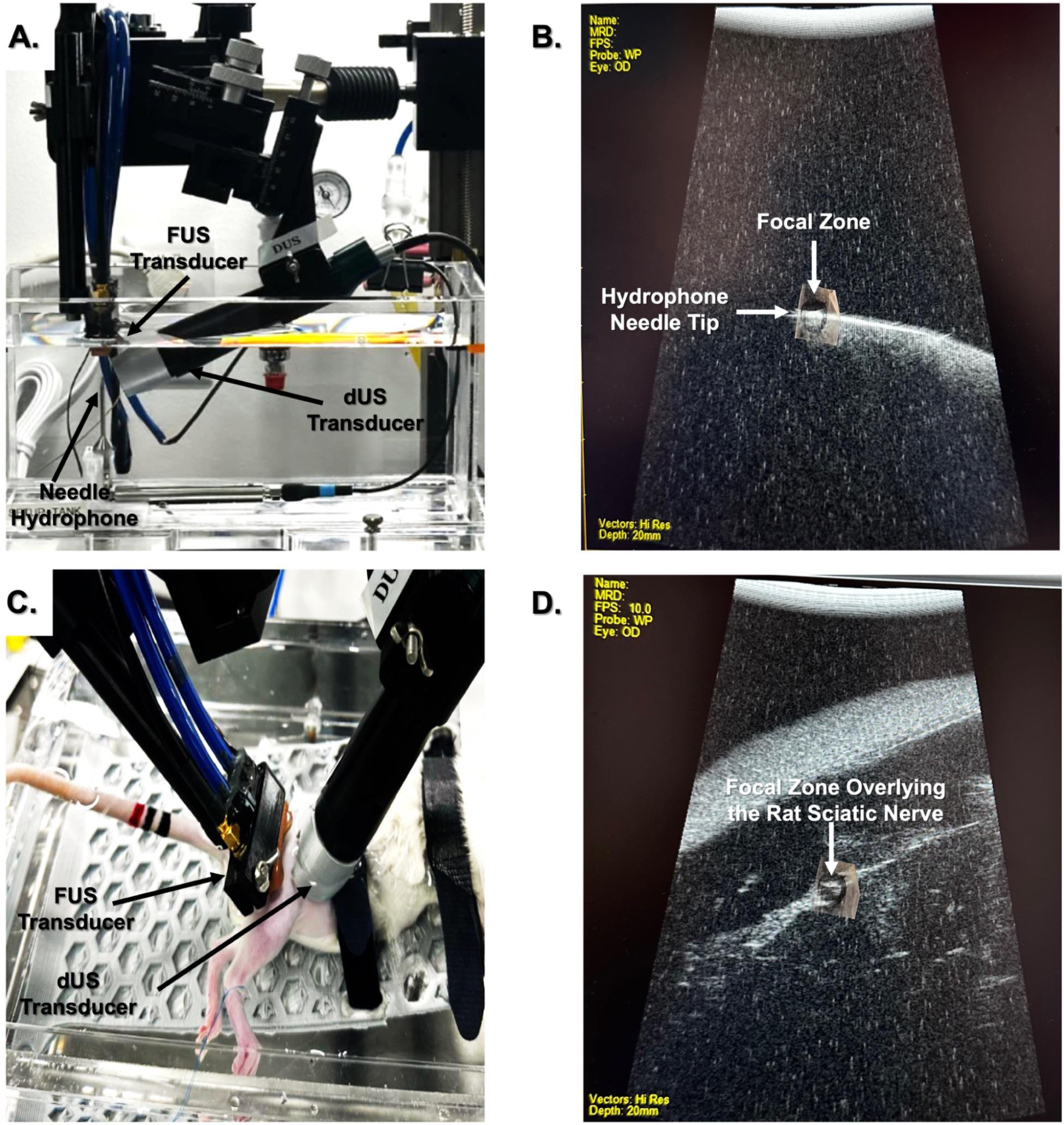
Diagnostic ultrasound (dUS)–guided focused ultrasound (FUS) system alignment, targeting, and experimental setup. **(A)** Alignment of diagnostic ultrasound (dUS) and focused ultrasound (FUS) transducers using a needle hydrophone to ensure co-registration of imaging and therapeutic focal zones. **(B)** Alignment of the needle hydrophone within the co-registered dUS–FUS system, with real-time visualization of the focal zone using UltraViewXL software via diagnostic ultrasound (dUS). **(C)** Positioning of co-aligned dUS and FUS transducers over the rat hindlimb for targeted focused ultrasound delivery to the sciatic nerve. **(D)** Representative dUS image demonstrating real-time visualization of the sciatic nerve prior to overlay of the FUS focal zone, corresponding to the region of maximal focal intensity.

### Hindpaw Incisional (HPI) Acute Pain Model

The hindpaw incisional model of acute postoperative pain was performed as originally described by Brennan *et al*.^38^ This model was utilized as it reliably produces reproducible mechanical and thermal hypersensitivity. General anesthesia was induced and maintained with inhaled isoflurane, and the plantar surface of the hindpaw was aseptically prepared with 10% povidone–iodine. A 1-cm longitudinal incision was made using a No. 11 scalpel blade through the skin and fascia of the plantar hindpaw, beginning approximately 0.5 cm proximal to the heel and extending distally toward the toes. The underlying flexor (plantaris) muscle was exposed by blunt dissection, elevated, and incised longitudinally while preserving its origin and insertion. Hemostasis was achieved with gentle pressure. The skin was closed with interrupted mattress sutures using non-absorbable 5-0 nylon, and a topical antibiotic ointment (polymyxin B, neomycin, bacitracin) was applied. If sutures remained in place, they were removed on postoperative day 14.

### Local anesthetic injection

Under dUS guidance, the sciatic nerve was identified approximately 1 cm proximal to its trifurcation and 0.2ml 0.5% bupivacaine hydrochloride (5 mg/mL; AuroMedics Pharma LLC, East Windsor, NJ) was injected adjacent to the nerve.

### Sensorimotor Behavioural Assessments

Quantitative measurements of nociceptive sensory function were obtained using established assays of thermal and mechanical sensitivity;^36, 37, 39, 40^ motor and non-nociceptive sensory function were evaluated through hindpaw extension and flexion responses.^36, 37, 41, 42^Animals underwent a 6-day acclimatization period prior to intervention to minimize stress-induced behavioral variability, including habituation to handling, the experimental environment, and behavioral testing apparatuses. Baseline behavioral assessments were performed over 2 consecutive days prior to intervention. Measurements were obtained 2 hours after emergence from general anesthesia on day 0/week 0 (day of intervention), day 1, twice weekly through week 4, once weekly from weeks 5–9, and at weeks 12, 14, and 16. Animal weight was also recorded at each time point. Behavioral assessments were conducted by three investigators (TA, NS, LM) who remained blinded to study arm allocation to the extent possible.

### Thermal Hyperalgesia (Modified Hargreaves; primarily assesses C fibers)

Thermal nociceptive sensitivity was evaluated using the hindpaw withdrawal latency to radiant heat stimulation, based on the method originally described by Hargreaves *et al.* (IITC Life Science, Woodland Hills, CA, USA).^39^ Animals were positioned individually on a transparent glass platform within a plexiglass enclosure and allowed to acclimate for 15 minutes prior to testing. To ensure consistent baseline behavior, animals were habituated to the testing environment in the plexiglass enclosures for 15 minutes daily over two consecutive days prior to the initial assessment without application of the thermal stimulus. Baseline measurements were obtained over two consecutive days prior to allocation to study arms and experimental procedures. During testing, a focused radiant heat source was applied to a defined ∼1 cm area of the proximal plantar surface of the hindpaw, and withdrawal latency was recorded. A cutoff latency of 20 seconds was implemented to prevent injury. Each trial was repeated five times with an inter-trial interval of approximately 3 minutes, and the mean latency was calculated for subsequent analysis.^39^

### Mechanical Hyperalgasia (Randall–Selitto; assesses both Aδ and C fibers)

Nociceptive withdrawal thresholds were measured using the Randall–Selitto paw pressure assay (IITC Life Science, Woodland Hills, CA, USA).^40^ In this test, a progressively increasing mechanical force is applied to the proximal plantar surface of the hindpaw until a withdrawal response or vocalization occurs. The force was increased at a controlled rate of approximately 5–10 g/s using calibrated forceps, allowing quantification on a linear scale, with a predefined cutoff of 250 g to prevent injury.

Animals were acclimated to handling by gentle interaction for 5–10 minutes daily over two consecutive days prior to testing, followed by a two-day period of habituation to the restraint cloth without application of mechanical force to the hindpaw. Baseline measurements were obtained over two consecutive days prior to allocation to study arms and experimental procedures. On each testing day, animals were handled for approximately 5 minutes before assessment to ensure consistent habituation. During testing, animals were lightly restrained in a soft cloth to limit movement. Each measurement was was repeated three times with an inter-trial interval of approximately 1 minute, and the mean value was calculated for subsequent analysis.^40^

### Hindpaw Flexion (grasp reflex; assesses both Aβ sensory and Aα motor fibers)

Motor and non-nociceptive sensory function were evaluated using the hindpaw flexion (grasp) reflex.^41^ This response is elicited when the plantar surface of the hindpaw contacts a solid substrate, resulting in flexion of the digits, and reflects the integrity of Aα motor fibers and Aβ sensory fibers, as well as proper function of the plantar flexor musculature. Animals were acclimated to handling for 5–10 minutes daily over two consecutive days prior to testing, and baseline measurements were obtained over two consecutive days prior to allocation to study arms and experimental procedures.

During assessment, rats were gently lifted and positioned such that the plantar surface of both hindpaws contacted a metal grip bar to elicit the grasp reflex. Flexion was graded using a semi-quantitative scale: 0 (no response), 0.5 (partial flexion), and 1 (full flexion).

### Hindpaw Extension (startle reflex, assesses Aα motor fibers)

Motor function was further assessed using the hindpaw extension (startle) reflex.^42^ This response is elicited following stimulation or when startled, resulting in extension of the digits and reflects the integrity of Aα motor fibers, as well as proper function of the plantar extension musculature. Animals were acclimated to handling for 5–10 minutes daily over two consecutive days prior to testing, and baseline measurements were obtained over two consecutive days prior to allocation to study arms and experimental procedures.

During assessment, rats were gently grasped around the body and lifted briefly off of the surface they were on; hindpaw digit extension was observed and graded using a semi-quantitative scale: 0 (no response), 0.5 (partial extension), and 1 (full extension).

### Statistical Analysis and Sample Size Determination

For the statistical analysis the continuous outcomes from the Randall–Selitto and modified Hargreaves assays were expressed as mean ± 95% confidence interval (CI) at each time point. Behavioral outcomes were initially analyzed using data from all animals within each treatment group and subsequently analyzed separately for male and female rats. Data were analyzed using a two-way mixed-model analysis of variance (ANOVA), with factors for treatment group and time, implemented in GraphPad Prism version 11 (GraphPad Software, San Diego, CA, USA). Where appropriate, post hoc comparisons were performed with Bonferroni correction to account for multiple testing. Statistical significance was defined as p < 0.05. Hindpaw flexion and extension outcomes were analyzed using contingency table analysis followed by Fisher’s exact test, considering the categorical nature of the response data. For hindpaw flexion and extension the behavioral responses were quantified and expressed as fractional percentages for comparative analysis across study groups and time points.

An a priori power analysis indicated that a sample size of n = 6 animals per group would provide 80% power (β = 0.8) to detect a standardized effect size of 1.25 at a significance level of α = 0.05. Twelve animals per group (6 males and 6 females) were included because two animals per group were euthanized at 6 time points for histological assessment, thus ensuring that 6 animals remained for longitudinal evaluation through the 16.0-week endpoint.

## Results

FUS was applied to approximately 140 animals during iterative adjustment of acoustic parameters to determine a FUS parameter set that achieved selective, reversible inhibition of nociceptive sensory behaviors without significant impairment of motor and non-pain sensory functions. The final optimized FUS parameter set was: spatial peak pulse average intensity of 572.72 W/cm²; mechanical index of 3.42; 60 seconds; 100% duty cycle.

### Study Arm Characteristics and Body Weight Assessment

Seventy-two rats were prospectively randomized into six experimental groups (n = 12 per group), with randomization performed to ensure balanced allocation of male and female animals within each group: Each group (No Intervention (Control), HPI Only (Disease Control), Sham FUS, FUS Only, FUS+HPI (Intervention), and LA+HPI (Positive Control)) comprised equal numbers of male and female animals (n = 6 per sex), except for the FUS+HPI group, which included 5 male and 7 female animals. The mean weight of animals in each group on Day 0 were similar (Table 1), and animals in all groups exhibited similar increases in weight over time. (Supplementary Figure 1).

**Table 1.** Study Arm Characteristics. Body weight across study arms. Data were presented as mean ± SEM. Ctrl, control; DOS, Day of Surgery; HPI, hindpaw incision; FUS, focused ultrasound; LA, local anesthetic.

| **Study Arm** | **Number of Animals**<br>**(Male/Female)** | **Body Weight on DOS**<br>**(gram, mean ± SEM)** |
| --- | --- | --- |
| No Intervention (Control) | 12 (6/6) | 149.8 ± 3.3 |
| HPI Only (Disease Control) | 12 (6/6) | 197.8 ± 20.50 |
| Sham FUS | 12 (6/6) | 221.0 ± 32.54 |
| FUS Only | 12 (6/6) | 189.2 ± 8.08 |
| FUS+HPI (Intervention) | 12 (5/7) | 189.3 ± 18.56 |
| LA+HPI (Positive Control) | 12 (6/6) | 184.9 ± 13.16 |

### Behavioral Assessment of Nociceptive and Sensorimotor Function

Qualitatively, animals in the FUS+HPI group were indistinguishable behaviorally from animals in the No Intervention group.

### Thermal Withdrawal Latency (modified Hargreaves)

#### Qualitative

Baseline values for animals in all 6 groups were approximately 15–18 seconds. HPI Only: On Day 0 (2 hours after emergence from general anesthesia), the HPI Only group had a substantial reduction in thermal withdrawal latency (to approximately 2 seconds), which then increased partially toward baseline until week 2.0 (to approximately 11 seconds), but then stayed relatively stable, and substantially less than baseline (at approximately 10-13 seconds) from week 2.0 until week 16.0.

#### FUS+HPI

On Day 0 (2 hours after emergence from general anesthesia), the FUS+HPI group had a small decrease in thermal withdrawal latency (to approximately 15 seconds), and then increased to and stayed at baseline values (approximately 16-19 seconds) from Day 1 until week 16.0, No Intervention, Sham FUS, FUS Only: There was no change in thermal withdrawal latency from baseline until week 16.0.

#### LA+HPI

On Day 0 (2 hours after emergence from general anesthesia), the LA+HPI group initially had a partial reduction in thermal withdrawal latency (to approximately 18 seconds); thermal sensitivity was reduced on Day 1 (approximately 8 seconds), and then increased until week 4.0 when it reached baseline values (approximately 16 seconds) and stayed at baseline until week 16.0. (Figure 3A, Supplementary Table 1, 2)

**Figure 3.**
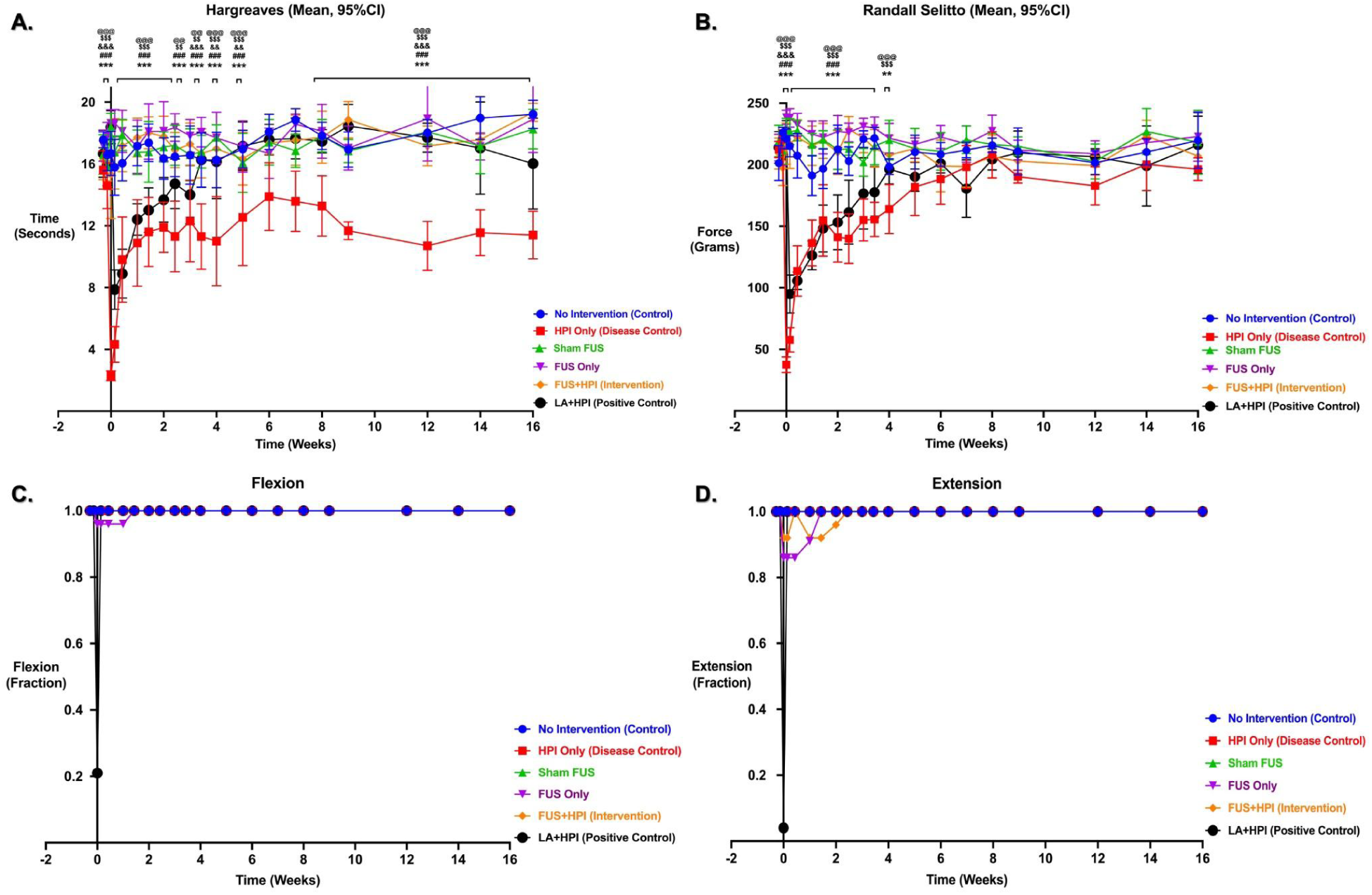
Behavioral assessment of nociceptive and sensorimotor function across study arms. Behavioral test results from the intervention and control study arms. **(A)** Thermal withdrawal latency (modified Hargreaves test). **(B)** Mechanical withdrawal threshold (Randall–Selitto test). **(C)** Hindpaw (HP) flexion (grasp) reflex. **(D)** Hindpaw (HP) extension (startle) reflex. **Panels A and B:** Data are presented as mean ± 95% confidence interval (CI). Statistical comparisons were performed using a two-way mixed-effects ANOVA with fixed effects for study arm and time, followed by Bonferroni-corrected post hoc testing. Statistical comparisons were conducted relative to the HPI Only (Disease Control) study arm. Significance is indicated as follows: ns (p ≥ 0.05), * (p < 0.05), ** (p < 0.01), *** (p < 0.001). **Panels C and D:** Data were analyzed using contingency table analysis followed by Fisher’s exact test, and behavioral responses are presented as fractional response values. Significance indicators: Control-$, Sham FUS-@, FUS Only-*, FUS+HPI (Intervention)-#, and LA+HPI (Positive Control)-&, all relative to the HPI Only (Disease Control) study arm. FUS, focused ultrasound; dUS, diagnostic ultrasound; HPI, hindpaw incision; LA, local anesthetic.

#### Quantitative

Compared to the HPI Only group: The FUS+HPI group had significantly prolonged thermal withdrawal latencies from Day 0–week 5.0 and week 8.0–16.0 (p = 0.003 to p < 0.001).

No Intervention, Sham FUS, and FUS Only had significantly prolonged thermal withdrawal latencies from Day 0–week 5.0 and week 8.0–16.0 (p < 0.05 to p < 0.001).

LA+HPI group had significantly prolonged thermal withdrawal latencies from Day 0–Day 1 (p = 0.009 to p < 0.001), with loss of significance from Day 3–week 2.0 (p = 0.291 to p > 0.999). Significance re-emerged at week 2.5 (p = 0.002), was again absent at week 3.0 (p = 0.716), and then remained significant from week 3.5–16.0 (p = 0.013 to p < 0.001)

Compared to the No Intervention group: FUS+HPI group had significantly reduced thermal withdrawal latencies at Day 0 (p = 0.003) with no significant differences observed from Day 1 through week 16.0. (Figure 3A, Supplementary Table 1, 2)

#### Male Rats

Compared with the HPI Only group, the No Intervention, Sham FUS, FUS Only, and FUS+HPI groups demonstrated significantly prolonged thermal withdrawal latencies from Day 0–week 5.0 and at weeks 7.0–9.0 (p < 0.05 to p < 0.001). The LA+HPI group demonstrated significant attenuation at Day 0–Day 1 and at weeks 3.5–9.0, with loss of significance at several intermediate and later time points (p = 0.048 to p < 0.001). (Supplementary Figure 2A, Supplementary Table 3, 4)

#### Female Rats

Compared with the HPI Only group, the No Intervention, Sham FUS, FUS Only, and FUS+HPI groups demonstrated significantly prolonged thermal withdrawal latencies from Day 0–week 2.5 and at weeks 9.0–12.0 (p < 0.05 to p < 0.001). The LA+HPI group demonstrated significant attenuation at Day 0, week 2.5, and weeks 9.0–12.0, with loss of significance at several intermediate and later time points (p < 0.05 to p < 0.001). (Supplementary Figure 3A, Supplementary Table 5, 6)

Overall, no substantial sex-based differences in thermal withdrawal latency outcomes were observed between male and female rats.

### Mechanical Withdrawal Threshold

#### Qualitative

Baseline values for animals in all 6 groups were approximately 180–240 g. HPI Only: On Day 0 (2 hours after emergence from general anesthesia), the HPI Only group had a substantial reduction in mechanical withdrawal threshold (to approximately 30–45 g), which then partially recovered toward baseline until week 3.5 (to approximately 150–170 g), but remained below baseline values through approximately week 6.0. Mechanical withdrawal thresholds subsequently approached baseline values from weeks 7.0–16.0.

#### FUS+HPI

On Day 0 (2 hours after emergence from general anesthesia), the FUS+HPI group demonstrated only a minimal reduction in mechanical withdrawal threshold (to approximately 215–220 g), and thresholds remained near baseline values (approximately 200–230 g) throughout the study period.

No Intervention, Sham FUS, and FUS Only: There was no substantial change in mechanical withdrawal thresholds from baseline until week 16.0.

#### LA+HPI

On Day 0 (2 hours after emergence from general anesthesia), the LA+HPI group initially demonstrated a partial reduction in mechanical withdrawal threshold (to approximately 225–230 g), followed by a more pronounced reduction on Day 1 (approximately 80–110 g). Mechanical withdrawal thresholds then gradually increased toward baseline values by approximately weeks 4.0–6.0 and remained near baseline thereafter. (Figure 3B, Supplementary Table 7, 8)

#### Quantitative

Compared to the HPI Only group: The FUS+HPI group had significantly prolonged mechanical withdrawal thresholds from Day 0–week 5.0 (p = 0.040 to p < 0.001), with no significant differences observed thereafter.

No Intervention, Sham FUS, and FUS Only groups had significantly prolonged mechanical withdrawal thresholds from Day 0–week 6.0 (p < 0.05 to p < 0.001), with no significant differences observed thereafter.

The LA+HPI group demonstrated significant attenuation at Day 0 and week 4.0 in male rats and at Day 0 only in female rats, with loss of significance at most intermediate and later time points (p = 0.008 to p < 0.001).

Compared to the No Intervention group: The FUS+HPI group had significantly reduced mechanical withdrawal thresholds at Day 0–week 3.5 (p = 0.010 to p < 0.001), with no significant differences observed thereafter. (Figure 3B, Supplementary Table 7, 8)

#### Male Rats

Compared with the HPI Only group, the No Intervention, Sham FUS, FUS Only, and FUS+HPI groups demonstrated significantly prolonged mechanical withdrawal thresholds from Day 0–week 5.0 (p < 0.05 to p < 0.001). The LA+HPI group demonstrated significant attenuation at Day 0–Day 1 and week 4.0, with loss of significance at most remaining time points (p = 0.008 to p < 0.001). (Supplementary Figure 2B, Supplementary Table 9, 10)

#### Female Rats

Compared with the HPI Only group, the No Intervention, Sham FUS, FUS Only, and FUS+HPI groups demonstrated significantly prolonged mechanical withdrawal thresholds from Day 0–week 3.5 (p < 0.05 to p < 0.001). The LA+HPI group demonstrated significant attenuation only at Day 0, with no significant differences observed thereafter (p < 0.001). (Supplementary Figure 3B, Supplementary Table 11, 12)

Overall, no substantial sex-based differences in mechanical withdrawal threshold outcomes were observed between male and female rats.

### Hindpaw Flexion (Grasp Reflex)

#### Qualitative

Baseline values for all animals in all 6 groups were 1.0 (grasp reflex intact). No Intervention, HPI Only, Sham FUS, and FUS+HPI: All animals (12/12 per group) demonstrated preserved hindpaw flexion responses at all assessed time points throughout the 16-week observation period, with no observable reductions in grasp reflex responses.

#### FUS Only

11/12 animals had preserved hindpaw flexion responses throughout the study duration. One animal had a reduction in flexion response on day 0–day 1, followed by recovery to baseline flexion response (score = 1.0) from week 0.5 onward and maintained through week 16.0.

#### LA+HPI

There was marked transient suppression of hindpaw flexion responses immediately following local anesthetic administration at week 0, followed by recovery to baseline flexion responses from day 1 onward and maintained through week 16.0. (Figure 3C; Supplementary Table 13)

#### Quantitative

Compared to the HPI Only group, No Intervention, Sham FUS, and FUS+HPI groups demonstrated preserved hindpaw flexion responses throughout all assessed time points, with no significant impairments observed.

The FUS Only group demonstrated transient mild reduction in hindpaw flexion responses during the acute post-intervention period at day 0-day 1 (mean score = 0.96); however, these changes were not statistically significant, and complete recovery to baseline responses was observed from week 0.5 onward.

The LA+HPI group demonstrated significant transient suppression of hindpaw flexion responses immediately following local anesthetic administration at day 0, followed by recovery to baseline responses from day 1 onward (Figure 3C; Supplementary Table 13).

#### Male Rats

Compared to the HPI Only group, male animals in the No Intervention, Sham FUS, and FUS+HPI groups maintained preserved hindpaw flexion responses throughout the study duration, with no significant impairments observed. The male FUS Only group demonstrated transient mild reduction in flexion responses during the acute post-intervention period at day 0– week 1.0 (mean score = 0.75–0.83); however, these changes were not statistically significant, followed by complete recovery thereafter. The male LA+HPI group demonstrated significant transient suppression of flexion responses at day 0 (mean score = 0.083), followed by recovery to baseline responses from day 1 onward (Supplementary Figure 2C; Supplementary Table 14).

#### Female Rats

Compared to the HPI Only group, female animals in the No Intervention, Sham FUS, FUS Only, and FUS+HPI groups maintained preserved hindpaw flexion responses throughout nearly all assessed time points, with no significant impairments observed. The female LA+HPI group demonstrated significant transient suppression of flexion responses at day 0 (mean score = 0.08), followed by recovery to baseline responses from day 1 onward (Supplementary Figure 3C; Supplementary Table 15).

Overall, no substantial sex-based differences in hindpaw flexion responses were observed between male and female rats over the 16-week study period.

### Hindpaw Extension (Startle Reflex)

#### Qualitative

Baseline values for all animals in all 6 groups were 1.0 (extension reflex intact).

No Intervention, HPI Only, and Sham FUS: All animals (12/12 per group) maintained preserved hindpaw extension responses at all assessed time points throughout the 16-week observation period, consistent with preserved Aα motor fiber-mediated function.

#### FUS Only

10/12 animals maintained preserved hindpaw extension responses throughout the study duration. Transient mild reductions in extension responses were observed during the acute post-intervention period at day 0– week 0.5, followed by complete recovery to baseline extension responses by approximately week 1.0 and maintained through week 16.0. No persistent or delayed motor deficits were identified.

#### FUS+HPI

11/12 animals maintained preserved hindpaw extension responses at all assessed time points, while approximately 1/12 animal demonstrated transient reduction in extension responses during the acute post-intervention period, followed by recovery to baseline responses by approximately week 1.5–2.0 with preserved responses thereafter.

#### LA+HPI

There was marked transient suppression of hindpaw extension responses immediately following local anesthetic administration at day 0, followed by rapid recovery to baseline extension responses by week 0.5 and maintained through week 16.0 (Figure 3D; Supplementary Table 16).

#### Quantitative

Compared to the HPI Only group: No Intervention and Sham FUS groups demonstrated preserved hindpaw extension responses throughout all assessed time points (mean score = 1), with no significant impairments observed.

FUS Only group demonstrated transient mild reduction in hindpaw extension responses during the acute post-intervention period at day 0–week 0.5 (mean score = 0.88–0.92), followed by complete recovery to baseline responses (mean score = 1) from approximately week 1.0 onward. These changes were not statistically significant.

FUS+HPI group demonstrated transient mild reduction in hindpaw extension responses during the acute post-intervention period (mean score = 0.92–0.96), followed by recovery to baseline responses (mean score = 1) by approximately week 1.5–2.0 and preserved responses thereafter. These changes were not statistically significant.

LA+HPI group demonstrated complete transient suppression of hindpaw extension responses immediately following local anesthetic administration at day 0 (mean score = 0.04), followed by recovery to baseline responses (mean score = 1) by approximately week 0.5. These changes were statistically significant (Figure 3D; Supplementary Table 16).

#### Male Rats

Compared to the HPI Only group, male animals in the No Intervention, Sham FUS, and FUS+HPI groups maintained preserved hindpaw extension responses throughout the study duration (mean score = 1). In the male FUS Only group, transient reduction in extension responses was observed during the acute post-intervention period (mean score = 1.0), followed by complete recovery thereafter; these changes were not statistically significant. The male LA+HPI group demonstrated transient suppression of extension responses immediately following local anesthetic administration (mean score = 0.00) with subsequent recovery to baseline responses; these changes were statistically significant (Supplementary Figure 2D; Supplementary Table 17).

#### Female Rats

Compared to the HPI Only group, female animals in the No Intervention, Sham FUS, and FUS Only groups maintained preserved hindpaw extension responses throughout nearly all assessed time points (mean score = 1). In the female FUS+HPI group, transient reduction in extension responses was observed during the acute post-intervention period (mean score = 0.88), followed by complete recovery without persistent impairment; these changes were not statistically significant. The female LA+HPI group demonstrated transient suppression of extension responses immediately following local anesthetic administration (mean score = 0.33) with recovery thereafter; these changes were statistically significant (Supplementary Figure 3D; Supplementary Table 18).

Overall, no substantial sex-based differences in hindpaw extension responses were observed between male and female rats throughout the 16-week study period. These findings support preservation of motor (Aα) fiber-associated function following FUS application.

## Discussion

Using a novel dUS-FUS platform and optimized FUS parameters, noninvasive application of acoustic energy to the sciatic nerve in an animal model of soft tissue injury produced sustained peripheral nerve blockade that prevented both acute and persistent pain behaviors for 16 weeks while preserving motor and non-nociceptive sensory functions.

Poorly controlled acute postoperative pain is associated with delayed recovery, impaired rehabilitation, increased opioid exposure, and elevated risk of developing chronic pain.^1,4,6^ Traditional peripheral nerve blockade techniques can provide effective analgesia but remain limited by their invasiveness, relatively short duration (<24 hours for single shot, typically maximum of 4 days for continuous due to increasing infection risk), the need for significant training to conduct, and the nonselective inhibition of nociceptive and non-nociceptive fibers from commonly used local anesthetics.^15^ Inhibition of peripheral nerve Aα, motor, fibers results in weakness, can impair patient mobility, prevents neurological examination immediately after surgery, and increase the risk of patient falls. Inhibition of peripheral nerve Aβ, non-pain sensory, fibers results in numbness, reducing patient satisfaction, and prevents neurological examination immediately after surgery.^15,16^

A nonsystemic, noninvasive modality capable of relatively long duration (weeks), selective nociceptive fiber inhibition which requires less training (only nerve identification with dUS, not driving a needle to a nerve) would have multiple advantages over systemic analgesics and traditional PNBs.

Previous preclinical investigations in which FUS was applied to the peripheral nervous system, have demonstrated parameter-dependent modulation of peripheral nerve conduction across various experimental models. Tsui et al. reported enhanced conduction velocity in bullfrog sciatic nerve using 3.5 MHz ultrasound at 1–3 W intensity assessed by compound action potential recordings.^48^ Ilham et al. demonstrated increased conduction velocity in A-and C-type axons in mouse sciatic nerve ex vivo following 1.1 MHz focused pulsed ultrasound exposure (0.91–28.2 W/cm²; 20–40% duty cycle) using single-unit electrophysiological recordings.^49^ In contrast, Colucci et al. demonstrated temporary conduction block in bullfrog sciatic nerve using continuous-wave HIFU (0.66–1.98 MHz; 370 W/cm²).^29^ Foley et al. suggested peripheral nerve conduction block in rabbit sciatic nerve using image-guided high-intensity focused ultrasound.^30^ Furthermore, they demonstrated partial and transient sciatic nerve conduction block in rat models using high-intensity focused ultrasound paradigms (3.2–5.7 MHz; up to 1930 W/cm²), although higher exposure settings produced histological evidence of demyelination and axonal injury.^31, 32^ Importantly, Lee et al. demonstrated that high-intensity focused ultrasound (2.68 MHz; 2290–2810 W/cm²) produced temporary and reversible blockade of compound action potentials in the sciatic nerve and sensory action potentials in the sural nerve of normal and diabetic rats, with recovery occurring within 10–30 min following stimulation, supporting the potential application of focused ultrasound for reversible sensory nerve blockade and peripheral analgesia.^50^

The present findings support selective modulation of nociceptive processing without global suppression of peripheral nerve function, consistent with preferential effects on smaller-diameter nociceptive fibers relative to larger myelinated sensorimotor fibers. Thermal nociception assessed by the Hargreaves assay primarily reflects unmyelinated C fiber activity, whereas Randall–Selitto mechanical thresholds reflect both C fiber and thinly myelinated Aδ fiber activities.

Young et al. demonstrated reversible modulation of frog sciatic nerve conduction using focused ultrasound, with greater effects observed in smaller C fibers relative to larger A fibers following short-duration ultrasound exposure (0.4–1.0 s).^46^ Prior studies further suggest that smaller unmyelinated C fibers exhibit greater sensitivity to focused ultrasound stimulation than larger Aα fibers, potentially reflecting electro-mechanical mechanisms underlying ultrasound-mediated neuromodulation.^43, 44^ Mechanistically, focused ultrasound–induced neuromodulation is thought to involve acoustic radiation force, membrane deformation, and modulation of mechanosensitive ion channels, resulting in altered neuronal excitability.^24,25^ Prior peripheral ultrasound neuromodulation studies further suggest that acoustic forces may alter membrane conductance, ion channel kinetics, and neuronal excitability in a reversible and intensity-dependent manner, thereby modulating nociceptive transmission without requiring permanent neural disruption.^24, 25, 47^

Clinical and translational investigations have similarly demonstrated ultrasound-mediated modulation of peripheral sensory and pain-related pathways. Previous studies have reported that ultrasound exposure can alter sensory and motor nerve latencies in healthy human median nerves, with these latency alterations appearing to be related to temperature changes induced by ultrasound’s thermal effects.^51^ Image-guided high-intensity focused ultrasound (HIFU) for large animal nerve ablation demonstrated in a pig model that ultrasound-guided HIFU could achieve targeted noninvasive peripheral nerve blockade, with significant reductions in compound action potential (CAP) amplitudes following femoral nerve sonication, supporting the potential of focused ultrasound for pain-related neuromodulation applications.^52^ Dababou et al. reviewed the clinical use of high-intensity focused ultrasound (HIFU) for cancer-related pain management, highlighting its role as a noninvasive image-guided therapy for pain palliation through local tissue denervation, tumor mass reduction, and neuromodulation of pain-related pathways.^53^ di Biase et al. highlighted focused ultrasound (FUS) as an emerging noninvasive strategy for chronic pain management, including MRgHIFU treatment of lumbar facet joint osteoarthritis through thermal ablation of facet joint nerve terminals and medial branch nerves, with additional potential applications in peripheral nerve ablation, phantom limb pain, knee osteoarthritis, and other refractory pain conditions. The authors further emphasized that the unique noninvasive nature of FUS, combined with its neuromodulatory and neuroablative capabilities, highlights its significant translational potential for chronic pain therapeutics.^54^ In addition, Lee et al. demonstrated in healthy human subjects that ultrasound imaging-guided 1.1 MHz pulsed focused ultrasound could target the median nerve and alter thermal pain perception. Using real-time displacement imaging, they confirmed nerve engagement during sonication and reported reduced needle-like thermal pain ratings during on-target FUS compared with sham or off-target stimulation, supporting the translational potential of focused ultrasound for noninvasive peripheral pain modulation.^55^

Despite the promising translational potential of noninvasive dUS-guided FUS-induced peripheral nerve blockade for pain management, this study has several limitations that warrant consideration. Although FUS preferentially attenuated nociceptive hypersensitivity, mild transient effects on motor and non-pain sensory function were also observed, suggesting that further optimization of acoustic parameters may be necessary to improve fiber selectivity. In addition, the behavioral assays used in this study represent indirect surrogate measures of peripheral nerve fiber function. Further large-animal and clinical studies are therefore needed to establish the safety, reproducibility, and clinical feasibility of this approach. Translation to humans may also be influenced by interindividual variability, and identifying acoustic parameters that selectively inhibit nociceptive fibers while preserving motor and non-pain sensory function may prove challenging. Moreover, application of FUS to peripheral nerves may produce procedural discomfort or pain; however, similar to conventional peripheral nerve blockade procedures, sedation and analgesia with agents such as Fentanyl and Midazolam may be sufficient for clinical application. Notably, no animal movement was observed during FUS application under relatively light inhaled general anesthesia, including across our cumulative experience in >300 rats.

Taken together, these findings support the concept that noninvasive dUS-guided FUS can produce selective and reversible attenuation of Aδ and C fiber-mediated nociceptive hypersensitivity while largely preserving Aα motor and Aβ non-pain sensory function, highlighting its potential as a future peripheral neuromodulatory strategy for pain management.

## Conclusions

In a preclinical model of pain, a single, short, noninvasive FUS application to the sciatic nerve, prior to injury, prevented both acute and persistent pain behaviors, while preserving motor and non-pain sensory behaviors. These findings suggest that FUS holds promise as a clinical tool to prevent postoperative pain and the development of persistent postsurgical pain. Future investigations are necessary to determine FUS parameters that may produce reversible and selective inhibition of peripheral nerve nociceptive fibers in humans.

## Contributors

NS and TAA performed all experiments, analyzed the data, and wrote the manuscript. LM and RN performed *in vivo* behavioural testing. MM and MK assisted with experimental procedures, including ultrasound setup and troubleshooting, and contributed to data analysis. KBP, CP, JGVM, and DY contributed to research conceptualization and interpretation of the data. All authors contributed to manuscript preparation, critically reviewed the content, and approved the final version of the manuscript. TAA conceptualized and designed all experiments and provided final approval of the manuscript.

## Disclaimer

The content is solely the responsibility of the authors and does not necessarily represent the official views of the NIH.

## Ethical Approval

Ethical approval for this experimental study was obtained from the Administrative Panel on Laboratory Animal Care at Stanford University (Protocol No. 33489 & 34494). All procedures involving rats were conducted in accordance with the Guide for the Care and Use of Laboratory Animals in a facility accredited by the Association for Assessment and Accreditation of Laboratory Animal Care (AAALAC) International.

## Data Availability

Data supporting the findings of this study are available from the corresponding author upon request.

## Abbreviations

ANOVA: analysis of variance
CAP: compound action potential
CI: confidence interval
CNS: central nervous system
dUS: diagnostic ultrasound
FUS: focused ultrasound
FZ: focal zone
HIFU: high-intensity focused ultrasound
HP: hindpaw
HPI: hindpaw incision
LA: local anesthetic
MI: mechanical index
MRI: magnetic resonance imaging
PNB: peripheral nerve block
POD: postoperative day.

## References

1. Gan TJ. Poorly controlled postoperative pain: prevalence, consequences, and prevention. J Pain Res. 2017 Sep 25;10:2287–2298. doi: 10.2147/JPR.S144066.

2. Gan TJ, Habib AS, Miller TE, White W, Apfelbaum JL. Incidence, patient satisfaction, and perceptions of post-surgical pain: results from a US national survey. Curr Med Res Opin. 2014;30(1):149–160.

3. Hackworth JC, Schneider JE, Do Valle M, Fam D, Argoff C, Offidani E, Potenziano J. The burden of acute pain in the U.S. in the wake of the opioid crisis. Front Pain Res (Lausanne). 2025;6:1642035.

4. Karcz M, Abd-Elsayed A, Chakravarthy K, Aman MM, Strand N, Malinowski MN, et al. Pathophysiology of Pain and Mechanisms of Neuromodulation: A Narrative Review (A Neuron Project). J Pain Res. 2024;17:3757–3790.

5. Khammissa RAG, Ballyram R, Fourie J, Bouckaert M, Lemmer J, Feller L. Selected pathobiological features and principles of pharmacological pain management. J Int Med Res. 2020;48(5):300060520903653.

6. Heinke B, Gingl E, Sandkühler J. Multiple targets of μ-opioid receptor-mediated presynaptic inhibition at primary afferent Aδ-and C-fibers. J Neurosci. 2011;31(4):1313–1322.

7. Ong CK, Lirk P, Tan CH, Seymour RA. An evidence-based update on nonsteroidal anti-inflammatory drugs. Clin Med Res. 2007;5(1):19–34.

8. Alorfi NM. Pharmacological Methods of Pain Management: Narrative Review of Medication Used. Int J Gen Med. 2023;16:3247–3256.

9. Lu Y, Sweitzer SM, Laurito CE, Yeomans DC. Differential opioid inhibition of C-and A delta-fiber mediated thermonociception after stimulation of the nucleus raphe magnus. Anesth Analg. 2004;98(2):414–419.

10. Lawal OD, Gold J, Murthy A, et al. Rate and Risk Factors Associated With Prolonged Opioid Use After Surgery: A Systematic Review and Meta-analysis. JAMA Netw Open. 2020;3(6):e207367.

11. Durand Z, Nechuta S, Krishnaswami S, Hurwitz EL, McPheeters M. Prevalence and Risk Factors Associated With Long-term Opioid Use After Injury Among Previously Opioid-Free Workers. JAMA Netw Open. 2019;2(7):e197222.

12. Capdevila X, Bringuier S, Borgeat A. Infectious risk of continuous peripheral nerve blocks. Anesthesiology. 2009;110(1):182–188.

13. Becker DE, Reed KL. Local anesthetics: review of pharmacological considerations. Anesth Prog. 2012;59(2):90–101.

14. Ilfeld BM, Duke KB, Donohue MC. The association between lower extremity continuous peripheral nerve blocks and patient falls after knee and hip arthroplasty. Anesth Analg. 2010;111(6):1552–1554.

15. Joshi G, Gandhi K, Shah N, Gadsden J, Corman SL. Peripheral nerve blocks in the management of postoperative pain: challenges and opportunities. J Clin Anesth. 2016;35:524–529.

16. Wang YM, Liang JN, Liu DC, Zhao BC, Zhang H, Zhang HF. Factors associated with postoperative rebound pain after peripheral nerve block anaesthesia for orthopedic trauma surgery. J Orthop Surg Res. 2025;20(1):931.

17. Moonen CTW, Kilroy JP, Klibanov AL. Focused Ultrasound: Noninvasive Image-Guided Therapy. Invest Radiol. 2025;60(3):205–219.

18. Li X, Liu Y. Focused ultrasound in modern medicine: bioengineering interfaces, molecular effects, and clinical breakthroughs. Front Bioeng Biotechnol. 2025;13:1610846.

19. He Y, Tan P, He M, et al. The primary treatment of prostate cancer with high-intensity focused ultrasound: A systematic review and meta-analysis. Medicine (Baltimore). 2020;99(41):e22610.

20. Tsai MC, Chang LT, Tam KW. Comparison of High-Intensity Focused Ultrasound and Conventional Surgery for Patients with Uterine Myomas: A Systematic Review and Meta-Analysis. J Minim Invasive Gynecol. 2021;28(10):1712–1724.

21. Metta V, Benamer HTS, Kapsas G, et al. MRI-Guided Focused Ultrasound in Parkinson’s Disease and Essential Tremor: Incisionless but Invasive. J Mov Disord. 2025;18(4):289–303.

22. Silva N, Green M, Roque D, Krishna V. The Use of Focused Ultrasound Ablation for Movement Disorders. Magn Reson Imaging Clin N Am. 2024;32(4):651–659.

23. Martínez-Fernández R, Máñez-Miró JU, Rodríguez-Rojas R, et al. Randomized Trial of Focused Ultrasound Subthalamotomy for Parkinson’s Disease. N Engl J Med. 2020;383(26):2501–2513.

24. Kamimura HAS, Conti A, Toschi N, Konofagou EE. Ultrasound neuromodulation: mechanisms and the potential of multimodal stimulation for neuronal function assessment. Front Phys. 2020;8:150.

25. Song M, Zhang M, He S, Li L, Hu H. Ultrasonic neuromodulation mediated by mechanosensitive ion channels: current and future. Front Neurosci. 2023;17:1232308.

26. Blackmore J, Shrivastava S, Sallet J, Butler CR, Cleveland RO. Ultrasound Neuromodulation: A Review of Results, Mechanisms and Safety. Ultrasound Med Biol. 2019;45(7):1509–1536.

27. Burgess MT, Apostolakis I, Konofagou EE. Power cavitation-guided blood-brain barrier opening with focused ultrasound and microbubbles. Phys Med Biol. 2018;63(6):065009.

28. Batts AJ, Ji R, Noel RL, et al. Ultrasound-mediated blood-brain barrier opening and safety evaluation. Theranostics. 2023;13(3):1180.

29. Colucci V, Strichartz G, Jolesz F, Vykhodtseva N, Hynynen K. Focused ultrasound effects on nerve action potential in vitro. Ultrasound Med Biol. 2009;35(10):1737–1747. Foley JL, Little JW, Vaezy S. Image-guided high-intensity focused ultrasound for conduction block of peripheral nerves. Ann Biomed Eng. 2007;35(1):109-119.

30. Foley JL, Little JW, Vaezy S. Effects of high-intensity focused ultrasound on nerve conduction. Muscle Nerve. 2008;37(2):241–250.

31. Foley JL, Little JW, Starr FL 3rd, Frantz C, Vaezy S. Image-guided HIFU neurolysis of peripheral nerves to treat spasticity and pain. Ultrasound Med Biol. 2004;30(9):1199–1207.

32. Lee YF, Lin CC, Cheng JS, Chen GS. Nerve conduction block in diabetic rats using high-intensity focused ultrasound for analgesic applications. Br J Anaesth. 2015;114(5):840–846.

33. Youn Y, Hellman A, Walling I, et al. High-intensity ultrasound treatment for vincristine-induced neuropathic pain. Neurosurgery. 2018;83(5):1068–1075.

34. Prabhala T, Hellman A, Walling I, et al. External focused ultrasound treatment for neuropathic pain induced by common peroneal nerve injury. Neurosci Lett. 2018;684:145–151.

35. Hellman A, Maietta T, Byraju K, et al. Effects of external low intensity focused ultrasound on electrophysiological changes in vivo in a rodent model of common peroneal nerve injury. Neuroscience. 2020;429:264–272.

36. Anderson TA, Pacharinsak C, Vilches-Moure J, Kantarci H, Zuchero JB, Butts-Pauly K, Yeomans D. Focused ultrasound-induced inhibition of peripheral nerve fibers in an animal model of acute pain. Reg Anesth Pain Med. 2023;48(9):462–470.

37. Anderson TA, Delgado J, Sun S, Behzadian N, Vilches-Moure J, Szlavik RB, Butts-Pauly K, Yeomans D. Dose-dependent effects of high intensity focused ultrasound on compound action potentials in an ex vivo rodent peripheral nerve model: comparison to local anesthetics.

38. Brennan TJ, Vandermeulen EP, Gebhart GF. Characterization of a rat model of incisional pain. Pain. 1996;64(3):493–502.

39. Hargreaves K, Dubner R, Brown F, et al. A new and sensitive method for measuring thermal nociception in cutaneous hyperalgesia. Pain 1988;32:77–88.

40. Randall LO, Selitto JJ. A method for measurement of analgesic activity on inflamed tissue. Arch Int Pharmacodyn Ther 1957;111:409–19.

41. Bertelli JA, Mira JC. The grasping test: a simple behavioral method for objective quantitative assessment of peripheral nerve regeneration in the rat. J Neurosci Methods 1995;58:151–5.

42. Davis M. Neurochemical modulation of sensory-motor reactivity: acoustic and tactile startle reflexes. Neurosci Biobehav Rev. 1980;4(2):241–263.

43. Legon W, Rowlands A, Opitz A, Sato TF, Tyler WJ. Pulsed ultrasound differentially stimulates somatosensory circuits in humans as indicated by EEG and fMRI. PLoS One. 2012;7:e51177.

44. Lele PP. Effects of focused ultrasonic radiation on peripheral nerve, with observations on local heating. Exp Neurol. 1963;8:47–83.

45. Jayathilake NJ, Phan TT, Kim J, Lee KP, Park JM. Modulating neuroplasticity for chronic pain relief: noninvasive neuromodulation as a promising approach. Exp Mol Med. 2025;57(3):501–514.

46. Young RR, Henneman E. Functional effects of focused ultrasound on mammalian nerves. Science. 1961;134(3489):1521–1522.

47. Xu D, Pollock M. Experimental nerve thermal injury. Brain. 1994;117(2):375–384.

48. Tsui PH, Wang SH, Huang CC. In vitro effects of ultrasound with different energies on the conduction properties of neural tissue. Ultrasonics. 2005;43(7):560–565.

49. Ilham SJ, Chen L, Guo T, Emadi S, Hoshino K, Feng B. In vitro single-unit recordings reveal increased peripheral nerve conduction velocity by focused pulsed ultrasound. Biomed Phys Eng Express. 2018;4(4):045004.

50. Lee YF, Lin CC, Cheng JS, Chen GS. Nerve conduction block in diabetic rats using high-intensity focused ultrasound for analgesic applications. Br J Anaesth. 2015 May;114(5):840–6. doi: 10.1093/bja/aeu443. Epub 2015 Jan 16. PMID: 25904608.

51. Moore JH, Gieck JH, Saliba EN, Perrin DH, Ball DW, McCue FC. The biophysical effects of ultrasound on median nerve distal latencies. Electromyogr Clin Neurophysiol. 2000;40(3):169–180.

52. Worthington A, Peng P, Rod K, Bril V, Tavakkoli J. Image-guided high intensity focused ultrasound system for large animal nerve ablation studies. IEEE Journal of Translational Engineering in Health and Medicine. 2016 Sep 7;4:1–6.

53. Dababou S, Marrocchio C, Scipione R, Erasmus HP, Ghanouni P, Anzidei M, et al. High-intensity focused ultrasound for pain management in patients with cancer. Radiographics. 2018;38:603–623.

54. di Biase L, Falato E, Caminiti ML, Pecoraro PM, Narducci F, Di Lazzaro V. Focused ultrasound (FUS) for chronic pain management: approved and potential applications. Neurol Res Int. 2021;2021:8438498.

55. Lee SA, Kamimura HAS, Konofagou EE. Displacement imaging during focused ultrasound median nerve modulation: a preliminary study in human pain sensation mitigation. IEEE Trans Ultrason Ferroelectr Freq Control. 2021;68:526–537.

