## Supplemental File for "Noninvasive Diagnostic Ultrasound-Guided Focused Ultrasound Enables Selective, Reversible Inhibition of Peripheral Nociceptive Fibers and Prevents Acute Pain"

version 3

### Supplementary Figures

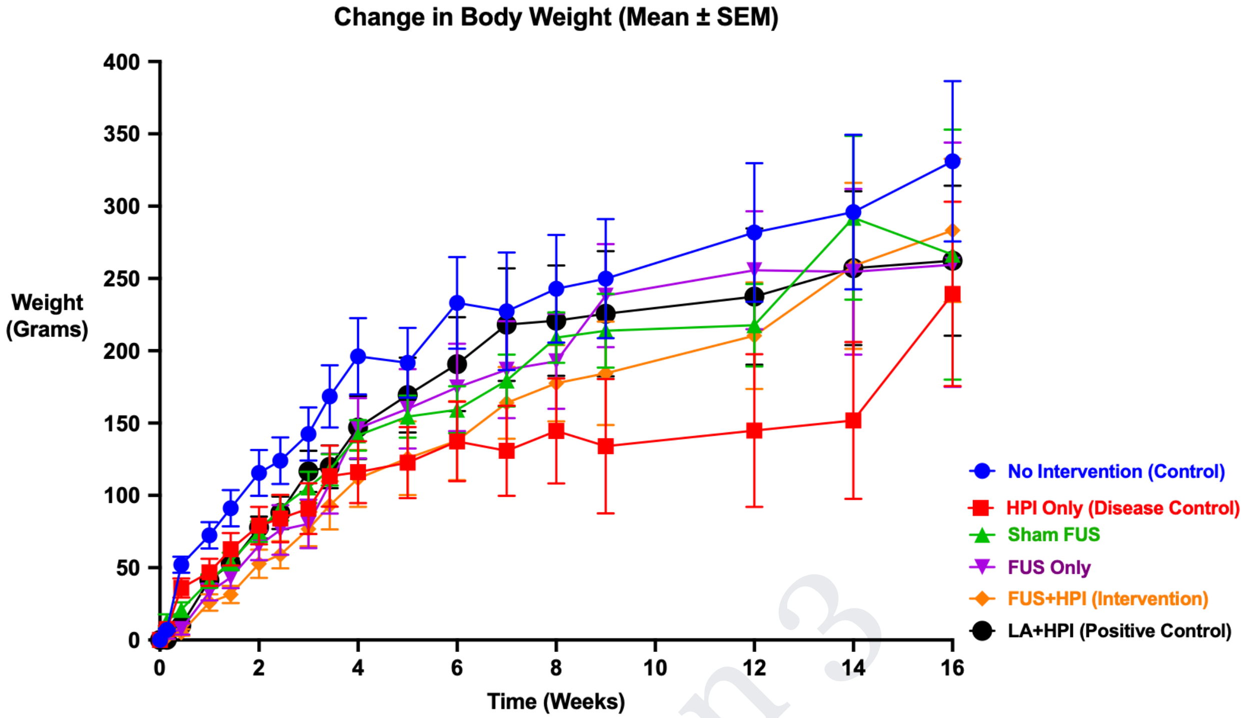

**Supplementary Figure 1 Body weight assessment.** Body weight measured across study arms over time. Data were represented as mean ± SEM. FUS, focused ultrasound; dUS, diagnostic ultrasound; HPI, hindpaw incision; LA, local anesthetic.

version 3

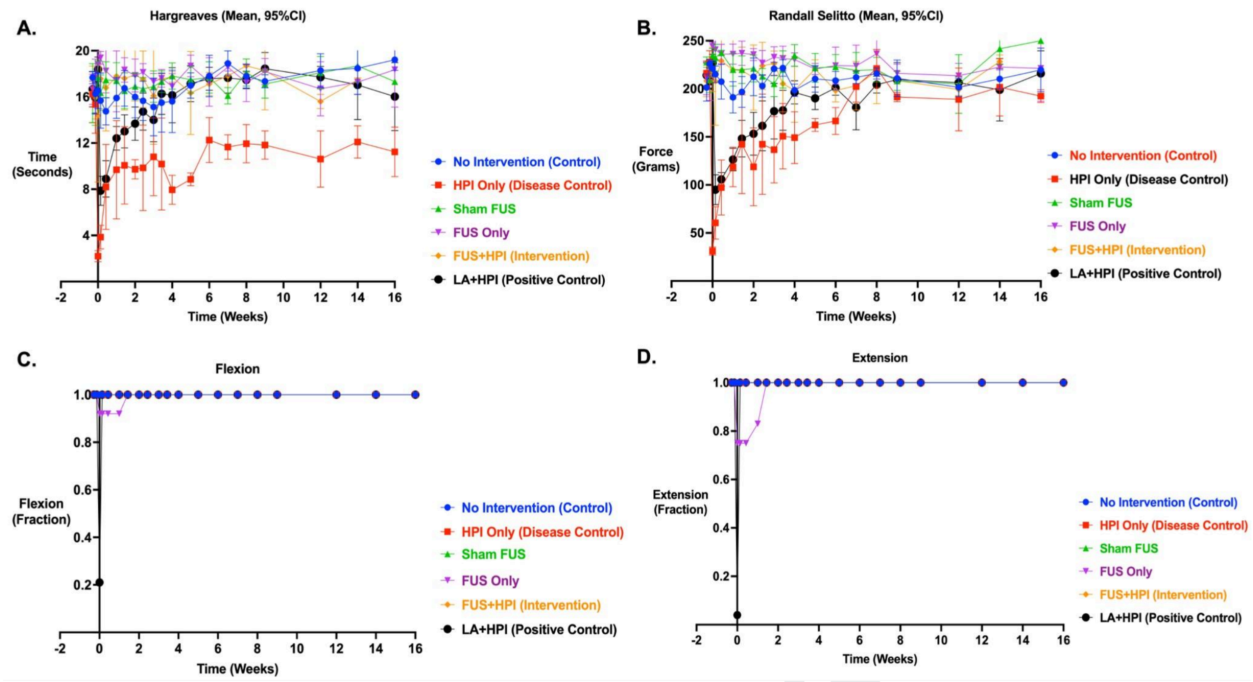

**Supplementary Figure 2 Behavioral assessment of nociceptive and sensorimotor function across study arms in male rats.** Behavioral test results from the intervention and control study arms. **(A)** Thermal withdrawal latency (modified Hargreaves test). **(B)** Mechanical withdrawal threshold (Randall–Selitto test). **(C)** Hindpaw (HP) flexion (grasp) reflex. **(D)** Hindpaw (HP) extension (startle) reflex. **Panels A and B:** Data are presented as mean ± 95% confidence interval (CI). Statistical comparisons were performed using a two-way mixed-effects ANOVA with fixed effects for study arm and time, followed by Bonferroni-corrected post hoc testing. Statistical comparisons were conducted relative to the HPI Only (Disease Control) study arm. Complete statistical results are provided in Supplementary Tables 4 and 9 for modified Hargreaves and Randall–Selitto testing, respectively. **Panels C and D:** Data were analyzed using contingency table analysis followed by Fisher's exact test, and behavioral responses are presented as fractional response values. Complete statistical results are provided in Supplementary Tables 14 and 17 for hindpaw flexion and extension testing, respectively. FUS, focused ultrasound; dUS, diagnostic ultrasound; HPI, hindpaw incision; LA, local anesthetic.

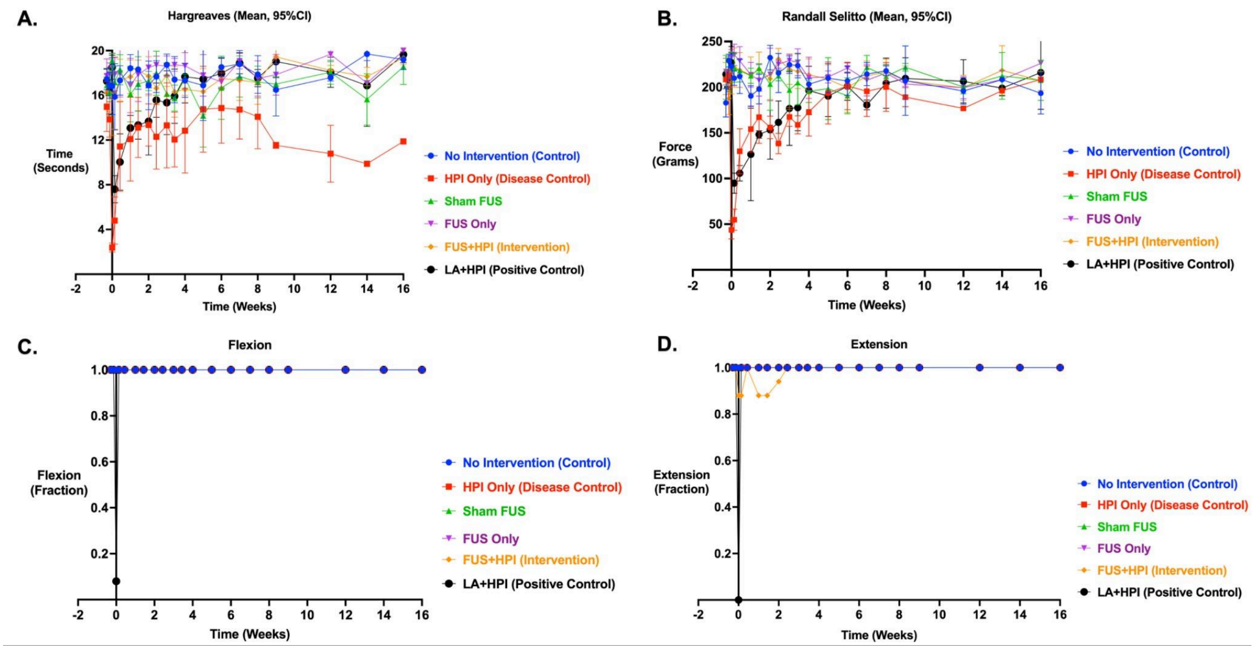

**Supplementary Figure 3 Behavioral assessment of nociceptive and sensorimotor function across study arms in female rats.** Behavioral test results from the intervention and control study arms. **(A)** Thermal withdrawal latency (modified Hargreaves test). **(B)** Mechanical withdrawal threshold (Randall–Selitto test). **(C)** Hindpaw (HP) flexion (grasp) reflex. **(D)** Hindpaw (HP) extension (startle) reflex. **Panels A and B:** Data are presented as mean ± 95% confidence interval (CI). Statistical comparisons were performed using a two-way mixed-effects ANOVA with fixed effects for study arm and time, followed by Bonferroni-corrected post hoc testing. Statistical comparisons were conducted relative to the HPI Only (Disease Control) study arm. Complete statistical results are provided in Supplementary Tables 6 and 11 for modified Hargreaves and Randall–Selitto testing, respectively. **Panels C and D:** Data were analyzed using contingency table analysis followed by Fisher's exact test, and behavioral responses are presented as fractional response values. Complete statistical results are provided in Supplementary Tables 15 and 18 for hindpaw flexion and extension testing, respectively. FUS, focused ultrasound; dUS, diagnostic ultrasound; HPI, hindpaw incision; LA, local anesthetic.

version 3

**Supplementary Tables**

| Time (Weeks) | Study Arms | Control | HPI Only | Sham FUS | FUS Only | FUS+HPI | LA+HPI |
|---|---|---|---|---|---|---|---|
| **-0.29** | **Number of Values (N)** | 12 | 12 | 12 | 12 | 12 | 12 |
| | **Mean** | 17.56 | 15.62 | 16.61 | 17.19 | 16.24 | 16.67 |
| | **Std. Deviation** | 0.92 | 2.2 | 2.26 | 1.79 | 2.23 | 2.69 |
| | **Std. Error of Mean** | 0.27 | 0.63 | 0.65 | 0.52 | 0.64 | 0.78 |
| | **Lower 95% CI of mean** | 16.98 | 14.22 | 15.18 | 16.05 | 14.82 | 14.96 |
| | **Upper 95% CI of mean** | 18.15 | 17.01 | 18.05 | 18.33 | 17.66 | 18.37 |
| **-0.14** | **Number of Values** | 12 | 12 | 12 | 12 | 12 | 12 |
| | **Mean** | 16.58 | 14.59 | 15.87 | 17.79 | 16.13 | 15.92 |
| | **Std. Deviation** | 1.8 | 2.61 | 2.41 | 1.59 | 2.19 | 2.35 |
| | **Std. Error of Mean** | 0.52 | 0.75 | 0.69 | 0.46 | 0.63 | 0.68 |
| | **Lower 95% CI of mean** | 15.44 | 12.93 | 14.35 | 16.77 | 14.74 | 14.43 |
| | **Upper 95% CI of mean** | 17.72 | 16.24 | 17.4 | 18.8 | 17.52 | 17.41 |
| **0** | **Number of Values** | 12 | 12 | 12 | 12 | 12 | 12 |
| | **Mean** | 16.69 | 2.29 | 18.4 | 18.62 | 15.33 | 18.38 |
| | **Std. Deviation** | 2.52 | 0.54 | 1.54 | 1.57 | 5.03 | 1.89 |

version 3

| | | | | | | | |
|---|---|---|---|---|---|---|---|
| | **Std. Error of Mean** | 0.73 | 0.16 | 0.44 | 0.45 | 1.45 | 0.55 |
| | **Lower 95% CI of mean** | 15.08 | 1.95 | 17.42 | 17.63 | 12.13 | 17.18 |
| | **Upper 95% CI of mean** | 18.29 | 2.64 | 19.37 | 19.62 | 18.52 | 19.58 |
| **0.14** | **Number of Values** | 12 | 12 | 12 | 12 | 12 | 12 |
| | **Mean** | 15.78 | 4.32 | 16.77 | 18.64 | 16.65 | 7.87 |
| | **Std. Deviation** | 3.2 | 2.04 | 1.83 | 1.55 | 4.03 | 2.25 |
| | **Std. Error of Mean** | 0.92 | 0.59 | 0.53 | 0.45 | 1.16 | 0.65 |
| | **Lower 95% CI of mean** | 13.74 | 3.03 | 15.61 | 17.65 | 14.09 | 6.44 |
| | **Upper 95% CI of mean** | 17.81 | 5.61 | 17.93 | 19.63 | 19.21 | 9.3 |
| **0.43** | **Number of Values** | 12 | 12 | 12 | 12 | 12 | 12 |
| | **Mean** | 16.04 | 9.81 | 17.88 | 18.11 | 17.17 | 8.9 |
| | **Std. Deviation** | 2.01 | 4.88 | 1.79 | 2.4 | 2.95 | 2.79 |
| | **Std. Error of Mean** | 0.58 | 1.41 | 0.52 | 0.69 | 0.85 | 0.8 |
| | **Lower 95% CI of mean** | 14.76 | 6.71 | 16.74 | 16.58 | 15.3 | 7.13 |
| | **Upper 95% CI of mean** | 17.32 | 12.91 | 19.02 | 19.63 | 19.04 | 10.67 |
| **1** | **Number of Values** | 12 | 12 | 12 | 12 | 12 | 12 |
| | **Mean** | 17.16 | 10.88 | 16.75 | 17.11 | 17.71 | 12.41 |

version 3

| | | | | | | | |
|---|---|---|---|---|---|---|---|
| | **Std. Deviation** | 2.24 | 4.94 | 2.57 | 3.03 | 2.21 | 1.8 |
| | **Std. Error of Mean** | 0.65 | 1.43 | 0.74 | 0.88 | 0.64 | 0.52 |
| | **Lower 95% CI of mean** | 15.74 | 7.75 | 15.12 | 15.19 | 16.3 | 11.27 |
| | **Upper 95% CI of mean** | 18.58 | 14.02 | 18.38 | 19.04 | 19.11 | 13.55 |
| **1.43** | **Number of Values** | 10 | 12 | 12 | 12 | 12 | 12 |
| | **Mean** | 17.37 | 11.6 | 16.78 | 18.1 | 18.02 | 13.01 |
| | **Std. Deviation** | 1.95 | 3.96 | 2.45 | 2.24 | 1.83 | 2.54 |
| | **Std. Error of Mean** | 0.62 | 1.14 | 0.71 | 0.65 | 0.53 | 0.73 |
| | **Lower 95% CI of mean** | 15.97 | 9.08 | 15.22 | 16.68 | 16.86 | 11.4 |
| | **Upper 95% CI of mean** | 18.76 | 14.11 | 18.33 | 19.52 | 19.18 | 14.62 |
| **2** | **Number of Values** | 10 | 10 | 12 | 10 | 12 | 10 |
| | **Mean** | 16.35 | 11.9 | 17.1 | 18.11 | 17.77 | 13.68 |
| | **Std. Deviation** | 2.17 | 2.61 | 1.23 | 1.96 | 2.33 | 2.32 |
| | **Std. Error of Mean** | 0.69 | 0.82 | 0.35 | 0.62 | 0.67 | 0.73 |
| | **Lower 95% CI of mean** | 14.8 | 10.04 | 16.32 | 16.71 | 16.29 | 12.02 |
| | **Upper 95% CI of mean** | 17.9 | 13.77 | 17.88 | 19.51 | 19.25 | 15.34 |
| **2.43** | **Number of Values** | 10 | 10 | 10 | 10 | 10 | 10 |

version 3

| | | | | | | | |
|---|---|---|---|---|---|---|---|
| | **Mean** | 16.47 | 11.31 | 17.21 | 18.36 | 16.89 | 14.72 |
| | **Std. Deviation** | 2.07 | 3.68 | 1.63 | 0.9 | 1.83 | 2.59 |
| | **Std. Error of Mean** | 0.66 | 1.17 | 0.51 | 0.29 | 0.58 | 0.82 |
| | **Lower 95% CI of mean** | 14.99 | 8.68 | 16.04 | 17.72 | 15.58 | 12.87 |
| | **Upper 95% CI of mean** | 17.95 | 13.95 | 18.37 | 19.01 | 18.2 | 16.57 |
| **3** | **Number of Values** | 10 | 10 | 10 | 10 | 10 | 10 |
| | **Mean** | 16.57 | 12.31 | 16.59 | 17.82 | 17.28 | 14.01 |
| | **Std. Deviation** | 3.04 | 4.27 | 2.1 | 1.09 | 2.21 | 2.36 |
| | **Std. Error of Mean** | 0.96 | 1.35 | 0.67 | 0.34 | 0.7 | 0.75 |
| | **Lower 95% CI of mean** | 14.39 | 9.26 | 15.08 | 17.04 | 15.7 | 12.32 |
| | **Upper 95% CI of mean** | 18.74 | 15.36 | 18.09 | 18.6 | 18.86 | 15.7 |
| **3.43** | **Number of Values** | 10 | 10 | 10 | 10 | 10 | 10 |
| | **Mean** | 16.26 | 11.3 | 16.79 | 18.06 | 16.68 | 16.27 |
| | **Std. Deviation** | 2.85 | 3.42 | 1.32 | 0.98 | 2.57 | 2.23 |
| | **Std. Error of Mean** | 0.9 | 1.08 | 0.44 | 0.31 | 0.81 | 0.71 |
| | **Lower 95% CI of mean** | 14.23 | 8.85 | 15.78 | 17.36 | 14.84 | 14.67 |
| | **Upper 95% CI of mean** | 18.3 | 13.75 | 17.81 | 18.76 | 18.52 | 17.87 |

version 3

| | | | | | | | |
|---|---|---|---|---|---|---|---|
| **4** | **Number of Values** | 8 | 8 | 8 | 9 | 8 | 10 |
| | **Mean** | 16.26 | 11 | 17.72 | 17.68 | 16.98 | 16.15 |
| | **Std. Deviation** | 2.6 | 4.17 | 1.01 | 1.47 | 0.81 | 2.99 |
| | **Std. Error of Mean** | 0.92 | 1.47 | 0.36 | 0.49 | 0.28 | 0.95 |
| | **Lower 95% CI of mean** | 14.09 | 7.52 | 16.88 | 16.55 | 16.3 | 14.01 |
| | **Upper 95% CI of mean** | 18.42 | 14.48 | 18.56 | 18.81 | 17.65 | 18.29 |
| **5** | **Number of Values** | 8 | 8 | 8 | 9 | 8 | 8 |
| | **Mean** | 16.96 | 12.54 | 16.24 | 18.29 | 16.34 | 17.15 |
| | **Std. Deviation** | 1.76 | 4.51 | 2.39 | 1.47 | 2.46 | 1.96 |
| | **Std. Error of Mean** | 0.62 | 1.59 | 0.84 | 0.49 | 0.87 | 0.69 |
| | **Lower 95% CI of mean** | 15.48 | 8.77 | 14.25 | 17.16 | 14.28 | 15.51 |
| | **Upper 95% CI of mean** | 18.43 | 16.3 | 18.23 | 19.41 | 18.39 | 18.79 |
| **6** | **Number of Values** | 8 | 8 | 8 | 8 | 8 | 8 |
| | **Mean** | 18.08 | 13.89 | 17.3 | 17.25 | 17.35 | 17.57 |
| | **Std. Deviation** | 1.65 | 3.16 | 1.85 | 1.44 | 0.82 | 1.23 |
| | **Std. Error of Mean** | 0.58 | 1.12 | 0.65 | 0.51 | 0.29 | 0.44 |
| | **Lower 95% CI of mean** | 16.7 | 11.25 | 15.76 | 16.05 | 16.67 | 16.54 |

version 3

| | | | | | | | |
|---|---|---|---|---|---|---|---|
| | **Upper 95% CI of mean** | 19.46 | 16.53 | 18.85 | 18.46 | 18.03 | 18.6 |
| **7** | **Number of Values** | 6 | 8 | 8 | 8 | 8 | 8 |
| | **Mean** | 18.85 | 13.58 | 16.7 | 18.82 | 17.53 | 17.65 |
| | **Std. Deviation** | 0.91 | 2.82 | 1.35 | 0.58 | 1.96 | 1.65 |
| | **Std. Error of Mean** | 0.37 | 1 | 0.48 | 0.21 | 0.69 | 0.58 |
| | **Lower 95% CI of mean** | 17.89 | 11.22 | 15.57 | 18.33 | 15.9 | 16.27 |
| | **Upper 95% CI of mean** | 19.81 | 15.94 | 17.82 | 19.3 | 19.17 | 19.03 |
| **8** | **Number of Values** | 6 | 8 | 8 | 8 | 8 | 6 |
| | **Mean** | 17.82 | 13.28 | 17.77 | 17.37 | 17.73 | 17.62 |
| | **Std. Deviation** | 1.1 | 2.81 | 1.31 | 1.53 | 1.88 | 0.77 |
| | **Std. Error of Mean** | 0.45 | 0.99 | 0.46 | 0.54 | 0.66 | 0.31 |
| | **Lower 95% CI of mean** | 16.66 | 10.93 | 16.67 | 16.09 | 16.16 | 16.81 |
| | **Upper 95% CI of mean** | 18.98 | 15.63 | 18.86 | 18.65 | 19.3 | 18.43 |
| **9** | **Number of Values** | 6 | 6 | 7 | 7 | 6 | 6 |
| | **Mean** | 16.93 | 11.68 | 17.07 | 18.04 | 19.05 | 18.89 |
| | **Std. Deviation** | 1.43 | 0.72 | 0.94 | 1.25 | 1.34 | 1.43 |
| | **Std. Error of Mean** | 0.58 | 0.29 | 0.36 | 0.47 | 0.55 | 0.58 |

version 3

| | | | | | | | |
|---|---|---|---|---|---|---|---|
| | **Lower 95% CI of mean** | 15.43 | 10.92 | 16.19 | 16.88 | 17.64 | 17.38 |
| | **Upper 95% CI of mean** | 18.43 | 12.44 | 17.94 | 19.2 | 20.46 | 20.39 |
| **12** | **Number of Values** | 5 | 6 | 6 | 6 | 5 | 6 |
| | **Mean** | 18.01 | 10.7 | 18.08 | 17.69 | 17.17 | 18.05 |
| | **Std. Deviation** | 1 | 1.98 | 0.99 | 2.41 | 1.48 | 0.95 |
| | **Std. Error of Mean** | 0.45 | 0.81 | 0.4 | 0.98 | 0.66 | 0.39 |
| | **Lower 95% CI of mean** | 16.77 | 8.62 | 17.05 | 15.17 | 15.33 | 17.05 |
| | **Upper 95% CI of mean** | 19.25 | 12.77 | 19.12 | 20.22 | 19.01 | 19.05 |
| **14** | **Number of Values** | 5 | 4 | 6 | 4 | 4 | 5 |
| | **Mean** | 18.97 | 11.55 | 17.16 | 17.22 | 17.58 | 17.71 |
| | **Std. Deviation** | 1.58 | 1.51 | 2.25 | 0.77 | 0.37 | 2.64 |
| | **Std. Error of Mean** | 0.7 | 0.75 | 0.92 | 0.38 | 0.19 | 1.18 |
| | **Lower 95% CI of mean** | 17.01 | 9.15 | 14.8 | 16 | 16.98 | 14.42 |
| | **Upper 95% CI of mean** | 20.93 | 13.94 | 19.53 | 18.44 | 18.17 | 20.99 |
| **16** | **Number of Values** | 3 | 4 | 4 | 3 | 2 | 4 |
| | **Mean** | 19.21 | 11.4 | 18.25 | 18.9 | 19.34 | 17.83 |
| | **Std. Deviation** | 0.8 | 1.58 | 1.3 | 1.91 | 0.42 | 2.12 |

version 3

| | | | | | | | |
|---|---|---|---|---|---|---|---|
| | **Std. Error of Mean** | 0.46 | 0.79 | 0.65 | 1.1 | 0.3 | 1.06 |
| | **Lower 95% CI of mean** | 17.23 | 8.89 | 16.18 | 14.17 | 15.53 | 14.45 |
| | **Upper 95% CI of mean** | 21.2 | 13.91 | 20.32 | 23.63 | 23.15 | 21.2 |

**Supplementary Table 1:** Descriptive statistical summary of the Hargreaves test (hindpaw thermal withdrawal latency) across all experimental groups, including No Intervention (Control; n = 12), HPI Only (Disease Control; n = 12), Sham FUS (n = 12), FUS Only (n = 12), FUS+HPI (Intervention; n = 12), and LA+HPI (Positive Control; n = 12) at each time point up to 16 weeks. Data are presented as mean, standard deviation (SD), standard error of the mean (SEM), and lower and upper 95% confidence intervals (CI). FUS = focused ultrasound; HPI = hindpaw incision; LA = local anesthetic.

version 3

| Time (Weeks) | Comparison | Summary | p Value |
|---|---|---|---|
| **-0.29** | **HPI Only vs Control** | ns | .343 |
| | HPI Only vs Sham FUS | ns | >.999 |
| | HPI Only vs FUS Only | ns | .949 |
| | HPI Only vs FUS+HPI | ns | >.999 |
| | HPI Only vs LA+HPI | ns | >.999 |
| **-0.14** | **HPI Only vs Control** | ns | .260 |
| | HPI Only vs Sham FUS | ns | >.999 |
| | HPI Only vs FUS Only | ** | .003 |
| | HPI Only vs FUS+HPI | ns | .940 |
| | HPI Only vs LA+HPI | ns | >.999 |
| **0** | **HPI Only vs Control** | *** | <.001 |

| | | | |
|---|---|---|---|
| | HPI Only vs Sham FUS | *** | <.001 |
| | HPI Only vs FUS Only | *** | <.001 |
| | HPI Only vs FUS+HPI | *** | <.001 |
| | HPI Only vs LA+HPI | *** | <.001 |
| **0.14** | **HPI Only vs Control** | *** | <.001 |
| | HPI Only vs Sham FUS | *** | <.001 |
| | HPI Only vs FUS Only | *** | <.001 |
| | HPI Only vs FUS+HPI | *** | <.001 |
| | HPI Only vs LA+HPI | ** | .009 |
| **0.43** | **HPI Only vs Control** | *** | <.001 |
| | HPI Only vs Sham FUS | *** | <.001 |
| | HPI Only vs FUS Only | *** | <.001 |
| | **HPI Only vs FUS+HPI** | *** | <.001 |
| | **HPI Only vs LA+HPI** | ns | >.999 |
| **1** | **HPI Only vs Control** | *** | <.001 |
| | **HPI Only vs Sham FUS** | *** | <.001 |
| | **HPI Only vs FUS Only** | *** | <.001 |
| | **HPI Only vs FUS+HPI** | *** | <.001 |
| | **HPI Only vs LA+HPI** | ns | >.999 |
| **1.43** | **HPI Only vs Control** | *** | <.001 |
| | **HPI Only vs Sham FUS** | *** | <.001 |

version 3

| | | | |
|---|---|---|---|
| | **HPI Only vs FUS Only** | *** | <.001 |
| | **HPI Only vs FUS+HPI** | *** | <.001 |
| | **HPI Only vs LA+HPI** | ns | >.999 |
| **2** | **HPI Only vs Control** | *** | <.001 |
| | **HPI Only vs Sham FUS** | *** | <.001 |
| | **HPI Only vs FUS Only** | *** | <.001 |
| | **HPI Only vs FUS+HPI** | *** | <.001 |
| | **HPI Only vs LA+HPI** | ns | .291 |
| **2.43** | **HPI Only vs Control** | *** | <.001 |
| | **HPI Only vs Sham FUS** | *** | <.001 |
| | **HPI Only vs FUS Only** | *** | <.001 |
| | **HPI Only vs FUS+HPI** | *** | <.001 |
| | **HPI Only vs LA+HPI** | ** | .002 |
| **3** | **HPI Only vs Control** | ** | .001 |
| | **HPI Only vs Sham FUS** | ** | .001 |
| | **HPI Only vs FUS Only** | *** | <.001 |
| | **HPI Only vs FUS+HPI** | *** | <.001 |
| | **HPI Only vs LA+HPI** | ns | .716 |
| **3.43** | **HPI Only vs Control** | *** | <.001 |
| | **HPI Only vs Sham FUS** | *** | <.001 |
| | **HPI Only vs FUS Only** | *** | <.001 |

version 3

| | | | |
|---|---|---|---|
| | **HPI Only vs FUS+HPI** | *** | <.001 |
| | **HPI Only vs LA+HPI** | *** | <.001 |
| **4** | **HPI Only vs Control** | ** | .002 |
| | **HPI Only vs Sham FUS** | *** | <.001 |
| | **HPI Only vs FUS Only** | *** | <.001 |
| | **HPI Only vs FUS+HPI** | *** | <.001 |
| | **HPI Only vs LA+HPI** | ** | .001 |
| **5** | **HPI Only vs Control** | *** | <.001 |
| | **HPI Only vs Sham FUS** | *** | <.001 |
| | **HPI Only vs FUS Only** | *** | <.001 |
| | **HPI Only vs FUS+HPI** | *** | <.001 |
| | **HPI Only vs LA+HPI** | *** | <.001 |
| **6** | **HPI Only vs Control** | *** | <.001 |
| | **HPI Only vs Sham FUS** | *** | <.001 |
| | **HPI Only vs FUS Only** | * | .010 |
| | **HPI Only vs FUS+HPI** | *** | <.001 |
| | **HPI Only vs LA+HPI** | *** | <.001 |
| **7** | **HPI Only vs Control** | *** | <.001 |
| | **HPI Only vs Sham FUS** | ** | .004 |
| | **HPI Only vs FUS Only** | *** | <.001 |
| | **HPI Only vs FUS+HPI** | *** | <.001 |

version 3

| | | | |
|---|---|---|---|
| | **HPI Only vs LA+HPI** | *** | <.001 |
| **8** | **HPI Only vs Control** | *** | <.001 |
| | **HPI Only vs Sham FUS** | *** | <.001 |
| | **HPI Only vs FUS Only** | ** | .007 |
| | **HPI Only vs FUS+HPI** | *** | <.001 |
| | **HPI Only vs LA+HPI** | *** | <.001 |
| **9** | **HPI Only vs Control** | *** | <.001 |
| | **HPI Only vs Sham FUS** | *** | <.001 |
| | **HPI Only vs FUS Only** | *** | <.001 |
| | **HPI Only vs FUS+HPI** | *** | <.001 |
| | **HPI Only vs LA+HPI** | *** | <.001 |
| **12** | **HPI Only vs Control** | *** | <.001 |
| | **HPI Only vs Sham FUS** | *** | <.001 |
| | **HPI Only vs FUS Only** | *** | <.001 |
| | **HPI Only vs FUS+HPI** | *** | <.001 |
| | **HPI Only vs LA+HPI** | *** | <.001 |
| **14** | **HPI Only vs Control** | ** | .001 |
| | **HPI Only vs Sham FUS** | * | .012 |
| | **HPI Only vs FUS Only** | * | .011 |
| | **HPI Only vs FUS+HPI** | ** | .003 |
| | **HPI Only vs LA+HPI** | ** | .007 |

version 3

| | | | |
|---|---|---|---|
| **16** | **HPI Only vs Control** | *** | <.001 |
| | **HPI Only vs Sham FUS** | *** | <.001 |
| | **HPI Only vs FUS Only** | *** | <.001 |
| | **HPI Only vs FUS+HPI** | ** | .001 |
| | **HPI Only vs LA+HPI** | * | .013 |

**Supplementary Table 2:** Summary of group-wise comparisons for the Hargreaves test (hindpaw thermal withdrawal latency) across all experimental groups, including No Intervention (Control; n = 12), HPI Only (Disease Control; n = 12), Sham FUS (n = 12), FUS Only (n = 12), FUS+HPI (Intervention; n = 12), and LA+HPI (Positive Control; n = 12) at each time point up to 16 weeks. Measurements were analyzed using a two-way mixed-effects ANOVA with fixed effects for treatment group and time, followed by Bonferroni-corrected post hoc multiple-comparison testing. Statistical significance is denoted as follows: ns ($p \geq 0.05$), * ($p < 0.05$), ** ($p < 0.01$), *** ($p < 0.001$).

| **Time (Weeks)** | **Study Arms** | **Control** | **HPI Only** | **Sham FUS** | **FUS Only** | **FUS+HPI** | **LA+HPI** |
|---|---|---|---|---|---|---|---|
| **-0.29** | **Number of Values (N)** | 6 | 6 | 6 | 6 | 4 | 6 |
| | Mean | 17.67 | 16.25 | 16.33 | 16.66 | 16.2 | 16.06 |
| | Std. Deviation | 0.67 | 1.4 | 3.19 | 1.64 | 2.52 | 3.64 |
| | Std. Error of Mean | 0.27 | 0.57 | 1.3 | 0.67 | 1.26 | 1.49 |
| | Lower 95% CI of mean | 16.97 | 14.77 | 12.98 | 14.94 | 12.18 | 12.25 |
| | Upper 95% CI of mean | 18.37 | 17.72 | 19.67 | 18.37 | 20.21 | 19.88 |
| **-0.14** | **Number of Values** | 6 | 6 | 6 | 6 | 4 | 6 |
| | Mean | 16.27 | 15.33 | 15.53 | 17.69 | 15.94 | 15.43 |

version 3

| | | | | | | | |
|---|---|---|---|---|---|---|---|
| | Std. Deviation | 2.2 | 3.12 | 2.49 | 1.98 | 1.9 | 2.19 |
| | Std. Error of Mean | 0.9 | 1.27 | 1.02 | 0.81 | 0.95 | 0.89 |
| | Lower 95% CI of mean | 13.96 | 12.06 | 12.92 | 15.61 | 12.93 | 13.13 |
| | Upper 95% CI of mean | 18.58 | 18.60 | 18.14 | 19.76 | 18.96 | 17.72 |
| **0** | **Number of Values** | 6 | 6 | 6 | 6 | 4 | 6 |
| | Mean | 16.54 | 2.2 | 17.67 | 18.78 | 14.4 | 18.26 |
| | Std. Deviation | 1.95 | 0.6 | 1.82 | 1.63 | 5.9 | 2.45 |
| | Std. Error of Mean | 0.8 | 0.25 | 0.74 | 0.66 | 2.95 | 1 |
| | Lower 95% CI of mean | 14.49 | 1.565 | 15.76 | 17.07 | 5.015 | 15.69 |
| | Upper 95% CI of mean | 18.58 | 2.831 | 19.58 | 20.49 | 23.79 | 20.83 |
| **0.14** | **Number of Values** | 6 | 6 | 6 | 6 | 4 | 6 |
| | Mean | 15.68 | 3.85 | 16.38 | 19.39 | 17.57 | 8.15 |
| | Std. Deviation | 2.98 | 1.29 | 1.91 | 0.69 | 2.65 | 2.94 |
| | Std. Error of Mean | 1.22 | 0.53 | 0.78 | 0.28 | 1.33 | 1.2 |
| | Lower 95% CI of mean | 12.55 | 2.496 | 14.37 | 18.66 | 13.35 | 5.058 |
| | **Upper 95% CI of mean** | 18.81 | 5.200 | 18.38 | 20.11 | 21.79 | 11.24 |
| **0.43** | **Number of Values** | 6 | 6 | 6 | 6 | 4 | 6 |

version 3

| | | | | | | | |
|---|---|---|---|---|---|---|---|
| | **Mean** | 14.75 | 8.2 | 17.47 | 18.25 | 16.79 | 7.75 |
| | **Std. Deviation** | 1.48 | 4.6 | 1.96 | 2.37 | 3.82 | 1.99 |
| | **Std. Error of Mean** | 0.6 | 1.88 | 0.8 | 0.97 | 1.91 | 0.81 |
| | **Lower 95% CI of mean** | 13.19 | 3.37 | 15.42 | 15.77 | 10.7 | 5.66 |
| | **Upper 95% CI of mean** | 16.3 | 13.03 | 19.52 | 20.74 | 22.87 | 9.84 |
| **1** | **Number of Values** | 6 | 6 | 6 | 6 | 4 | 6 |
| | **Mean** | 15.91 | 9.69 | 17.4 | 17.35 | 17.79 | 11.74 |
| | **Std. Deviation** | 2.22 | 5.32 | 1.97 | 3.3 | 2.64 | 2.02 |
| | **Std. Error of Mean** | 0.9 | 2.17 | 0.8 | 1.35 | 1.32 | 0.82 |
| | **Lower 95% CI of mean** | 13.58 | 4.11 | 15.33 | 13.88 | 13.59 | 9.63 |
| | **Upper 95% CI of mean** | 18.24 | 15.27 | 19.46 | 20.81 | 21.99 | 13.86 |
| **1.43** | **Number of Values** | 6 | 6 | 6 | 6 | 4 | 6 |
| | **Mean** | 16.75 | 10.07 | 16.53 | 18.33 | 17.6 | 12.68 |
| | **Std. Deviation** | 1.94 | 4.19 | 1.94 | 1.32 | 2.79 | 3.38 |
| | **Std. Error of Mean** | 0.79 | 1.71 | 0.79 | 0.54 | 1.4 | 1.38 |
| | **Lower 95% CI of mean** | 14.71 | 5.68 | 14.49 | 16.94 | 13.16 | 9.13 |
| | **Upper 95% CI of mean** | 18.78 | 14.47 | 18.57 | 19.72 | 22.04 | 16.23 |

version 3

| | | | | | | | |
|---|---|---|---|---|---|---|---|
| **2** | **Number of Values** | 6 | 4 | 6 | 6 | 4 | 6 |
| | **Mean** | 15.99 | 9.74 | 16.91 | 17.86 | 17.9 | 13.69 |
| | **Std. Deviation** | 2.26 | 0.85 | 1.44 | 1.78 | 2.29 | 2 |
| | **Std. Error of Mean** | 0.92 | 0.42 | 0.59 | 0.73 | 1.15 | 0.82 |
| | **Lower 95% CI of mean** | 13.61 | 8.39 | 15.4 | 15.99 | 14.25 | 11.59 |
| | **Upper 95% CI of mean** | 18.36 | 11.08 | 18.42 | 19.73 | 21.54 | 15.8 |
| **2.43** | **Number of Values** | 6 | 4 | 6 | 6 | 3 | 6 |
| | **Mean** | 15.66 | 9.85 | 16.69 | 18.12 | 17.51 | 14.16 |
| | **Std. Deviation** | 2.13 | 3.77 | 1.23 | 0.95 | 2.17 | 3.11 |
| | **Std. Error of Mean** | 0.87 | 1.89 | 0.5 | 0.39 | 1.25 | 1.27 |
| | **Lower 95% CI of mean** | 13.43 | 3.84 | 15.4 | 17.12 | 12.12 | 10.89 |
| | **Upper 95% CI of mean** | 17.89 | 15.85 | 17.98 | 19.12 | 22.91 | 17.42 |
| **3** | **Number of Values** | 6 | 4 | 6 | 6 | 3 | 6 |
| | **Mean** | 15.12 | 10.81 | 16.89 | 17.31 | 16.09 | 13.12 |
| | **Std. Deviation** | 3.12 | 3.44 | 1.71 | 0.9 | 3.71 | 1.61 |
| | **Std. Error of Mean** | 1.27 | 1.72 | 0.7 | 0.37 | 2.14 | 0.66 |
| | **Lower 95% CI of mean** | 11.84 | 5.34 | 15.1 | 16.37 | 6.88 | 11.43 |

version 3

| | | | | | | | |
|---|---|---|---|---|---|---|---|
| | **Upper 95% CI of mean** | 18.39 | 16.28 | 18.68 | 18.25 | 25.3 | 14.81 |
| **3.43** | **Number of Values** | 6 | 4 | 6 | 6 | 3 | 6 |
| | **Mean** | 15.5 | 10.19 | 17.32 | 17.62 | 17.54 | 16.5 |
| | **Std. Deviation** | 3.19 | 4.07 | 1.17 | 0.96 | 0.4 | 2.26 |
| | **Std. Error of Mean** | 1.3 | 2.03 | 0.48 | 0.39 | 0.23 | 0.92 |
| | **Lower 95% CI of mean** | 12.16 | 3.72 | 16.09 | 16.61 | 16.54 | 14.13 |
| | **Upper 95% CI of mean** | 18.85 | 16.66 | 18.55 | 18.62 | 18.54 | 18.88 |
| **4** | **Number of Values** | 5 | 3 | 5 | 5 | 3 | 6 |
| | **Mean** | 15.61 | 7.95 | 17.81 | 16.91 | 17.73 | 15.14 |
| | **Std. Deviation** | 3.08 | 1.1 | 1.31 | 1.25 | 0.41 | 3.1 |
| | **Std. Error of Mean** | 1.38 | 0.64 | 0.58 | 0.56 | 0.24 | 1.26 |
| | **Lower 95% CI of mean** | 11.79 | 5.22 | 16.19 | 15.36 | 16.71 | 11.89 |
| | **Upper 95% CI of mean** | 19.44 | 10.69 | 19.43 | 18.46 | 18.75 | 18.38 |
| **5** | **Number of Values** | 5 | 3 | 5 | 5 | 3 | 5 |
| | **Mean** | 16.99 | 8.86 | 17.48 | 18.68 | 16.34 | 16.96 |
| | **Std. Deviation** | 1.26 | 0.5 | 1.31 | 1.08 | 3.17 | 2 |
| | **Std. Error of Mean** | 0.56 | 0.29 | 0.59 | 0.48 | 1.83 | 0.89 |

version 3

| | | | | | | | |
|---|---|---|---|---|---|---|---|
| | **Lower 95% CI of mean** | 15.43 | 7.62 | 15.86 | 17.35 | 8.46 | 14.49 |
| | **Upper 95% CI of mean** | 18.55 | 10.09 | 19.11 | 20.02 | 24.22 | 19.44 |
| **6** | **Number of Values** | 5 | 3 | 5 | 4 | 3 | 5 |
| | **Mean** | 17.81 | 12.26 | 17.68 | 17.29 | 17.11 | 17.34 |
| | **Std. Deviation** | 2.06 | 1.72 | 1.55 | 2.14 | 0.87 | 1.32 |
| | **Std. Error of Mean** | 0.92 | 1 | 0.69 | 1.07 | 0.5 | 0.59 |
| | **Lower 95% CI of mean** | 15.26 | 7.98 | 15.76 | 13.88 | 14.95 | 15.7 |
| | **Upper 95% CI of mean** | 20.37 | 16.55 | 19.61 | 20.7 | 19.28 | 18.99 |
| **7** | **Number of Values** | 3 | 3 | 5 | 4 | 3 | 5 |
| | **Mean** | 18.9 | 11.66 | 16.1 | 18.58 | 17.82 | 16.92 |
| | **Std. Deviation** | 0.97 | 0.94 | 0.81 | 0.36 | 2.15 | 1.62 |
| | **Std. Error of Mean** | 0.56 | 0.54 | 0.36 | 0.18 | 1.24 | 0.72 |
| | **Lower 95% CI of mean** | 16.49 | 9.33 | 15.1 | 18 | 12.48 | 14.9 |
| | **Upper 95% CI of mean** | 21.3 | 14 | 17.1 | 19.16 | 23.15 | 18.93 |
| **8** | **Number of Values** | 3 | 3 | 5 | 4 | 3 | 3 |
| | **Mean** | 17.78 | 11.94 | 18.11 | 17.23 | 18.71 | 17.7 |
| | **Std. Deviation** | 0.9 | 1.46 | 1.35 | 1.67 | 0.52 | 1 |

version 3

| | | | | | | | |
|---|---|---|---|---|---|---|---|
| | **Std. Error of Mean** | 0.52 | 0.85 | 0.6 | 0.83 | 0.3 | 0.58 |
| | **Lower 95% CI of mean** | 15.55 | 8.3 | 16.44 | 14.58 | 17.42 | 15.21 |
| | **Upper 95% CI of mean** | 20.01 | 15.58 | 19.78 | 19.88 | 19.99 | 20.19 |
| **9** | **Number of Values** | 3 | 3 | 4 | 4 | 2 | 3 |
| | **Mean** | 17.36 | 11.83 | 17.11 | 18.19 | 18.27 | 18.74 |
| | **Std. Deviation** | 0.5 | 1.08 | 1.24 | 1.16 | 2.45 | 1.85 |
| | **Std. Error of Mean** | 0.29 | 0.62 | 0.62 | 0.58 | 1.73 | 1.07 |
| | **Lower 95% CI of mean** | 16.13 | 9.15 | 15.13 | 16.34 | -3.71 | 14.13 |
| | **Upper 95% CI of mean** | 18.59 | 14.52 | 19.08 | 20.04 | 40.25 | 23.34 |
| **12** | **Number of Values** | 3 | 3 | 3 | 4 | 2 | 3 |
| | **Mean** | 18.29 | 10.62 | 18.09 | 16.71 | 15.6 | 18.03 |
| | **Std. Deviation** | 1.3 | 2.16 | 1.25 | 2.38 | 0.44 | 0.92 |
| | **Std. Error of Mean** | 0.75 | 1.25 | 0.72 | 1.19 | 0.31 | 0.53 |
| | **Lower 95% CI of mean** | 15.07 | 5.26 | 14.99 | 12.91 | 11.66 | 15.75 |
| | **Upper 95% CI of mean** | 21.5 | 15.97 | 21.18 | 20.5 | 19.54 | 20.31 |
| **14** | **Number of Values** | 3 | 3 | 3 | 3 | 2 | 2 |
| | **Mean** | 18.48 | 12.11 | 18.69 | 17.27 | 17.47 | 18.97 |

version 3

| | | | | | | | |
|---|---|---|---|---|---|---|---|
| | **Std. Deviation** | 2 | 1.23 | 1.26 | 0.93 | 0.13 | 1.35 |
| | **Std. Error of Mean** | 1.15 | 0.71 | 0.73 | 0.54 | 0.09 | 0.96 |
| | **Lower 95% CI of mean** | 13.52 | 9.04 | 15.55 | 14.95 | 16.33 | 6.83 |
| | **Upper 95% CI of mean** | 23.45 | 15.17 | 21.83 | 19.58 | 18.61 | 31.1 |
| **16** | **Number of Values** | 3 | 3 | 1 | 2 | 0 | 2 |
| | **Mean** | 19.21 | 11.24 | 17.32 | 18.35 | | 16.02 |
| | **Std. Deviation** | 0.8 | 1.9 | 0 | 2.33 | | 0.38 |
| | **Std. Error of Mean** | 0.46 | 1.09 | 0 | 1.65 | | 0.27 |
| | **Lower 95% CI of mean** | 17.23 | 6.53 | | -2.62 | | 12.59 |
| | **Upper 95% CI of mean** | 21.2 | 15.95 | | 39.32 | | 19.45 |

version 3

**Supplementary Table 3:** Descriptive statistical summary of modified Hargreaves testing (thermal withdrawal latency) across all experimental groups in male rats, including No Intervention (Control; n = 6), HPI Only (Disease Control; n = 6), Sham FUS (n = 6), FUS Only (n = 6), FUS+HPI (Intervention; n = 5), and LA+HPI (Positive Control; n = 6) at each time point up to 16 weeks. Data are presented as mean, standard deviation (SD), standard error of the mean (SEM), and lower and upper 95% confidence intervals (CI). FUS = focused ultrasound; HPI = hindpaw incision; LA = local anesthetic.

| Time (Weeks) | Comparison | Summary | p Value |
|---|---|---|---|
| **-0.29** | **HPI Only vs Control** | ns | >.999 |
| | **HPI Only vs Sham FUS** | ns | >.999 |
| | **HPI Only vs FUS Only** | ns | >.999 |

| | | | |
|---|---|---|---|
| | **HPI Only vs FUS+HPI** | ns | >.999 |
| | **HPI Only vs LA+HPI** | ns | >.999 |
| **-0.14** | **HPI Only vs Control** | ns | >.999 |
| | **HPI Only vs Sham FUS** | ns | >.999 |
| | **HPI Only vs FUS Only** | ns | >.999 |
| | **HPI Only vs FUS+HPI** | ns | >.999 |
| | **HPI Only vs LA+HPI** | ns | >.999 |
| **0** | **HPI Only vs Control** | *** | <.001 |
| | **HPI Only vs Sham FUS** | *** | <.001 |
| | **HPI Only vs FUS Only** | *** | <.001 |
| | **HPI Only vs FUS+HPI** | *** | <.001 |
| | **HPI Only vs LA+HPI** | *** | <.001 |
| **0.14** | **HPI Only vs Control** | *** | <.001 |
| | **HPI Only vs Sham FUS** | *** | <.001 |
| | **HPI Only vs FUS Only** | *** | <.001 |
| | **HPI Only vs FUS+HPI** | *** | <.001 |
| | **HPI Only vs LA+HPI** | * | .048 |
| **0.43** | **HPI Only vs Control** | ** | .003 |
| | **HPI Only vs Sham FUS** | *** | <.001 |
| | **HPI Only vs FUS Only** | *** | <.001 |
| | **HPI Only vs FUS+HPI** | *** | <.001 |

version 3

| | | | |
|---|---|---|---|
| | **HPI Only vs LA+HPI** | ns | >.999 |
| **1** | **HPI Only vs Control** | * | .026 |
| | **HPI Only vs Sham FUS** | ** | .003 |
| | **HPI Only vs FUS Only** | ** | .003 |
| | **HPI Only vs FUS+HPI** | ** | .003 |
| | **HPI Only vs LA+HPI** | ns | >.999 |
| **1.43** | **HPI Only vs Control** | ** | .009 |
| | **HPI Only vs Sham FUS** | * | .012 |
| | **HPI Only vs FUS Only** | *** | <.001 |
| | **HPI Only vs FUS+HPI** | ** | .006 |
| | **HPI Only vs LA+HPI** | ns | >.999 |
| **2** | **HPI Only vs Control** | ** | .001 |
| | **HPI Only vs Sham FUS** | *** | <.001 |
| | **HPI Only vs FUS Only** | *** | <.001 |
| | **HPI Only vs FUS+HPI** | *** | <.001 |
| | **HPI Only vs LA+HPI** | ns | .104 |
| **2.43** | **HPI Only vs Control** | * | .029 |
| | **HPI Only vs Sham FUS** | ** | .007 |
| | **HPI Only vs FUS Only** | *** | <.001 |
| | **HPI Only vs FUS+HPI** | * | .011 |
| | **HPI Only vs LA+HPI** | ns | .220 |

version 3

| | | | |
|---|---|---|---|
| **3** | **HPI Only vs Control** | ns | .457 |
| | **HPI Only vs Sham FUS** | * | .044 |
| | **HPI Only vs FUS Only** | * | .025 |
| | **HPI Only vs FUS+HPI** | ns | .148 |
| | **HPI Only vs LA+HPI** | ns | >.999 |
| **3.43** | **HPI Only vs Control** | ** | .007 |
| | **HPI Only vs Sham FUS** | *** | <.001 |
| | **HPI Only vs FUS Only** | *** | <.001 |
| | **HPI Only vs FUS+HPI** | ** | .001 |
| | **HPI Only vs LA+HPI** | ** | .001 |
| **4** | **HPI Only vs Control** | ** | .006 |
| | **HPI Only vs Sham FUS** | *** | <.001 |
| | **HPI Only vs FUS Only** | ** | .002 |
| | **HPI Only vs FUS+HPI** | *** | <.001 |
| | **HPI Only vs LA+HPI** | ** | .009 |
| **5** | **HPI Only vs Control** | *** | <.001 |
| | **HPI Only vs Sham FUS** | *** | <.001 |
| | **HPI Only vs FUS Only** | *** | <.001 |
| | **HPI Only vs FUS+HPI** | *** | <.001 |
| | **HPI Only vs LA+HPI** | *** | <.001 |
| **6** | **HPI Only vs Control** | * | .034 |

version 3

| | | | |
|---|---|---|---|
| | **HPI Only vs Sham FUS** | * | .017 |
| | **HPI Only vs FUS Only** | ns | .126 |
| | **HPI Only vs FUS+HPI** | ns | .065 |
| | **HPI Only vs LA+HPI** | * | .028 |
| **7** | **HPI Only vs Control** | *** | <.001 |
| | **HPI Only vs Sham FUS** | * | .031 |
| | **HPI Only vs FUS Only** | ** | .001 |
| | **HPI Only vs FUS+HPI** | ** | .002 |
| | **HPI Only vs LA+HPI** | ** | .008 |
| **8** | **HPI Only vs Control** | ** | .003 |
| | **HPI Only vs Sham FUS** | *** | <.001 |
| | **HPI Only vs FUS Only** | ** | .007 |
| | **HPI Only vs FUS+HPI** | *** | <.001 |
| | **HPI Only vs LA+HPI** | ** | .003 |
| **9** | **HPI Only vs Control** | * | .027 |
| | **HPI Only vs Sham FUS** | ns | .059 |
| | **HPI Only vs FUS Only** | * | .042 |
| | **HPI Only vs FUS+HPI** | * | .021 |
| | **HPI Only vs LA+HPI** | ** | .007 |
| **12** | **HPI Only vs Control** | ** | .002 |
| | **HPI Only vs Sham FUS** | ** | .003 |

version 3

| | | | |
|---|---|---|---|
| | **HPI Only vs FUS Only** | ** | .004 |
| | **HPI Only vs FUS+HPI** | ns | .061 |
| | **HPI Only vs LA+HPI** | ** | .003 |
| **14** | **HPI Only vs Control** | * | .015 |
| | **HPI Only vs Sham FUS** | * | .013 |
| | **HPI Only vs FUS Only** | ns | .157 |
| | **HPI Only vs FUS+HPI** | ns | .115 |
| | **HPI Only vs LA+HPI** | * | .030 |
| **16** | **HPI Only vs Control** | * | .024 |
| | **HPI Only vs Sham FUS** | ns | .069 |
| | **HPI Only vs FUS Only** | ns | .267 |
| | **HPI Only vs FUS+HPI** | ns | >.999 |
| | **HPI Only vs LA+HPI** | ns | >.999 |

version 3

**Supplementary Table 4:** Summary of group-wise comparisons for modified Hargreaves testing (thermal withdrawal latency) across all experimental groups in male rats, including No Intervention (Control; n = 6), HPI Only (Disease Control; n = 6), Sham FUS (n = 6), FUS Only (n = 6), FUS+HPI (Intervention; n = 5), and LA+HPI (Positive Control; n = 6) at each time point up to 16 weeks. Measurements were analyzed using a two-way mixed-effects ANOVA with fixed effects for treatment group and time, followed by Bonferroni-corrected post hoc multiple-comparison testing. Statistical significance is denoted as follows: ns ($p \geq 0.05$), * ($p < 0.05$), ** ($p < 0.01$), *** ($p < 0.001$).

| **Time (Weeks)** | **Study Arms** | **Control** | **HPI Only** | **Sham FUS** | **FUS Only** | **FUS+HPI** | **LA+HPI** |
|---|---|---|---|---|---|---|---|
| **-0.29** | **Number of Values (N)** | 6 | 6 | 6 | 6 | 8 | 6 |
| | **Mean** | 17.46 | 14.99 | 16.9 | 17.73 | 16.27 | 17.27 |

| | | | | | | | |
|---|---|---|---|---|---|---|---|
| | **Std. Deviation** | 1.18 | 2.78 | 0.93 | 1.92 | 2.26 | 1.34 |
| | **Std. Error of Mean** | 0.48 | 1.14 | 0.38 | 0.78 | 0.8 | 0.55 |
| | **Lower 95% CI of mean** | 16.21 | 12.07 | 15.92 | 15.72 | 14.38 | 15.86 |
| | **Upper 95% CI of mean** | 18.7 | 17.9 | 17.88 | 19.74 | 18.15 | 18.68 |
| **-0.14** | **Number of Values** | 6 | 6 | 6 | 6 | 8 | 6 |
| | **Mean** | 16.88 | 13.84 | 16.22 | 17.89 | 16.22 | 16.42 |
| | **Std. Deviation** | 1.42 | 1.98 | 2.5 | 1.28 | 2.44 | 2.6 |
| | **Std. Error of Mean** | 0.58 | 0.81 | 1.02 | 0.52 | 0.86 | 1.06 |
| | **Lower 95% CI of mean** | 15.39 | 11.76 | 13.6 | 16.55 | 14.18 | 13.69 |
| | **Upper 95% CI of mean** | 18.37 | 15.92 | 18.84 | 19.23 | 18.26 | 19.14 |
| **0** | **Number of Values** | 6 | 6 | 6 | 6 | 8 | 6 |
| | **Mean** | 16.83 | 2.39 | 19.13 | 18.47 | 15.79 | 18.51 |
| | **Std. Deviation** | 3.18 | 0.5 | 0.78 | 1.64 | 4.91 | 1.35 |
| | **Std. Error of Mean** | 1.3 | 0.21 | 0.32 | 0.67 | 1.74 | 0.55 |
| | **Lower 95% CI of mean** | 13.49 | 1.86 | 18.31 | 16.75 | 11.68 | 17.09 |
| | **Upper 95% CI of mean** | 20.17 | 2.92 | 19.94 | 20.19 | 19.89 | 19.93 |

version 3

| | | | | | | | |
|---|---|---|---|---|---|---|---|
| **0.14** | **Number of Values** | 6 | 6 | 6 | 6 | 8 | 6 |
| | **Mean** | 15.88 | 4.79 | 17.17 | 17.89 | 16.2 | 7.6 |
| | **Std. Deviation** | 3.7 | 2.63 | 1.83 | 1.87 | 4.67 | 1.52 |
| | **Std. Error of Mean** | 1.51 | 1.07 | 0.75 | 0.76 | 1.65 | 0.62 |
| | **Lower 95% CI of mean** | 11.99 | 2.03 | 15.25 | 15.93 | 12.3 | 6.01 |
| | **Upper 95% CI of mean** | 19.76 | 7.55 | 19.09 | 19.86 | 20.1 | 9.19 |
| **0.43** | **Number of Values** | 6 | 6 | 6 | 6 | 8 | 6 |
| | **Mean** | 17.34 | 11.42 | 18.29 | 17.96 | 17.36 | 10.04 |
| | **Std. Deviation** | 1.65 | 5 | 1.68 | 2.65 | 2.7 | 3.15 |
| | **Std. Error of Mean** | 0.67 | 2.04 | 0.69 | 1.08 | 0.95 | 1.29 |
| | **Lower 95% CI of mean** | 15.61 | 6.17 | 16.53 | 15.18 | 15.11 | 6.74 |
| | **Upper 95% CI of mean** | 19.06 | 16.67 | 20.05 | 20.75 | 19.62 | 13.35 |
| **1** | **Number of Values** | 6 | 6 | 6 | 6 | 8 | 6 |
| | **Mean** | 18.42 | 12.08 | 16.1 | 16.88 | 17.66 | 13.08 |
| | **Std. Deviation** | 1.52 | 4.69 | 3.1 | 3.04 | 2.17 | 1.39 |
| | **Std. Error of Mean** | 0.62 | 1.91 | 1.27 | 1.24 | 0.77 | 0.57 |

version 3

| | | | | | | | |
|---|---|---|---|---|---|---|---|
| | **Lower 95% CI of mean** | 16.82 | 7.16 | 12.85 | 13.69 | 15.85 | 11.62 |
| | **Upper 95% CI of mean** | 20.01 | 17 | 19.36 | 20.06 | 19.47 | 14.54 |
| **1.43** | **Number of Values** | 4 | 6 | 6 | 6 | 8 | 6 |
| | **Mean** | 18.31 | 13.12 | 17.02 | 17.87 | 18.23 | 13.34 |
| | **Std. Deviation** | 1.79 | 3.36 | 3.05 | 3.02 | 1.33 | 1.58 |
| | **Std. Error of Mean** | 0.9 | 1.37 | 1.25 | 1.23 | 0.47 | 0.64 |
| | **Lower 95% CI of mean** | 15.45 | 9.59 | 13.81 | 14.7 | 17.12 | 11.68 |
| | **Upper 95% CI of mean** | 21.16 | 16.65 | 20.22 | 21.04 | 19.34 | 14.99 |
| **2** | **Number of Values** | 4 | 6 | 6 | 4 | 8 | 4 |
| | **Mean** | 16.89 | 13.35 | 17.29 | 18.48 | 17.71 | 13.67 |
| | **Std. Deviation** | 2.22 | 2.35 | 1.07 | 2.42 | 2.51 | 3.07 |
| | **Std. Error of Mean** | 1.11 | 0.96 | 0.44 | 1.21 | 0.89 | 1.54 |
| | **Lower 95% CI of mean** | 13.36 | 10.88 | 16.16 | 14.63 | 15.61 | 8.78 |
| | **Upper 95% CI of mean** | 20.41 | 15.82 | 18.42 | 22.34 | 19.8 | 18.56 |
| **2.43** | **Number of Values** | 4 | 6 | 4 | 4 | 7 | 4 |
| | **Mean** | 17.69 | 12.29 | 17.98 | 18.72 | 16.63 | 15.57 |

version 3

| | | | | | | | |
|---|---|---|---|---|---|---|---|
| | **Std. Deviation** | 1.45 | 3.61 | 2.02 | 0.81 | 1.79 | 1.56 |
| | **Std. Error of Mean** | 0.72 | 1.47 | 1.01 | 0.4 | 0.68 | 0.78 |
| | **Lower 95% CI of mean** | 15.39 | 8.5 | 14.78 | 17.44 | 14.98 | 13.09 |
| | **Upper 95% CI of mean** | 19.98 | 16.07 | 21.19 | 20.01 | 18.28 | 18.05 |
| **3** | **Number of Values** | 4 | 6 | 4 | 4 | 7 | 4 |
| | **Mean** | 18.74 | 13.31 | 16.13 | 18.59 | 17.78 | 15.34 |
| | **Std. Deviation** | 0.97 | 4.76 | 2.82 | 0.95 | 1.32 | 2.91 |
| | **Std. Error of Mean** | 0.48 | 1.94 | 1.41 | 0.48 | 0.5 | 1.46 |
| | **Lower 95% CI of mean** | 17.2 | 8.31 | 11.64 | 17.08 | 16.56 | 10.7 |
| | **Upper 95% CI of mean** | 20.28 | 18.31 | 20.62 | 20.1 | 19 | 19.97 |
| **3.43** | **Number of Values** | 4 | 6 | 3 | 4 | 7 | 4 |
| | **Mean** | 17.41 | 12.05 | 15.75 | 18.73 | 16.31 | 15.92 |
| | **Std. Deviation** | 2.12 | 3.08 | 1.03 | 0.58 | 3.05 | 2.48 |
| | **Std. Error of Mean** | 1.06 | 1.26 | 0.59 | 0.29 | 1.15 | 1.24 |
| | **Lower 95% CI of mean** | 14.04 | 8.81 | 13.2 | 17.8 | 13.48 | 11.97 |
| | **Upper 95% CI of mean** | 20.77 | 15.28 | 18.29 | 19.65 | 19.13 | 19.86 |

version 3

| | | | | | | | |
|---|---|---|---|---|---|---|---|
| **4** | **Number of Values** | 3 | 5 | 3 | 4 | 5 | 4 |
| | **Mean** | 17.33 | 12.83 | 17.56 | 18.64 | 16.53 | 17.67 |
| | **Std. Deviation** | 1.35 | 4.32 | 0.29 | 1.2 | 0.61 | 2.4 |
| | **Std. Error of Mean** | 0.78 | 1.93 | 0.17 | 0.6 | 0.27 | 1.2 |
| | **Lower 95% CI of mean** | 13.98 | 7.47 | 16.85 | 16.73 | 15.77 | 13.85 |
| | **Upper 95% CI of mean** | 20.67 | 18.19 | 18.27 | 20.55 | 17.28 | 21.49 |
| **5** | **Number of Values** | 3 | 5 | 3 | 4 | 5 | 3 |
| | **Mean** | 16.89 | 14.74 | 14.17 | 17.79 | 16.33 | 17.45 |
| | **Std. Deviation** | 2.78 | 4.38 | 2.49 | 1.9 | 2.35 | 2.29 |
| | **Std. Error of Mean** | 1.6 | 1.96 | 1.44 | 0.95 | 1.05 | 1.32 |
| | **Lower 95% CI of mean** | 10 | 9.31 | 7.99 | 14.77 | 13.42 | 11.76 |
| | **Upper 95% CI of mean** | 23.79 | 20.18 | 20.34 | 20.81 | 19.25 | 23.15 |
| **6** | **Number of Values** | 3 | 5 | 3 | 4 | 5 | 3 |
| | **Mean** | 18.52 | 14.86 | 16.67 | 17.21 | 17.49 | 17.95 |
| | **Std. Deviation** | 0.77 | 3.58 | 2.5 | 0.51 | 0.85 | 1.21 |
| | **Std. Error of Mean** | 0.44 | 1.6 | 1.44 | 0.25 | 0.38 | 0.7 |

version 3

| | | | | | | | |
|---|---|---|---|---|---|---|---|
| | **Lower 95% CI of mean** | 16.62 | 10.42 | 10.47 | 16.4 | 16.44 | 14.96 |
| | **Upper 95% CI of mean** | 20.42 | 19.31 | 22.88 | 18.02 | 18.55 | 20.95 |
| **7** | **Number of Values** | 3 | 5 | 3 | 4 | 5 | 3 |
| | **Mean** | 18.8 | 14.73 | 17.69 | 19.06 | 17.36 | 18.87 |
| | **Std. Deviation** | 1.06 | 3.01 | 1.63 | 0.71 | 2.08 | 0.84 |
| | **Std. Error of Mean** | 0.61 | 1.35 | 0.94 | 0.36 | 0.93 | 0.48 |
| | **Lower 95% CI of mean** | 16.16 | 10.99 | 13.64 | 17.93 | 14.79 | 16.8 |
| | **Upper 95% CI of mean** | 21.44 | 18.47 | 21.75 | 20.19 | 19.94 | 20.95 |
| **8** | **Number of Values** | 3 | 5 | 3 | 4 | 5 | 3 |
| | **Mean** | 17.87 | 14.08 | 17.19 | 17.51 | 17.14 | 17.54 |
| | **Std. Deviation** | 1.5 | 3.26 | 1.26 | 1.63 | 2.21 | 0.68 |
| | **Std. Error of Mean** | 0.86 | 1.46 | 0.73 | 0.81 | 0.99 | 0.39 |
| | **Lower 95% CI of mean** | 14.15 | 10.03 | 14.06 | 14.92 | 14.39 | 15.84 |
| | **Upper 95% CI of mean** | 21.58 | 18.13 | 20.31 | 20.09 | 19.88 | 19.23 |
| **9** | **Number of Values** | 3 | 3 | 3 | 3 | 4 | 3 |
| | **Mean** | 16.5 | 11.53 | 17.01 | 17.84 | 19.44 | 19.03 |

version 3

| | | | | | | | |
|---|---|---|---|---|---|---|---|
| | **Std. Deviation** | 2.08 | 0.24 | 0.58 | 1.61 | 0.63 | 1.27 |
| | **Std. Error of Mean** | 1.2 | 0.14 | 0.33 | 0.93 | 0.31 | 0.74 |
| | **Lower 95% CI of mean** | 11.34 | 10.93 | 15.58 | 13.85 | 18.44 | 15.87 |
| | **Upper 95% CI of mean** | 21.66 | 12.13 | 18.45 | 21.84 | 20.44 | 22.2 |
| **12** | **Number of Values** | 2 | 3 | 3 | 2 | 3 | 3 |
| | **Mean** | 17.59 | 10.78 | 18.08 | 19.67 | 18.22 | 18.07 |
| | **Std. Deviation** | 0.21 | 2.26 | 0.94 | 0.47 | 0.41 | 1.2 |
| | **Std. Error of Mean** | 0.15 | 1.3 | 0.55 | 0.33 | 0.23 | 0.69 |
| | **Lower 95% CI of mean** | 15.67 | 5.18 | 15.73 | 15.44 | 17.21 | 15.1 |
| | **Upper 95% CI of mean** | 19.51 | 16.38 | 20.42 | 23.9 | 19.22 | 21.04 |
| **14** | **Number of Values** | 2 | 1 | 3 | 1 | 2 | 3 |
| | **Mean** | 19.7 | 9.87 | 15.64 | 17.09 | 17.68 | 16.87 |
| | **Std. Deviation** | 0.42 | 0 | 2.02 | 0 | 0.59 | 3.23 |
| | **Std. Error of Mean** | 0.3 | 0 | 1.17 | 0 | 0.42 | 1.87 |
| | **Lower 95% CI of mean** | 15.89 | | 10.62 | | 12.34 | 8.84 |
| | **Upper 95% CI of mean** | 23.51 | | 20.66 | | 23.02 | 24.9 |

version 3

| | | | | | | | |
|---|---|---|---|---|---|---|---|
| **16** | **Number of Values** | 0 | 1 | 3 | 1 | 2 | 2 |
| | **Mean** | | 11.87 | 18.56 | 20 | 19.34 | 19.64 |
| | **Std. Deviation** | | 0 | 1.4 | 0 | 0.42 | 0.52 |
| | **Std. Error of Mean** | | 0 | 0.81 | 0 | 0.3 | 0.37 |
| | **Lower 95% CI of mean** | | | 15.08 | | 15.53 | 15 |
| | **Upper 95% CI of mean** | | | 22.04 | | 23.15 | 24.27 |

version 3

**Supplementary Table 5:** Descriptive statistical summary of modified Hargreaves testing (thermal withdrawal latency) across all experimental groups in female rats, including No Intervention (Control; n = 6), HPI Only (Disease Control; n = 6), Sham FUS (n = 6), FUS Only (n = 6), FUS+HPI (Intervention; n = 7), and LA+HPI (Positive Control; n = 6) at each time point up to 16 weeks. Data are presented as mean, standard deviation (SD), standard error of the mean (SEM), and lower and upper 95% confidence intervals (CI). FUS = focused ultrasound; HPI = hindpaw incision; LA = local anesthetic.

| Time (Weeks) | Comparison | Sum mary | p Value |
|---|---|---|---|
| **-0.29** | **HPI Only vs Control** | ns | .332 |
| | HPI Only vs Sham FUS | ns | >.999 |
| | HPI Only vs FUS Only | ns | .179 |
| | HPI Only vs FUS+HPI | ns | >.999 |
| | HPI Only vs LA+HPI | ns | .497 |
| **-0.14** | **HPI Only vs Control** | ns | .157 |

| | | | |
|---|---|---|---|
| | HPI Only vs Sham FUS | ns | .602 |
| | HPI Only vs FUS Only | * | .017 |
| | HPI Only vs FUS+HPI | ns | >.999 |
| | HPI Only vs LA+HPI | ns | .408 |
| **0** | **HPI Only vs Control** | *** | <.001 |
| | HPI Only vs Sham FUS | *** | <.001 |
| | HPI Only vs FUS Only | *** | <.001 |
| | HPI Only vs FUS+HPI | *** | <.001 |
| | HPI Only vs LA+HPI | *** | <.001 |
| **0.14** | **HPI Only vs Control** | *** | <.001 |
| | **HPI Only vs Sham FUS** | *** | <.001 |
| | **HPI Only vs FUS Only** | *** | <.001 |
| | **HPI Only vs FUS+HPI** | *** | <.001 |
| | **HPI Only vs LA+HPI** | ns | >.999 |
| **0.43** | **HPI Only vs Control** | * | .018 |
| | **HPI Only vs Sham FUS** | ** | .004 |
| | **HPI Only vs FUS Only** | ** | .007 |
| | **HPI Only vs FUS+HPI** | * | .030 |
| | **HPI Only vs LA+HPI** | ns | >.999 |
| **1** | **HPI Only vs Control** | ** | .001 |
| | **HPI Only vs Sham FUS** | ns | .103 |

version 3

| | | | |
|---|---|---|---|
| | **HPI Only vs FUS Only** | * | .025 |
| | **HPI Only vs FUS+HPI** | ** | .009 |
| | **HPI Only vs LA+HPI** | ns | >.999 |
| **1.43** | **HPI Only vs Control** | ** | .004 |
| | **HPI Only vs Sham FUS** | ns | .060 |
| | **HPI Only vs FUS Only** | * | .011 |
| | **HPI Only vs FUS+HPI** | ** | .009 |
| | **HPI Only vs LA+HPI** | ns | >.999 |
| **2** | **HPI Only vs Control** | * | .021 |
| | **HPI Only vs Sham FUS** | * | .014 |
| | **HPI Only vs FUS Only** | * | .012 |
| | **HPI Only vs FUS+HPI** | * | .019 |
| | **HPI Only vs LA+HPI** | ns | >.999 |
| **2.43** | **HPI Only vs Control** | ** | .001 |
| | **HPI Only vs Sham FUS** | *** | <.001 |
| | **HPI Only vs FUS Only** | ** | .005 |
| | **HPI Only vs FUS+HPI** | * | .018 |
| | **HPI Only vs LA+HPI** | * | .047 |
| **3** | **HPI Only vs Control** | ns | .108 |
| | **HPI Only vs Sham FUS** | ns | >.999 |
| | **HPI Only vs FUS Only** | ns | .397 |

version 3

| | | | |
|---|---|---|---|
| | **HPI Only vs FUS+HPI** | ns | .359 |
| | **HPI Only vs LA+HPI** | ns | >.999 |
| **3.43** | **HPI Only vs Control** | ** | .002 |
| | **HPI Only vs Sham FUS** | * | .035 |
| | **HPI Only vs FUS Only** | ** | .003 |
| | **HPI Only vs FUS+HPI** | ns | .123 |
| | **HPI Only vs LA+HPI** | * | .025 |
| **4** | **HPI Only vs Control** | ns | .074 |
| | **HPI Only vs Sham FUS** | ns | .056 |
| | **HPI Only vs FUS Only** | ns | .097 |
| | **HPI Only vs FUS+HPI** | ns | .300 |
| | **HPI Only vs LA+HPI** | ns | .052 |
| **5** | **HPI Only vs Control** | ns | .566 |
| | **HPI Only vs Sham FUS** | ns | >.999 |
| | **HPI Only vs FUS Only** | ns | >.999 |
| | **HPI Only vs FUS+HPI** | ns | >.999 |
| | **HPI Only vs LA+HPI** | ns | .313 |
| **6** | **HPI Only vs Control** | ns | .202 |
| | **HPI Only vs Sham FUS** | ns | >.999 |
| | **HPI Only vs FUS Only** | ns | >.999 |
| | **HPI Only vs FUS+HPI** | ns | >.999 |

version 3

| | | | |
|---|---|---|---|
| | **HPI Only vs LA+HPI** | ns | .390 |
| **7** | **HPI Only vs Control** | * | .011 |
| | **HPI Only vs Sham FUS** | ns | .061 |
| | **HPI Only vs FUS Only** | ns | .211 |
| | **HPI Only vs FUS+HPI** | ns | .300 |
| | **HPI Only vs LA+HPI** | * | .010 |
| **8** | **HPI Only vs Control** | ns | .134 |
| | **HPI Only vs Sham FUS** | ns | .315 |
| | **HPI Only vs FUS Only** | ns | .276 |
| | **HPI Only vs FUS+HPI** | ns | .423 |
| | **HPI Only vs LA+HPI** | ns | .203 |
| **9** | **HPI Only vs Control** | * | .022 |
| | **HPI Only vs Sham FUS** | * | .012 |
| | **HPI Only vs FUS Only** | ns | .261 |
| | **HPI Only vs FUS+HPI** | ** | .001 |
| | **HPI Only vs LA+HPI** | ** | .001 |
| **12** | **HPI Only vs Control** | * | .020 |
| | **HPI Only vs Sham FUS** | ** | .006 |
| | **HPI Only vs FUS Only** | * | .017 |
| | **HPI Only vs FUS+HPI** | ** | .005 |
| | **HPI Only vs LA+HPI** | ** | .006 |

version 3

| | | | |
|---|---|---|---|
| **14** | **HPI Only vs Control** | ns | .765 |
| | **HPI Only vs Sham FUS** | ns | >.999 |
| | **HPI Only vs FUS Only** | ns | .661 |
| | **HPI Only vs FUS+HPI** | ns | .601 |
| | **HPI Only vs LA+HPI** | ns | >.999 |
| **16** | **HPI Only vs Control** | ns | .100 |
| | **HPI Only vs Sham FUS** | ns | .791 |
| | **HPI Only vs FUS Only** | ns | >.999 |
| | **HPI Only vs FUS+HPI** | ns | >.999 |
| | **HPI Only vs LA+HPI** | ns | .892 |

version 3

**Supplementary Table 6:** Summary of group-wise comparisons for modified Hargreaves testing (thermal withdrawal latency) across all experimental groups in female rats, including No Intervention (Control; n = 6), HPI Only (Disease Control; n = 6), Sham FUS (n = 6), FUS Only (n = 6), FUS+HPI (Intervention; n = 7), and LA+HPI (Positive Control; n = 6) at each time point up to 16 weeks. Measurements were analyzed using a two-way mixed-effects ANOVA with fixed effects for treatment group and time, followed by Bonferroni-corrected post hoc multiple-comparison testing. Statistical significance is denoted as follows: ns ($p \geq 0.05$), * ($p < 0.05$), ** ($p < 0.01$), *** ($p < 0.001$).**Supplementary Table 7:** Descriptive statistical summary of Randall–Selitto testing (hindpaw mechanical withdrawal thresholds) across all experimental

| Time (Weeks) | Study Arms | Control | HPI Only | Sham FUS | FUS Only | FUS+HPI | LA+HPI |
|---|---|---|---|---|---|---|---|
| **-0.29** | **Number of Values (N)** | 12 | 12 | 12 | 12 | 12 | 12 |
| | **Mean** | 201.3 | 212.3 | 219.2 | 214.0 | 209.0 | 214.1 |
| | **Std. Deviation** | 24.94 | 11.65 | 22.60 | 19.16 | 22.16 | 17.02 |

| | | | | | | | |
|---|---|---|---|---|---|---|---|
| | **Std. Error of Mean** | 7.199 | 3.364 | 6.524 | 5.531 | 6.396 | 4.912 |
| | **Lower 95% CI of mean** | 185.5 | 204.9 | 204.9 | 201.8 | 194.9 | 203.3 |
| | **Upper 95% CI of mean** | 217.1 | 219.7 | 233.6 | 226.2 | 223.1 | 224.9 |
| **-0.14** | **Number of Values** | 12 | 12 | 12 | 12 | 12 | 12 |
| | **Mean** | 226.0 | 220.4 | 211.4 | 211.9 | 197.3 | 207.9 |
| | **Std. Deviation** | 7.632 | 13.75 | 19.23 | 22.04 | 25.12 | 16.80 |
| | **Std. Error of Mean** | 2.203 | 3.971 | 5.551 | 6.363 | 7.252 | 4.851 |
| | **Lower 95% CI of mean** | 221.1 | 211.7 | 199.2 | 197.9 | 181.3 | 197.2 |
| | **Upper 95% CI of mean** | 230.8 | 229.2 | 223.6 | 225.9 | 213.3 | 218.6 |
| **0** | **Number of Values** | 12 | 12 | 12 | 12 | 12 | 12 |
| | **Mean** | 221.0 | 37.44 | 229.2 | 238.4 | 217.1 | 226.8 |
| | **Std. Deviation** | 14.16 | 11.13 | 14.90 | 10.22 | 29.50 | 13.33 |
| | **Std. Error of Mean** | 4.087 | 3.214 | 4.302 | 2.950 | 8.515 | 3.847 |
| | **Lower 95% CI of mean** | 212.0 | 30.36 | 219.8 | 232.0 | 198.4 | 218.3 |
| | **Upper 95% CI of mean** | 229.9 | 44.51 | 238.7 | 244.9 | 235.8 | 235.3 |
| **0.14** | **Number of Values** | 12 | 12 | 12 | 12 | 12 | 12 |
| | **Mean** | 215.0 | 57.71 | 226.8 | 238.0 | 217.0 | 94.93 |

version 3

| | | | | | | | |
|---|---|---|---|---|---|---|---|
| | **Std. Deviation** | 15.27 | 17.51 | 18.55 | 13.41 | 35.41 | 26.44 |
| | **Std. Error of Mean** | 4.407 | 5.055 | 5.356 | 3.872 | 10.22 | 7.634 |
| | **Lower 95% CI of mean** | 205.3 | 46.59 | 215.0 | 229.5 | 194.5 | 78.13 |
| | **Upper 95% CI of mean** | 224.7 | 68.84 | 238.6 | 246.6 | 239.5 | 111.7 |
| **0.43** | **Number of Values** | 12 | 12 | 12 | 12 | 12 | 12 |
| | **Mean** | 207.4 | 113.6 | 227.7 | 232.5 | 222.2 | 105.7 |
| | **Std. Deviation** | 32.21 | 36.14 | 14.70 | 16.18 | 25.28 | 44.43 |
| | **Std. Error of Mean** | 9.297 | 10.43 | 4.242 | 4.672 | 7.299 | 12.82 |
| | **Lower 95% CI of mean** | 186.9 | 90.62 | 218.4 | 222.2 | 206.2 | 77.48 |
| | **Upper 95% CI of mean** | 227.8 | 136.5 | 237.1 | 242.8 | 238.3 | 133.9 |
| **1** | **Number of Values** | 12 | 12 | 12 | 12 | 12 | 12 |
| | **Mean** | 191.2 | 136.4 | 216.2 | 224.9 | 215.4 | 126.4 |
| | **Std. Deviation** | 28.79 | 33.05 | 16.47 | 18.60 | 28.21 | 14.53 |
| | **Std. Error of Mean** | 8.310 | 9.541 | 4.756 | 5.369 | 8.143 | 4.194 |
| | **Lower 95% CI of mean** | 172.9 | 115.4 | 205.7 | 213.1 | 197.5 | 117.1 |
| | **Upper 95% CI of mean** | 209.5 | 157.4 | 226.6 | 236.7 | 233.3 | 135.6 |
| **1.43** | **Number of Values** | 10 | 12 | 12 | 12 | 12 | 10 |

version 3

| | | | | | | | |
|---|---|---|---|---|---|---|---|
| | **Mean** | 196.7 | 154.7 | 220.3 | 222.2 | 219.4 | 148.1 |
| | **Std. Deviation** | 25.90 | 51.24 | 12.90 | 23.69 | 25.31 | 25.67 |
| | **Std. Error of Mean** | 8.190 | 14.79 | 3.724 | 6.839 | 7.307 | 8.116 |
| | **Lower 95% CI of mean** | 178.2 | 122.2 | 212.1 | 207.2 | 203.3 | 129.8 |
| | **Upper 95% CI of mean** | 215.2 | 187.3 | 228.5 | 237.3 | 235.5 | 166.5 |
| **2** | **Number of Values** | 10 | 10 | 12 | 10 | 11 | 10 |
| | **Mean** | 196.7 | 154.7 | 220.3 | 222.2 | 219.4 | 148.1 |
| | **Std. Deviation** | 25.90 | 51.24 | 12.90 | 23.69 | 25.31 | 25.67 |
| | **Std. Error of Mean** | 8.190 | 14.79 | 3.724 | 6.839 | 7.307 | 8.116 |
| | **Lower 95% CI of mean** | 178.2 | 122.2 | 212.1 | 207.2 | 203.3 | 129.8 |
| | **Upper 95% CI of mean** | 215.2 | 187.3 | 228.5 | 237.3 | 235.5 | 166.5 |
| **2.43** | **Number of Values** | 10 | 10 | 10 | 10 | 10 | 10 |
| | **Mean** | 203.0 | 139.9 | 212.8 | 226.1 | 228.8 | 161.5 |
| | **Std. Deviation** | 18.78 | 32.50 | 8.300 | 12.37 | 16.68 | 36.19 |
| | **Std. Error of Mean** | 5.940 | 10.28 | 2.625 | 3.912 | 5.275 | 11.44 |
| | **Lower 95% CI of mean** | 189.5 | 116.7 | 206.9 | 217.3 | 216.9 | 135.6 |
| | **Upper 95% CI of mean** | 216.4 | 163.2 | 218.7 | 235.0 | 240.7 | 187.3 |

version 3

| | | | | | | | |
|---|---|---|---|---|---|---|---|
| **3** | **Number of Values** | 10 | 10 | 10 | 10 | 10 | 10 |
| | **Mean** | 220.8 | 155.1 | 202.0 | 231.2 | 221.7 | 176.6 |
| | **Std. Deviation** | 10.89 | 27.57 | 18.25 | 9.949 | 19.32 | 31.47 |
| | **Std. Error of Mean** | 3.444 | 8.720 | 5.770 | 3.316 | 6.109 | 9.951 |
| | **Lower 95% CI of mean** | 213.0 | 135.3 | 188.9 | 223.6 | 207.8 | 154.1 |
| | **Upper 95% CI of mean** | 228.6 | 174.8 | 215.0 | 238.9 | 235.5 | 199.1 |
| **3.43** | **Number of Values** | 10 | 10 | 10 | 10 | 10 | 10 |
| | **Mean** | 221.6 | 155.5 | 215.1 | 229.7 | 213.3 | 177.6 |
| | **Std. Deviation** | 13.96 | 22.48 | 23.30 | 13.95 | 23.41 | 34.91 |
| | **Std. Error of Mean** | 4.413 | 7.109 | 7.766 | 4.650 | 7.803 | 11.64 |
| | **Lower 95% CI of mean** | 211.6 | 139.4 | 197.2 | 219.0 | 195.3 | 150.7 |
| | **Upper 95% CI of mean** | 231.6 | 171.6 | 233.0 | 240.4 | 231.3 | 204.4 |
| **4** | **Number of Values** | 8 | 8 | 8 | 8 | 8 | 8 |
| | **Mean** | 198.4 | 164.0 | 220.0 | 221.5 | 207.9 | 196.3 |
| | **Std. Deviation** | 9.262 | 28.82 | 23.24 | 18.53 | 23.11 | 13.76 |
| | **Std. Error of Mean** | 3.275 | 10.19 | 8.217 | 6.176 | 8.170 | 4.864 |
| | **Lower 95% CI of mean** | 190.7 | 139.9 | 200.6 | 207.2 | 188.6 | 184.8 |

version 3

| | | | | | | | |
|---|---|---|---|---|---|---|---|
| | **Upper 95% CI of mean** | 206.2 | 188.1 | 239.4 | 235.7 | 227.2 | 207.9 |
| **5** | **Number of Values** | 8 | 8 | 8 | 8 | 8 | 8 |
| | **Mean** | 210.2 | 181.9 | 213.0 | 216.5 | 213.1 | 190.2 |
| | **Std. Deviation** | 19.88 | 33.60 | 23.93 | 8.007 | 15.67 | 10.76 |
| | **Std. Error of Mean** | 7.030 | 11.88 | 8.459 | 2.669 | 5.541 | 4.394 |
| | **Lower 95% CI of mean** | 193.6 | 153.8 | 193.0 | 210.3 | 200.0 | 178.9 |
| | **Upper 95% CI of mean** | 226.8 | 210.0 | 233.0 | 222.6 | 226.2 | 201.4 |
| **6** | **Number of Values** | 8 | 8 | 8 | 8 | 8 | 8 |
| | **Mean** | 208.4 | 188.3 | 210.8 | 222.5 | 199.0 | 199.5 |
| | **Std. Deviation** | 12.07 | 29.51 | 22.06 | 13.55 | 24.02 | 7.079 |
| | **Std. Error of Mean** | 4.926 | 10.43 | 7.799 | 4.789 | 8.491 | 2.890 |
| | **Lower 95% CI of mean** | 195.7 | 163.6 | 192.4 | 211.2 | 178.9 | 192.1 |
| | **Upper 95% CI of mean** | 221.1 | 212.9 | 229.2 | 233.8 | 219.1 | 207.0 |
| **7** | **Number of Values** | 8 | 8 | 8 | 8 | 8 | 8 |
| | **Mean** | 211.9 | 198.1 | 220.2 | 216.8 | 208.9 | 191.0 |
| | **Std. Deviation** | 8.048 | 25.33 | 15.11 | 19.93 | 19.16 | 26.40 |
| | **Std. Error of Mean** | 3.286 | 8.956 | 5.712 | 7.532 | 6.774 | 10.78 |

version 3

| | | | | | | | |
|---|---|---|---|---|---|---|---|
| | **Lower 95% CI of mean** | 203.4 | 176.9 | 206.2 | 198.4 | 192.9 | 163.3 |
| | **Upper 95% CI of mean** | 220.3 | 219.3 | 234.2 | 235.2 | 224.9 | 218.7 |
| **8** | **Number of Values** | 8 | 8 | 8 | 8 | 8 | 8 |
| | **Mean** | 215.9 | 208.0 | 216.4 | 227.4 | 217.2 | 210.3 |
| | **Std. Deviation** | 7.256 | 27.08 | 12.13 | 17.36 | 20.04 | 13.01 |
| | **Std. Error of Mean** | 2.962 | 9.575 | 4.585 | 6.563 | 7.085 | 5.313 |
| | **Lower 95% CI of mean** | 208.3 | 185.4 | 205.2 | 211.4 | 200.4 | 196.6 |
| | **Upper 95% CI of mean** | 223.5 | 230.7 | 227.6 | 243.5 | 233.9 | 223.9 |
| **9** | **Number of Values** | 6 | 6 | 6 | 6 | 6 | 6 |
| | **Mean** | 210.8 | 190.3 | 215.0 | 211.9 | 207.6 | 209.4 |
| | **Std. Deviation** | 23.84 | 6.584 | 9.561 | 10.01 | 12.83 | 19.07 |
| | **Std. Error of Mean** | 9.734 | 2.688 | 3.903 | 4.086 | 5.238 | 7.786 |
| | **Lower 95% CI of mean** | 185.8 | 183.4 | 205.0 | 201.4 | 194.1 | 189.4 |
| | **Upper 95% CI of mean** | 235.9 | 197.2 | 225.0 | 222.4 | 221.0 | 229.4 |
| **12** | **Number of Values** | 6 | 6 | 6 | 6 | 6 | 6 |
| | **Mean** | 201.6 | 182.8 | 202.6 | 208.9 | 199.3 | 208.0 |
| | **Std. Deviation** | 11.94 | 19.47 | 17.75 | 13.12 | 8.865 | 6.341 |

version 3

| | | | | | | | |
|---|---|---|---|---|---|---|---|
| | **Std. Error of Mean** | 4.876 | 7.947 | 7.245 | 5.355 | 3.965 | 2.589 |
| | **Lower 95% CI of mean** | 189.0 | 162.4 | 184.0 | 195.2 | 188.3 | 201.3 |
| | **Upper 95% CI of mean** | 214.1 | 203.3 | 221.3 | 222.7 | 210.3 | 214.6 |
| **14** | **Number of Values** | 4 | 4 | 4 | 4 | 4 | 4 |
| | **Mean** | 210.3 | 200.3 | 227.0 | 217.9 | 223.4 | 211.2 |
| | **Std. Deviation** | 9.373 | 21.80 | 23.35 | 8.465 | 12.95 | 25.23 |
| | **Std. Error of Mean** | 4.192 | 10.90 | 9.531 | 3.456 | 6.477 | 11.29 |
| | **Lower 95% CI of mean** | 198.7 | 165.6 | 202.5 | 209.0 | 202.8 | 179.8 |
| | **Upper 95% CI of mean** | 221.9 | 235.0 | 251.5 | 226.8 | 244.0 | 242.5 |
| **16** | **Number of Values** | 4 | 4 | 4 | 4 | 4 | 4 |
| | **Mean** | 219.7 | 196.4 | 218.2 | 222.8 | 206.9 | 218.9 |
| | **Std. Deviation** | 20.08 | 9.414 | 26.51 | 17.68 | 7.587 | 17.19 |
| | **Std. Error of Mean** | 10.04 | 4.707 | 13.25 | 10.21 | 5.365 | 8.597 |
| | **Lower 95% CI of mean** | 187.7 | 181.4 | 176.0 | 178.9 | 138.8 | 191.5 |
| | **Upper 95% CI of mean** | 251.6 | 211.4 | 260.3 | 266.8 | 275.1 | 246.3 |

groups, including No Intervention (Control; n = 12), HPI Only (Disease Control; n = 12), Sham FUS (n = 12), FUS Only (n = 12), FUS+HPI (Intervention; n = 12), and LA+HPI (Positive Control; n = 12) at each time point up to 16 weeks. Data are presented as mean, standard

version 3

deviation (SD), standard error of the mean (SEM), and lower and upper 95% confidence intervals (CI). FUS = focused ultrasound; HPI = hindpaw incision; LA = local anesthetic.

| Time (Weeks) | Comparison | Summary | p Value |
|---|---|---|---|
| **-0.29** | **HPI Only vs Control** | ns | >.999 |
| | **HPI Only vs Sham FUS** | ns | >.999 |
| | **HPI Only vs FUS Only** | ns | >.999 |
| | **HPI Only vs FUS+HPI** | ns | >.999 |
| | **HPI Only vs LA+HPI** | ns | >.999 |
| **-0.14** | **HPI Only vs Control** | ns | >.999 |
| | **HPI Only vs Sham FUS** | ns | >.999 |
| | **HPI Only vs FUS Only** | ns | >.999 |
| | **HPI Only vs FUS+HPI** | ns | .054 |
| | **HPI Only vs LA+HPI** | ns | >.999 |
| **0** | **HPI Only vs Control** | *** | <.001 |
| | **HPI Only vs Sham FUS** | *** | <.001 |
| | **HPI Only vs FUS Only** | *** | <.001 |
| | **HPI Only vs FUS+HPI** | *** | <.001 |
| | **HPI Only vs LA+HPI** | *** | <.001 |
| **0.14** | **HPI Only vs Control** | *** | <.001 |
| | **HPI Only vs Sham FUS** | *** | <.001 |

version 3

| | | | |
|---|---|---|---|
| | **HPI Only vs FUS Only** | *** | <.001 |
| | **HPI Only vs FUS+HPI** | *** | <.001 |
| | **HPI Only vs LA+HPI** | ** | .003 |
| **0.43** | **HPI Only vs Control** | *** | <.001 |
| | **HPI Only vs Sham FUS** | *** | <.001 |
| | **HPI Only vs FUS Only** | *** | <.001 |
| | **HPI Only vs FUS+HPI** | *** | <.001 |
| | **HPI Only vs LA+HPI** | ns | >.999 |
| **1** | **HPI Only vs Control** | *** | <.001 |
| | **HPI Only vs Sham FUS** | *** | <.001 |
| | **HPI Only vs FUS Only** | *** | <.001 |
| | **HPI Only vs FUS+HPI** | *** | <.001 |
| | **HPI Only vs LA+HPI** | ns | >.999 |
| **1.43** | **HPI Only vs Control** | ** | .004 |
| | **HPI Only vs Sham FUS** | *** | <.001 |
| | **HPI Only vs FUS Only** | *** | <.001 |
| | **HPI Only vs FUS+HPI** | *** | <.001 |
| | **HPI Only vs LA+HPI** | ns | >.999 |
| **2** | **HPI Only vs Control** | *** | <.001 |

version 3

| | | | |
|---|---|---|---|
| | **HPI Only vs Sham FUS** | *** | <.001 |
| | **HPI Only vs FUS Only** | *** | <.001 |
| | **HPI Only vs FUS+HPI** | *** | <.001 |
| | **HPI Only vs LA+HPI** | ns | >.999 |
| **2.43** | **HPI Only vs Control** | *** | <.001 |
| | **HPI Only vs Sham FUS** | *** | <.001 |
| | **HPI Only vs FUS Only** | *** | <.001 |
| | **HPI Only vs FUS+HPI** | *** | <.001 |
| | **HPI Only vs LA+HPI** | ns | >.999 |
| **3** | **HPI Only vs Control** | *** | <.001 |
| | **HPI Only vs Sham FUS** | *** | <.001 |
| | **HPI Only vs FUS Only** | *** | <.001 |
| | **HPI Only vs FUS+HPI** | *** | <.001 |
| | **HPI Only vs LA+HPI** | ns | .460 |
| **3.43** | **HPI Only vs Control** | *** | <.001 |
| | **HPI Only vs Sham FUS** | *** | <.001 |
| | **HPI Only vs FUS Only** | *** | <.001 |
| | **HPI Only vs FUS+HPI** | *** | <.001 |
| | **HPI Only vs LA+HPI** | ns | .692 |

version 3

| | | | |
|---|---|---|---|
| **4** | **HPI Only vs Control** | * | .011 |
| | **HPI Only vs Sham FUS** | *** | <.001 |
| | **HPI Only vs FUS Only** | *** | <.001 |
| | **HPI Only vs FUS+HPI** | *** | <.001 |
| | **HPI Only vs LA+HPI** | * | .012 |
| **5** | **HPI Only vs Control** | ns | .175 |
| | **HPI Only vs Sham FUS** | ns | .064 |
| | **HPI Only vs FUS Only** | ns | .099 |
| | **HPI Only vs FUS+HPI** | ns | .058 |
| | **HPI Only vs LA+HPI** | ns | >.999 |
| **6** | **HPI Only vs Control** | ns | >.999 |
| | **HPI Only vs Sham FUS** | ns | .637 |
| | **HPI Only vs FUS Only** | ns | .127 |
| | **HPI Only vs FUS+HPI** | ns | >.999 |
| | **HPI Only vs LA+HPI** | ns | >.999 |
| **7** | **HPI Only vs Control** | ns | >.999 |
| | **HPI Only vs Sham FUS** | ns | .537 |
| | **HPI Only vs FUS Only** | ns | >.999 |
| | **HPI Only vs FUS+HPI** | ns | >.999 |

version 3

| | | | |
|---|---|---|---|
| | **HPI Only vs LA+HPI** | ns | >.999 |
| **8** | **HPI Only vs Control** | ns | >.999 |
| | **HPI Only vs Sham FUS** | ns | >.999 |
| | **HPI Only vs FUS Only** | ns | >.999 |
| | **HPI Only vs FUS+HPI** | ns | >.999 |
| | **HPI Only vs LA+HPI** | ns | >.999 |
| **9** | **HPI Only vs Control** | ns | >.999 |
| | **HPI Only vs Sham FUS** | ns | .066 |
| | **HPI Only vs FUS Only** | ns | .507 |
| | **HPI Only vs FUS+HPI** | ns | >.999 |
| | **HPI Only vs LA+HPI** | ns | .273 |
| **12** | **HPI Only vs Control** | ns | >.999 |
| | **HPI Only vs Sham FUS** | ns | .152 |
| | **HPI Only vs FUS Only** | ns | .414 |
| | **HPI Only vs FUS+HPI** | ns | .951 |
| | **HPI Only vs LA+HPI** | ns | .119 |
| **14** | **HPI Only vs Control** | ns | >.999 |
| | **HPI Only vs Sham FUS** | ns | >.999 |
| | **HPI Only vs FUS Only** | ns | >.999 |

version 3

| | | | |
|---|---|---|---|
| | **HPI Only vs FUS+HPI** | ns | >.999 |
| | **HPI Only vs LA+HPI** | ns | >.999 |
| **16** | **HPI Only vs Control** | ns | .910 |
| | **HPI Only vs Sham FUS** | ns | >.999 |
| | **HPI Only vs FUS Only** | ns | >.999 |
| | **HPI Only vs FUS+HPI** | ns | >.999 |
| | **HPI Only vs LA+HPI** | ns | >.999 |

version 3

**Supplementary Table 8:** Summary of group-wise comparisons for Randall–Selitto testing (hindpaw mechanical withdrawal thresholds) across all experimental groups, including No Intervention (Control; n = 12), HPI Only (Disease Control; n = 12), Sham FUS (n = 12), FUS Only (n = 12), FUS+HPI (Intervention; n = 12), and LA+HPI (Positive Control; n = 12) at each time point up to 16 weeks. Measurements were analyzed using a two-way mixed-effects ANOVA with fixed effects for treatment group and time, followed by Bonferroni-corrected post hoc multiple-comparison testing. Statistical significance is denoted as follows: ns ($p \geq 0.05$), * ($p < 0.05$), ** ($p < 0.01$), *** ($p < 0.001$).

| Time (Weeks) | Study Arms | Control | HPI Only | Sham FUS | FUS Only | FUS+HPI | LA+HPI |
|---|---|---|---|---|---|---|---|
| **-0.29** | **Number of Values (N)** | 6 | 6 | 6 | 6 | 4 | 6 |
| | **Mean** | 219.7 | 216.3 | 227.7 | 211 | 208.2 | 213.2 |
| | **Std. Deviation** | 13.42 | 15.89 | 6.8 | 21.37 | 16.89 | 21.61 |
| | **Std. Error of Mean** | 5.48 | 6.49 | 2.78 | 8.73 | 8.44 | 8.82 |
| | **Lower 95% CI of mean** | 205.6 | 199.6 | 220.6 | 188.5 | 181.3 | 190.5 |

| | | | | | | | |
|---|---|---|---|---|---|---|---|
| | **Upper 95% CI of mean** | 233.8 | 233 | 234.9 | 233.4 | 235.1 | 235.9 |
| **-0.14** | **Number of Values** | 6 | 6 | 6 | 6 | 4 | 6 |
| | **Mean** | 222.9 | 228 | 209.1 | 207 | 216.4 | 210.5 |
| | **Std. Deviation** | 5.5 | 14.61 | 16.42 | 24.01 | 4.58 | 10.62 |
| | **Std. Error of Mean** | 2.24 | 5.97 | 6.7 | 9.8 | 2.29 | 4.34 |
| | **Lower 95% CI of mean** | 217.1 | 212.6 | 191.9 | 181.8 | 209.1 | 199.4 |
| | **Upper 95% CI of mean** | 228.6 | 243.3 | 226.4 | 232.2 | 223.7 | 221.7 |
| **0** | **Number of Values** | 6 | 6 | 6 | 6 | 4 | 6 |
| | **Mean** | 219 | 31.16 | 234.9 | 244.2 | 220.1 | 228.3 |
| | **Std. Deviation** | 10.13 | 5.15 | 9.15 | 5.69 | 23.43 | 13.98 |
| | **Std. Error of Mean** | 4.13 | 2.1 | 3.74 | 2.32 | 11.72 | 5.71 |
| | **Lower 95% CI of mean** | 208.4 | 25.75 | 225.3 | 238.3 | 182.8 | 213.6 |
| | **Upper 95% CI of mean** | 229.7 | 36.56 | 244.5 | 250.2 | 257.4 | 243 |
| **0.14** | **Number of Values** | 6 | 6 | 6 | 6 | 4 | 6 |
| | **Mean** | 220.3 | 60.5 | 232.5 | 240.7 | 207.8 | 107 |
| | **Std. Deviation** | 18.7 | 21.1 | 17.37 | 12.86 | 46.72 | 32.8 |
| | **Std. Error of Mean** | 7.63 | 8.62 | 7.09 | 5.25 | 23.36 | 13.39 |

version 3

| | | | | | | | |
|---|---|---|---|---|---|---|---|
| | **Lower 95% CI of mean** | 200.7 | 38.35 | 214.2 | 227.2 | 133.5 | 72.58 |
| | **Upper 95% CI of mean** | 239.9 | 82.65 | 250.7 | 254.2 | 282.2 | 141.4 |
| **0.43** | **Number of Values** | 6 | 6 | 6 | 6 | 4 | 6 |
| | **Mean** | 202.9 | 97.31 | 237.4 | 236.1 | 229 | 98.88 |
| | **Std. Deviation** | 41.09 | 35.82 | 3.25 | 12.81 | 21.95 | 15.07 |
| | **Std. Error of Mean** | 16.78 | 14.62 | 1.33 | 5.23 | 10.98 | 6.15 |
| | **Lower 95% CI of mean** | 159.8 | 59.71 | 234 | 222.7 | 194.1 | 83.06 |
| | **Upper 95% CI of mean** | 246 | 134.9 | 240.9 | 249.6 | 264 | 114.7 |
| **1** | **Number of Values** | 6 | 6 | 6 | 6 | 4 | 6 |
| | **Mean** | 192 | 118.7 | 219.9 | 236.1 | 219.7 | 126 |
| | **Std. Deviation** | 40.24 | 25.66 | 21.4 | 13.09 | 31.13 | 20.92 |
| | **Std. Error of Mean** | 16.43 | 10.48 | 8.74 | 5.34 | 15.56 | 8.54 |
| | **Lower 95% CI of mean** | 149.7 | 91.77 | 197.4 | 222.4 | 170.1 | 104.1 |
| | **Upper 95% CI of mean** | 234.2 | 145.6 | 242.3 | 249.8 | 269.2 | 148 |
| **1.43** | **Number of Values** | 6 | 6 | 6 | 6 | 4 | 6 |
| | **Mean** | 195.9 | 142.3 | 220 | 237 | 219.1 | 151.2 |
| | **Std. Deviation** | 32.24 | 64.17 | 18.01 | 15.78 | 24.83 | 22.69 |

version 3

| | | | | | | | |
|---|---|---|---|---|---|---|---|
| | **Std. Error of Mean** | 13.16 | 26.2 | 7.35 | 6.44 | 12.42 | 9.27 |
| | **Lower 95% CI of mean** | 162 | 74.92 | 201.1 | 220.4 | 179.6 | 127.3 |
| | **Upper 95% CI of mean** | 229.7 | 209.6 | 238.9 | 253.5 | 258.6 | 175 |
| **2** | **Number of Values** | 6 | 4 | 6 | 6 | 3 | 6 |
| | **Mean** | 199.2 | 118.8 | 221.2 | 235.6 | 211.6 | 159.4 |
| | **Std. Deviation** | 34.05 | 41.31 | 17.49 | 7.72 | 30 | 31.39 |
| | **Std. Error of Mean** | 13.9 | 20.66 | 7.14 | 3.15 | 17.32 | 12.81 |
| | **Lower 95% CI of mean** | 163.5 | 53.04 | 202.8 | 227.5 | 137.1 | 126.4 |
| | **Upper 95% CI of mean** | 235 | 184.5 | 239.5 | 243.7 | 286.2 | 192.3 |
| **2.43** | **Number of Values** | 6 | 4 | 6 | 6 | 3 | 6 |
| | **Mean** | 194.6 | 142.3 | 213 | 226.9 | 224 | 159.8 |
| | **Std. Deviation** | 11.67 | 53.2 | 10.22 | 16.18 | 21.76 | 36.46 |
| | **Std. Error of Mean** | 4.76 | 26.6 | 4.17 | 6.61 | 12.57 | 14.89 |
| | **Lower 95% CI of mean** | 182.3 | 57.63 | 202.2 | 209.9 | 169.9 | 121.6 |
| | **Upper 95% CI of mean** | 206.8 | 226.9 | 223.7 | 243.8 | 278 | 198.1 |
| **3** | **Number of Values** | 6 | 4 | 6 | 5 | 3 | 6 |
| | **Mean** | 218.3 | 136.5 | 205.3 | 232.9 | 226.3 | 171.8 |

version 3

| | | | | | | | |
|---|---|---|---|---|---|---|---|
| | **Std. Deviation** | 11.66 | 35.3 | 16.77 | 12.45 | 11.48 | 36.27 |
| | **Std. Error of Mean** | 4.76 | 17.65 | 6.85 | 5.57 | 6.63 | 14.81 |
| | **Lower 95% CI of mean** | 206.1 | 80.35 | 187.7 | 217.5 | 197.8 | 133.7 |
| | **Upper 95% CI of mean** | 230.6 | 192.7 | 222.9 | 248.4 | 254.8 | 209.9 |
| **3.43** | **Number of Values** | 6 | 4 | 6 | 5 | 3 | 5 |
| | **Mean** | 220.3 | 150.7 | 220.2 | 231.6 | 205.3 | 162.3 |
| | **Std. Deviation** | 15.24 | 34.8 | 24.27 | 14.49 | 25.84 | 35.13 |
| | **Std. Error of Mean** | 6.22 | 17.4 | 9.91 | 6.48 | 14.92 | 15.71 |
| | **Lower 95% CI of mean** | 204.3 | 95.35 | 194.7 | 213.6 | 141.1 | 118.7 |
| | **Upper 95% CI of mean** | 236.3 | 206.1 | 245.7 | 249.6 | 269.5 | 205.9 |
| **4** | **Number of Values** | 5 | 3 | 5 | 5 | 3 | 5 |
| | **Mean** | 195.6 | 149.1 | 234.7 | 230.4 | 198.3 | 198.4 |
| | **Std. Deviation** | 10.31 | 23.84 | 12.96 | 7.88 | 29.41 | 10.97 |
| | **Std. Error of Mean** | 4.61 | 13.76 | 5.8 | 3.52 | 16.98 | 4.91 |
| | **Lower 95% CI of mean** | 182.8 | 89.92 | 218.6 | 220.6 | 125.3 | 184.8 |
| | **Upper 95% CI of mean** | 208.4 | 208.3 | 250.8 | 240.2 | 271.4 | 212 |
| **5** | **Number of Values** | 5 | 3 | 5 | 5 | 3 | 3 |

version 3

| | | | | | | | |
|---|---|---|---|---|---|---|---|
| | **Mean** | 210.5 | 162.4 | 221.8 | 220.7 | 220.5 | 191.5 |
| | **Std. Deviation** | 24.33 | 5.92 | 17.19 | 6.72 | 8.53 | 10.02 |
| | **Std. Error of Mean** | 10.88 | 3.42 | 7.69 | 3.01 | 4.93 | 5.78 |
| | **Lower 95% CI of mean** | 180.3 | 147.7 | 200.4 | 212.3 | 199.3 | 166.6 |
| | **Upper 95% CI of mean** | 240.7 | 177.1 | 243.1 | 229 | 241.7 | 216.4 |
| **6** | **Number of Values** | 3 | 3 | 5 | 4 | 3 | 3 |
| | **Mean** | 210 | 166.5 | 222.9 | 224.1 | 198.3 | 204.9 |
| | **Std. Deviation** | 12.55 | 12.41 | 14.56 | 19.68 | 10.6 | 4.04 |
| | **Std. Error of Mean** | 7.25 | 7.17 | 6.51 | 9.84 | 6.12 | 2.34 |
| | **Lower 95% CI of mean** | 178.8 | 135.7 | 204.8 | 192.8 | 172 | 194.9 |
| | **Upper 95% CI of mean** | 241.2 | 197.3 | 241 | 255.5 | 224.6 | 215 |
| **7** | **Number of Values** | 3 | 3 | 4 | 4 | 3 | 3 |
| | **Mean** | 209.4 | 202.4 | 219.1 | 223.8 | 203.2 | 182.7 |
| | **Std. Deviation** | 4.16 | 14.4 | 19.07 | 22.16 | 22.33 | 30.73 |
| | **Std. Error of Mean** | 2.4 | 8.32 | 9.53 | 11.08 | 12.89 | 17.74 |
| | **Lower 95% CI of mean** | 199 | 166.6 | 188.8 | 188.5 | 147.8 | 106.4 |
| | **Upper 95% CI of mean** | 219.7 | 238.2 | 249.4 | 259 | 258.7 | 259.1 |

version 3

| | | | | | | | |
|---|---|---|---|---|---|---|---|
| **8** | **Number of Values** | 3 | 3 | 4 | 4 | 3 | 3 |
| | **Mean** | 213.8 | 221 | 219.9 | 236.4 | 218.2 | 208.6 |
| | **Std. Deviation** | 9.85 | 17.51 | 13.04 | 17.64 | 29.67 | 3.91 |
| | **Std. Error of Mean** | 5.69 | 10.11 | 6.52 | 8.82 | 17.13 | 2.26 |
| | **Lower 95% CI of mean** | 189.4 | 177.5 | 199.1 | 208.4 | 144.5 | 198.9 |
| | **Upper 95% CI of mean** | 238.3 | 264.5 | 240.6 | 264.5 | 291.9 | 218.3 |
| **9** | **Number of Values** | 3 | 3 | 3 | 4 | 2 | 3 |
| | **Mean** | 214.6 | 191.5 | 208.2 | 216.2 | 208.6 | 206.3 |
| | **Std. Deviation** | 15.86 | 4.36 | 8.74 | 9.69 | 2.6 | 20.97 |
| | **Std. Error of Mean** | 9.16 | 2.52 | 5.05 | 4.84 | 1.84 | 12.11 |
| | **Lower 95% CI of mean** | 175.2 | 180.7 | 186.5 | 200.8 | 185.2 | 154.2 |
| | **Upper 95% CI of mean** | 254 | 202.4 | 229.9 | 231.6 | 231.9 | 258.4 |
| **12** | **Number of Values** | 3 | 3 | 3 | 4 | 2 | 3 |
| | **Mean** | 207.4 | 188.9 | 205 | 213.7 | 199.3 | 209.1 |
| | **Std. Deviation** | 9.61 | 28.88 | 26.95 | 13.12 | 4.53 | 6.4 |
| | **Std. Error of Mean** | 5.55 | 16.67 | 15.56 | 6.56 | 3.2 | 3.69 |
| | **Lower 95% CI of mean** | 183.5 | 117.2 | 138.1 | 192.8 | 158.6 | 193.2 |

version 3

| | | | | | | | |
|---|---|---|---|---|---|---|---|
| | **Upper 95% CI of mean** | 231.3 | 260.7 | 272 | 234.5 | 239.9 | 225 |
| **14** | **Number of Values** | 3 | 3 | 3 | 4 | 2 | 2 |
| | **Mean** | 211.6 | 201.7 | 241.6 | 222.7 | 228.3 | 213 |
| | **Std. Deviation** | 13.01 | 26.49 | 14.47 | 3.25 | 5.02 | 3.42 |
| | **Std. Error of Mean** | 7.51 | 15.29 | 8.36 | 1.62 | 3.55 | 2.42 |
| | **Lower 95% CI of mean** | 179.3 | 135.9 | 205.7 | 217.5 | 183.2 | 182.3 |
| | **Upper 95% CI of mean** | 243.9 | 267.5 | 277.6 | 227.9 | 273.4 | 243.7 |
| **16** | **Number of Values** | 3 | 3 | 1 | 2 | 0 | 2 |
| | **Mean** | 228.5 | 192.4 | 250 | 221.2 | | 216 |
| | **Std. Deviation** | 11.92 | 5.9 | 0 | 24.7 | | 13.77 |
| | **Std. Error of Mean** | 6.88 | 3.41 | 0 | 17.47 | | 9.73 |
| | **Lower 95% CI of mean** | 198.8 | 177.7 | | -0.68 | | 92.33 |
| | **Upper 95% CI of mean** | 258.1 | 207 | | 443.1 | | 339.7 |

version 3

**Supplementary Table 9:** Descriptive statistical summary of Randall–Selitto testing (hindpaw mechanical withdrawal thresholds) across all experimental groups in male rats, including No Intervention (Control; n = 6), HPI Only (Disease Control; n = 6), Sham FUS (n = 6), FUS Only (n = 6), FUS+HPI (Intervention; n = 4), and LA+HPI (Positive Control; n = 6) at each time point up to 16 weeks. Data are presented as mean, standard deviation (SD), standard error of the mean (SEM), and lower and upper 95% confidence intervals (CI). FUS = focused ultrasound; HPI = hindpaw incision; LA = local anesthetic.

| Time (Weeks) | Comparison | Summary | p Value |
|---|---|---|---|

| | | | |
|---|---|---|---|
| **-0.29** | **HPI Only vs Control** | ns | >.999 |
| | HPI Only vs Sham FUS | ns | >.999 |
| | HPI Only vs FUS Only | ns | >.999 |
| | HPI Only vs FUS+HPI | ns | >.999 |
| | HPI Only vs LA+HPI | ns | >.999 |
| **-0.14** | **HPI Only vs Control** | ns | >.999 |
| | HPI Only vs Sham FUS | ns | .654 |
| | HPI Only vs FUS Only | ns | .392 |
| | HPI Only vs FUS+HPI | ns | >.999 |
| | HPI Only vs LA+HPI | ns | .896 |
| **0** | **HPI Only vs Control** | *** | <.001 |
| | HPI Only vs Sham FUS | *** | <.001 |
| | HPI Only vs FUS Only | *** | <.001 |
| | HPI Only vs FUS+HPI | *** | <.001 |
| | HPI Only vs LA+HPI | *** | <.001 |
| **0.14** | **HPI Only vs Control** | *** | <.001 |
| | HPI Only vs Sham FUS | *** | <.001 |
| | HPI Only vs FUS Only | *** | <.001 |
| | HPI Only vs FUS+HPI | *** | <.001 |
| | HPI Only vs LA+HPI | ns | .059 |
| **0.43** | **HPI Only vs Control** | *** | <.001 |

version 3

| | | | |
|---|---|---|---|
| | HPI Only vs Sham FUS | *** | <.001 |
| | HPI Only vs FUS Only | *** | <.001 |
| | **HPI Only vs FUS+HPI** | *** | <.001 |
| | **HPI Only vs LA+HPI** | ns | >.999 |
| **1** | **HPI Only vs Control** | ** | .002 |
| | **HPI Only vs Sham FUS** | *** | <.001 |
| | **HPI Only vs FUS Only** | *** | <.001 |
| | **HPI Only vs FUS+HPI** | *** | <.001 |
| | **HPI Only vs LA+HPI** | ns | >.999 |
| **1.43** | **HPI Only vs Control** | ns | .144 |
| | **HPI Only vs Sham FUS** | ** | .007 |
| | **HPI Only vs FUS Only** | *** | <.001 |
| | **HPI Only vs FUS+HPI** | * | .018 |
| | **HPI Only vs LA+HPI** | ns | >.999 |
| **2** | **HPI Only vs Control** | ** | .005 |
| | **HPI Only vs Sham FUS** | *** | <.001 |
| | **HPI Only vs FUS Only** | *** | <.001 |
| | **HPI Only vs FUS+HPI** | ** | .003 |
| | **HPI Only vs LA+HPI** | ns | .735 |
| **2.43** | **HPI Only vs Control** | ns | .154 |
| | **HPI Only vs Sham FUS** | * | .015 |

version 3

| | | | |
|---|---|---|---|
| | **HPI Only vs FUS Only** | ** | .002 |
| | **HPI Only vs FUS+HPI** | ** | .009 |
| | **HPI Only vs LA+HPI** | ns | >.999 |
| **3** | **HPI Only vs Control** | *** | <.001 |
| | **HPI Only vs Sham FUS** | ** | .002 |
| | **HPI Only vs FUS Only** | *** | <.001 |
| | **HPI Only vs FUS+HPI** | *** | <.001 |
| | **HPI Only vs LA+HPI** | ns | .352 |
| **3.43** | **HPI Only vs Control** | * | .012 |
| | **HPI Only vs Sham FUS** | * | .012 |
| | **HPI Only vs FUS Only** | ** | .006 |
| | **HPI Only vs FUS+HPI** | ns | .067 |
| | **HPI Only vs LA+HPI** | ns | >.999 |
| **4** | **HPI Only vs Control** | * | .020 |
| | **HPI Only vs Sham FUS** | *** | <.001 |
| | **HPI Only vs FUS Only** | *** | <.001 |
| | **HPI Only vs FUS+HPI** | * | .014 |
| | **HPI Only vs LA+HPI** | ** | .008 |
| **5** | **HPI Only vs Control** | ns | .057 |
| | **HPI Only vs Sham FUS** | * | .013 |
| | **HPI Only vs FUS Only** | * | .028 |

version 3

| | | | |
|---|---|---|---|
| | **HPI Only vs FUS+HPI** | * | .014 |
| | **HPI Only vs LA+HPI** | ns | .794 |
| **6** | **HPI Only vs Control** | ns | .127 |
| | **HPI Only vs Sham FUS** | * | .028 |
| | **HPI Only vs FUS Only** | * | .044 |
| | **HPI Only vs FUS+HPI** | ns | .574 |
| | **HPI Only vs LA+HPI** | ns | .244 |
| **7** | **HPI Only vs Control** | ns | >.999 |
| | **HPI Only vs Sham FUS** | ns | >.999 |
| | **HPI Only vs FUS Only** | ns | >.999 |
| | **HPI Only vs FUS+HPI** | ns | >.999 |
| | **HPI Only vs LA+HPI** | ns | >.999 |
| **8** | **HPI Only vs Control** | ns | >.999 |
| | **HPI Only vs Sham FUS** | ns | >.999 |
| | **HPI Only vs FUS Only** | ns | >.999 |
| | **HPI Only vs FUS+HPI** | ns | >.999 |
| | **HPI Only vs LA+HPI** | ns | >.999 |
| **9** | **HPI Only vs Control** | ns | >.999 |
| | **HPI Only vs Sham FUS** | ns | >.999 |
| | **HPI Only vs FUS Only** | ns | .723 |
| | **HPI Only vs FUS+HPI** | ns | >.999 |

version 3

| | | | |
|---|---|---|---|
| | **HPI Only vs LA+HPI** | ns | >.999 |
| **12** | **HPI Only vs Control** | ns | >.999 |
| | **HPI Only vs Sham FUS** | ns | >.999 |
| | **HPI Only vs FUS Only** | ns | >.999 |
| | **HPI Only vs FUS+HPI** | ns | >.999 |
| | **HPI Only vs LA+HPI** | ns | >.999 |
| **14** | **HPI Only vs Control** | ns | >.999 |
| | **HPI Only vs Sham FUS** | ns | .176 |
| | **HPI Only vs FUS Only** | ns | >.999 |
| | **HPI Only vs FUS+HPI** | ns | >.999 |
| | **HPI Only vs LA+HPI** | ns | >.999 |
| **16** | **HPI Only vs Control** | ns | .472 |
| | **HPI Only vs Sham FUS** | ns | .972 |
| | **HPI Only vs FUS Only** | ns | >.999 |
| | **HPI Only vs FUS+HPI** | ns | >.999 |
| | **HPI Only vs LA+HPI** | ns | >.999 |

**Supplementary Table 10:** Summary of group-wise comparisons for Randall–Selitto testing (hindpaw mechanical withdrawal thresholds) across all experimental groups in male rats, including No Intervention (Control; n = 6), HPI Only (Disease Control; n = 6), Sham FUS (n = 6), FUS Only (n = 6), FUS+HPI (Intervention; n = 4), and LA+HPI (Positive Control; n = 6) at each time point up to 16 weeks. Measurements were analyzed using a two-way mixed-effects ANOVA with fixed effects for treatment group and time, followed by Bonferroni-corrected post hoc multiple-comparison testing. Statistical significance is denoted as follows: ns ($p \geq 0.05$), * ($p < 0.05$), ** ($p < 0.01$), *** ($p < 0.001$).

version 3

| Time (Weeks) | Study Arms | Control | HPI Only | Sham FUS | FUS Only | FUS+HPI | LA+HPI |
|---|---|---|---|---|---|---|---|
| **-0.29** | **Number of Values (N)** | 6 | 6 | 6 | 6 | 8 | 6 |
| | **Mean** | 182.9 | 208.4 | 210.8 | 217 | 209.4 | 214.9 |
| | **Std. Deviation** | 19.42 | 3 | 30.07 | 18.13 | 25.47 | 12.97 |
| | **Std. Error of Mean** | 7.93 | 1.22 | 12.28 | 7.4 | 9 | 5.3 |
| | **Lower 95% CI of mean** | 162.5 | 205.3 | 179.2 | 198 | 188.1 | 201.3 |
| | **Upper 95% CI of mean** | 203.3 | 211.5 | 242.3 | 236.1 | 230.7 | 228.5 |
| **-0.14** | **Number of Values** | 6 | 6 | 6 | 6 | 8 | 6 |
| | **Mean** | 229.1 | 212.9 | 213.7 | 216.7 | 187.7 | 205.3 |
| | **Std. Deviation** | 8.66 | 8.09 | 23.06 | 20.88 | 25.87 | 22.18 |
| | **Std. Error of Mean** | 3.54 | 3.3 | 9.41 | 8.53 | 9.15 | 9.06 |
| | **Lower 95% CI of mean** | 220 | 204.4 | 189.5 | 194.8 | 166.1 | 182 |
| | **Upper 95% CI of mean** | 238.1 | 221.3 | 237.9 | 238.6 | 209.4 | 228.5 |
| **0** | **Number of Values** | 6 | 6 | 6 | 6 | 8 | 6 |
| | **Mean** | 222.9 | 43.72 | 223.6 | 232.6 | 215.6 | 225.3 |
| | **Std. Deviation** | 18.15 | 12.31 | 18.14 | 10.8 | 33.53 | 13.77 |

version 3

| | | | | | | | |
|---|---|---|---|---|---|---|---|
| | **Std. Error of Mean** | 7.41 | 5.03 | 7.41 | 4.41 | 11.85 | 5.62 |
| | **Lower 95% CI of mean** | 203.8 | 30.8 | 204.6 | 221.3 | 187.6 | 210.8 |
| | **Upper 95% CI of mean** | 241.9 | 56.64 | 242.7 | 244 | 243.6 | 239.7 |
| **0.14** | **Number of Values** | 6 | 6 | 6 | 6 | 8 | 6 |
| | **Mean** | 209.8 | 54.93 | 221.1 | 235.4 | 221.6 | 82.85 |
| | **Std. Deviation** | 9.84 | 14.51 | 19.46 | 14.63 | 31.03 | 10.59 |
| | **Std. Error of Mean** | 4.02 | 5.92 | 7.95 | 5.97 | 10.97 | 4.32 |
| | **Lower 95% CI of mean** | 199.5 | 39.7 | 200.7 | 220.1 | 195.7 | 71.74 |
| | **Upper 95% CI of mean** | 220.1 | 70.15 | 241.6 | 250.8 | 247.5 | 93.96 |
| **0.43** | **Number of Values** | 6 | 6 | 6 | 6 | 8 | 6 |
| | **Mean** | 211.8 | 129.9 | 218 | 228.8 | 218.8 | 112.5 |
| | **Std. Deviation** | 23.36 | 30.88 | 15.42 | 19.5 | 27.54 | 63.27 |
| | **Std. Error of Mean** | 9.54 | 12.61 | 6.3 | 7.96 | 9.74 | 25.83 |
| | **Lower 95% CI of mean** | 187.3 | 97.45 | 201.8 | 208.4 | 195.8 | 46.14 |
| | **Upper 95% CI of mean** | 236.3 | 162.3 | 234.2 | 249.3 | 241.9 | 178.9 |
| **1** | **Number of Values** | 6 | 6 | 6 | 6 | 8 | 6 |

version 3

| | | | | | | | |
|---|---|---|---|---|---|---|---|
| | **Mean** | 190.4 | 154.2 | 212.4 | 213.7 | 213.3 | 126.7 |
| | **Std. Deviation** | 14.21 | 31.47 | 10.3 | 16.96 | 28.63 | 5.14 |
| | **Std. Error of Mean** | 5.8 | 12.85 | 4.21 | 6.92 | 10.12 | 2.1 |
| | **Lower 95% CI of mean** | 175.5 | 121.1 | 201.6 | 195.9 | 189.4 | 121.3 |
| | **Upper 95% CI of mean** | 205.3 | 187.2 | 223.3 | 231.4 | 237.2 | 132.1 |
| **1.43** | **Number of Values** | 4 | 6 | 6 | 6 | 8 | 4 |
| | **Mean** | 198 | 167.2 | 220.6 | 207.4 | 219.5 | 143.6 |
| | **Std. Deviation** | 16.61 | 35.85 | 6.45 | 21.51 | 27.25 | 32.75 |
| | **Std. Error of Mean** | 8.31 | 14.64 | 2.63 | 8.78 | 9.63 | 16.38 |
| | **Lower 95% CI of mean** | 171.5 | 129.6 | 213.8 | 184.9 | 196.7 | 91.52 |
| | **Upper 95% CI of mean** | 224.4 | 204.8 | 227.4 | 230 | 242.3 | 195.8 |
| **2** | **Number of Values** | 4 | 6 | 6 | 4 | 8 | 4 |
| | **Mean** | 232.3 | 155.9 | 203.3 | 214 | 209.2 | 143.9 |
| | **Std. Deviation** | 13.67 | 15.4 | 35.84 | 29.32 | 32.7 | 23.87 |
| | **Std. Error of Mean** | 6.84 | 6.29 | 14.63 | 14.66 | 11.56 | 11.93 |
| | **Lower 95% CI of mean** | 210.6 | 139.7 | 165.7 | 167.3 | 181.9 | 105.9 |

version 3

| | | | | | | | |
|---|---|---|---|---|---|---|---|
| | **Upper 95% CI of mean** | 254.1 | 172 | 240.9 | 260.6 | 236.6 | 181.9 |
| **2.43** | **Number of Values** | 4 | 6 | 4 | 4 | 7 | 4 |
| | **Mean** | 215.5 | 138.3 | 212.5 | 225.1 | 230.9 | 163.9 |
| | **Std. Deviation** | 21.9 | 14 | 5.7 | 4.49 | 15.58 | 41.23 |
| | **Std. Error of Mean** | 10.95 | 5.72 | 2.85 | 2.25 | 5.89 | 20.61 |
| | **Lower 95% CI of mean** | 180.7 | 123.6 | 203.5 | 217.9 | 216.5 | 98.3 |
| | **Upper 95% CI of mean** | 250.4 | 153 | 221.6 | 232.2 | 245.3 | 229.5 |
| **3** | **Number of Values** | 4 | 6 | 4 | 4 | 7 | 4 |
| | **Mean** | 224.6 | 167.4 | 196.9 | 229.1 | 219.7 | 183.9 |
| | **Std. Deviation** | 9.91 | 12.78 | 21.78 | 6.85 | 22.37 | 25.69 |
| | **Std. Error of Mean** | 4.95 | 5.22 | 10.89 | 3.42 | 8.46 | 12.85 |
| | **Lower 95% CI of mean** | 208.8 | 154 | 162.3 | 218.2 | 199 | 143 |
| | **Upper 95% CI of mean** | 240.3 | 180.8 | 231.6 | 240 | 240.4 | 224.8 |
| **3.43** | **Number of Values** | 4 | 6 | 3 | 4 | 6 | 4 |
| | **Mean** | 223.6 | 158.7 | 205 | 227.3 | 217.3 | 196.6 |
| | **Std. Deviation** | 13.71 | 12.34 | 21.65 | 15.01 | 23.5 | 27.04 |

version 3

| | | | | | | | |
|---|---|---|---|---|---|---|---|
| | **Std. Error of Mean** | 6.86 | 5.04 | 12.5 | 7.51 | 9.59 | 13.52 |
| | **Lower 95% CI of mean** | 201.8 | 145.8 | 151.3 | 203.4 | 192.6 | 153.6 |
| | **Upper 95% CI of mean** | 245.5 | 171.7 | 258.8 | 251.2 | 241.9 | 239.7 |
| **4** | **Number of Values** | 3 | 5 | 3 | 4 | 5 | 3 |
| | **Mean** | 203.2 | 172.9 | 195.5 | 210.3 | 213.6 | 193 |
| | **Std. Deviation** | 5.75 | 30.08 | 10.83 | 23.07 | 19.81 | 19.85 |
| | **Std. Error of Mean** | 3.32 | 13.45 | 6.25 | 11.53 | 8.86 | 11.46 |
| | **Lower 95% CI of mean** | 188.9 | 135.5 | 168.6 | 173.6 | 189 | 143.7 |
| | **Upper 95% CI of mean** | 217.5 | 210.2 | 222.4 | 247 | 238.2 | 242.3 |
| **5** | **Number of Values** | 3 | 5 | 3 | 4 | 5 | 3 |
| | **Mean** | 209.8 | 193.7 | 198.4 | 211.2 | 208.7 | 188.8 |
| | **Std. Deviation** | 14.12 | 38.72 | 30.02 | 6.68 | 18.12 | 13.56 |
| | **Std. Error of Mean** | 8.15 | 17.32 | 17.33 | 3.34 | 8.1 | 7.83 |
| | **Lower 95% CI of mean** | 174.7 | 145.6 | 123.8 | 200.6 | 186.2 | 155.1 |
| | **Upper 95% CI of mean** | 244.9 | 241.7 | 273 | 221.8 | 231.2 | 222.5 |
| **6** | **Number of Values** | 3 | 5 | 3 | 4 | 5 | 3 |

| | | | | | | | |
|---|---|---|---|---|---|---|---|
| | **Mean** | 206.8 | 201.3 | 190.6 | 220.9 | 199.4 | 194.2 |
| | **Std. Deviation** | 14.1 | 29.63 | 17.37 | 5.81 | 30.87 | 4.71 |
| | **Std. Error of Mean** | 8.14 | 13.25 | 10.03 | 2.9 | 13.8 | 2.72 |
| | **Lower 95% CI of mean** | 171.8 | 164.5 | 147.5 | 211.6 | 161.1 | 182.5 |
| | **Upper 95% CI of mean** | 241.8 | 238.1 | 233.8 | 230.1 | 237.8 | 205.9 |
| **7** | **Number of Values** | 3 | 5 | 3 | 3 | 5 | 3 |
| | **Mean** | 214.4 | 195.5 | 221.7 | 207.5 | 212.3 | 199.2 |
| | **Std. Deviation** | 11.21 | 31.58 | 11.59 | 15.1 | 18.83 | 24.38 |
| | **Std. Error of Mean** | 6.47 | 14.12 | 6.69 | 8.72 | 8.42 | 14.08 |
| | **Lower 95% CI of mean** | 186.6 | 156.3 | 192.9 | 170 | 188.9 | 138.6 |
| | **Upper 95% CI of mean** | 242.2 | 234.7 | 250.5 | 245 | 235.7 | 259.8 |
| **8** | **Number of Values** | 3 | 5 | 3 | 3 | 5 | 3 |
| | **Mean** | 217.9 | 200.2 | 211.8 | 215.5 | 216.6 | 211.9 |
| | **Std. Deviation** | 4.73 | 30.45 | 11.42 | 7.82 | 16.17 | 20 |
| | **Std. Error of Mean** | 2.73 | 13.62 | 6.6 | 4.52 | 7.23 | 11.55 |
| | **Lower 95% CI of mean** | 206.1 | 162.4 | 183.4 | 196 | 196.5 | 162.2 |

version 3

| | | | | | | | |
|---|---|---|---|---|---|---|---|
| | **Upper 95% CI of mean** | 229.6 | 238 | 240.2 | 234.9 | 236.7 | 261.6 |
| **9** | **Number of Values** | 3 | 3 | 3 | 2 | 4 | 3 |
| | **Mean** | 207.1 | 189 | 221.8 | 203.5 | 207.1 | 212.6 |
| | **Std. Deviation** | 33.58 | 9.2 | 3.69 | 1.95 | 16.47 | 20.96 |
| | **Std. Error of Mean** | 19.39 | 5.31 | 2.13 | 1.38 | 8.23 | 12.1 |
| | **Lower 95% CI of mean** | 123.7 | 166.2 | 212.6 | 185.9 | 180.9 | 160.5 |
| | **Upper 95% CI of mean** | 290.5 | 211.9 | 231 | 221 | 233.3 | 264.7 |
| **12** | **Number of Values** | 3 | 3 | 3 | 2 | 3 | 3 |
| | **Mean** | 195.8 | 176.7 | 200.2 | 199.5 | 199.3 | 206.8 |
| | **Std. Deviation** | 12.76 | 1.24 | 6.6 | 8.66 | 12.12 | 7.45 |
| | **Std. Error of Mean** | 7.37 | 0.71 | 3.81 | 6.12 | 7 | 4.3 |
| | **Lower 95% CI of mean** | 164.1 | 173.7 | 183.9 | 121.7 | 169.1 | 188.3 |
| | **Upper 95% CI of mean** | 227.5 | 179.8 | 216.6 | 277.2 | 229.4 | 225.3 |
| **14** | **Number of Values** | 2 | 1 | 3 | 2 | 2 | 3 |
| | **Mean** | 208.4 | 196.1 | 212.3 | 208.3 | 218.4 | 209.9 |
| | **Std. Deviation** | 0.12 | 0 | 22.57 | 7.14 | 19.49 | 35.52 |

version 3

| | | | | | | | |
|---|---|---|---|---|---|---|---|
| | **Std. Error of Mean** | 0.08 | 0 | 13.03 | 5.05 | 13.79 | 20.51 |
| | **Lower 95% CI of mean** | 207.3 | | 156.3 | 144.2 | 43.26 | 121.7 |
| | **Upper 95% CI of mean** | 209.4 | | 268.4 | 272.5 | 393.6 | 298.2 |
| **16** | **Number of Values** | 1 | 1 | 3 | 1 | 2 | 2 |
| | **Mean** | 193.3 | 208.5 | 207.5 | 226 | 206.9 | 221.8 |
| | **Std. Deviation** | 0 | 0 | 19.43 | 0 | 7.59 | 25.76 |
| | **Std. Error of Mean** | 0 | 0 | 11.22 | 0 | 5.37 | 18.22 |
| | **Lower 95% CI of mean** | | | 159.3 | | 138.8 | -9.65 |
| | **Upper 95% CI of mean** | | | 255.8 | | 275.1 | 453.3 |

version 3

**Supplementary Table 11:** Descriptive statistical summary of Randall–Selitto testing (hindpaw mechanical withdrawal thresholds) across all experimental groups in female rats, including No Intervention (Control; n = 6), HPI Only (Disease Control; n = 6), Sham FUS (n = 6), FUS Only (n = 6), FUS+HPI (Intervention; n = 8), and LA+HPI (Positive Control; n = 6) at each time point up to 16 weeks. Data are presented as mean, standard deviation (SD), standard error of the mean (SEM), and lower and upper 95% confidence intervals (CI). FUS = focused ultrasound; HPI = hindpaw incision; LA = local anesthetic.

| Time (Weeks) | Comparison | Summary | p Value |
|---|---|---|---|
| **-0.29** | **HPI Only vs Control** | ns | .481 |
| | **HPI Only vs Sham FUS** | ns | >.999 |
| | **HPI Only vs FUS Only** | ns | >.999 |
| | **HPI Only vs FUS+HPI** | ns | >.999 |

| | | | |
|---|---|---|---|
| | **HPI Only vs LA+HPI** | ns | >.999 |
| **-0.14** | **HPI Only vs Control** | ns | >.999 |
| | **HPI Only vs Sham FUS** | ns | >.999 |
| | **HPI Only vs FUS Only** | ns | >.999 |
| | **HPI Only vs FUS+HPI** | ns | .210 |
| | **HPI Only vs LA+HPI** | ns | >.999 |
| **0** | **HPI Only vs Control** | *** | <.001 |
| | **HPI Only vs Sham FUS** | *** | <.001 |
| | **HPI Only vs FUS Only** | *** | <.001 |
| | **HPI Only vs FUS+HPI** | *** | <.001 |
| | **HPI Only vs LA+HPI** | *** | <.001 |
| **0.14** | **HPI Only vs Control** | *** | <.001 |
| | **HPI Only vs Sham FUS** | *** | <.001 |
| | **HPI Only vs FUS Only** | *** | <.001 |
| | **HPI Only vs FUS+HPI** | *** | <.001 |
| | **HPI Only vs LA+HPI** | ns | .242 |
| **0.43** | **HPI Only vs Control** | ** | .003 |
| | **HPI Only vs Sham FUS** | ** | .001 |
| | **HPI Only vs FUS Only** | *** | <.001 |
| | **HPI Only vs FUS+HPI** | ** | .004 |
| | **HPI Only vs LA+HPI** | ns | >.999 |

version 3

| | | | |
|---|---|---|---|
| **1** | **HPI Only vs Control** | * | .026 |
| | **HPI Only vs Sham FUS** | *** | <.001 |
| | **HPI Only vs FUS Only** | *** | <.001 |
| | **HPI Only vs FUS+HPI** | ** | .001 |
| | **HPI Only vs LA+HPI** | ns | .204 |
| **1.43** | **HPI Only vs Control** | ns | .239 |
| | **HPI Only vs Sham FUS** | ** | .008 |
| | **HPI Only vs FUS Only** | ns | .089 |
| | **HPI Only vs FUS+HPI** | * | .040 |
| | **HPI Only vs LA+HPI** | ns | >.999 |
| **2** | **HPI Only vs Control** | *** | <.001 |
| | **HPI Only vs Sham FUS** | * | .016 |
| | **HPI Only vs FUS Only** | * | .038 |
| | **HPI Only vs FUS+HPI** | * | .023 |
| | **HPI Only vs LA+HPI** | ns | >.999 |
| **2.43** | **HPI Only vs Control** | *** | <.001 |
| | **HPI Only vs Sham FUS** | ** | .001 |
| | **HPI Only vs FUS Only** | *** | <.001 |
| | **HPI Only vs FUS+HPI** | *** | <.001 |
| | **HPI Only vs LA+HPI** | ns | >.999 |
| **3** | **HPI Only vs Control** | *** | <.001 |

version 3

| | | | |
|---|---|---|---|
| | **HPI Only vs Sham FUS** | ns | .139 |
| | **HPI Only vs FUS Only** | ** | .002 |
| | **HPI Only vs FUS+HPI** | ** | .003 |
| | **HPI Only vs LA+HPI** | ns | >.999 |
| **3.43** | **HPI Only vs Control** | ** | .005 |
| | **HPI Only vs Sham FUS** | ns | .086 |
| | **HPI Only vs FUS Only** | ** | .005 |
| | **HPI Only vs FUS+HPI** | ** | .010 |
| | **HPI Only vs LA+HPI** | ns | .222 |
| **4** | **HPI Only vs Control** | ns | .518 |
| | **HPI Only vs Sham FUS** | ns | >.999 |
| | **HPI Only vs FUS Only** | ns | >.999 |
| | **HPI Only vs FUS+HPI** | ns | .188 |
| | **HPI Only vs LA+HPI** | ns | >.999 |
| **5** | **HPI Only vs Control** | ns | >.999 |
| | **HPI Only vs Sham FUS** | ns | >.999 |
| | **HPI Only vs FUS Only** | ns | >.999 |
| | **HPI Only vs FUS+HPI** | ns | >.999 |
| | **HPI Only vs LA+HPI** | ns | >.999 |
| **6** | **HPI Only vs Control** | ns | >.999 |
| | **HPI Only vs Sham FUS** | ns | >.999 |

version 3

| | | | |
|---|---|---|---|
| | **HPI Only vs FUS Only** | ns | >.999 |
| | **HPI Only vs FUS+HPI** | ns | >.999 |
| | **HPI Only vs LA+HPI** | ns | >.999 |
| **7** | **HPI Only vs Control** | ns | >.999 |
| | **HPI Only vs Sham FUS** | ns | >.999 |
| | **HPI Only vs FUS Only** | ns | >.999 |
| | **HPI Only vs FUS+HPI** | ns | >.999 |
| | **HPI Only vs LA+HPI** | ns | >.999 |
| **8** | **HPI Only vs Control** | ns | >.999 |
| | **HPI Only vs Sham FUS** | ns | >.999 |
| | **HPI Only vs FUS Only** | ns | >.999 |
| | **HPI Only vs FUS+HPI** | ns | >.999 |
| | **HPI Only vs LA+HPI** | ns | >.999 |
| **9** | **HPI Only vs Control** | ns | >.999 |
| | **HPI Only vs Sham FUS** | ns | .858 |
| | **HPI Only vs FUS Only** | ns | >.999 |
| | **HPI Only vs FUS+HPI** | ns | >.999 |
| | **HPI Only vs LA+HPI** | ns | >.999 |
| **12** | **HPI Only vs Control** | ns | .683 |
| | **HPI Only vs Sham FUS** | ns | .288 |
| | **HPI Only vs FUS Only** | ns | .542 |

version 3

| | | | |
|---|---|---|---|
| | **HPI Only vs FUS+HPI** | ns | .348 |
| | **HPI Only vs LA+HPI** | ns | .086 |
| **14** | **HPI Only vs Control** | ns | >.999 |
| | **HPI Only vs Sham FUS** | ns | >.999 |
| | **HPI Only vs FUS Only** | ns | >.999 |
| | **HPI Only vs FUS+HPI** | ns | >.999 |
| | **HPI Only vs LA+HPI** | ns | >.999 |
| **16** | **HPI Only vs Control** | ns | >.999 |
| | **HPI Only vs Sham FUS** | ns | >.999 |
| | **HPI Only vs FUS Only** | ns | >.999 |
| | **HPI Only vs FUS+HPI** | ns | >.999 |
| | **HPI Only vs LA+HPI** | ns | >.999 |

version 3

**Supplementary Table 12:** Summary of group-wise comparisons for Randall–Selitto testing (hindpaw mechanical withdrawal thresholds) across all experimental groups in female rats, including No Intervention (Control; n = 6), HPI Only (Disease Control; n = 6), Sham FUS (n = 6), FUS Only (n = 6), FUS+HPI (Intervention; n = 8), and LA+HPI (Positive Control; n = 6) at each time point up to 16 weeks. Measurements were analyzed using a two-way mixed-effects ANOVA with fixed effects for treatment group and time, followed by Bonferroni-corrected post hoc multiple-comparison testing. Statistical significance is denoted as follows: ns ($p \geq 0.05$), * ($p < 0.05$), ** ($p < 0.01$), *** ($p < 0.001$).

| **Time (Weeks)** | **Study Arms** | **No Response N (%)** | **Partial Flexion N (%)** | **Full Flexion N (%)** | **Summary** | **Fisher's exact p-value** |
|---|---|---|---|---|---|---|
| **-0.29** | **Control** | 0 | 0 | 12 (100%) | ns | >0.9999 |
| | **HPI Only** | 0 | 0 | 12 (100%) | | |

| | | | | | | |
|---|---|---|---|---|---|---|
| | **Sham FUS** | 0 | 0 | 12 (100%) | | |
| | **FUS Only** | 0 | 0 | 12 (100%) | | |
| | **FUS+HPI** | 0 | 0 | 12 (100%) | | |
| | **LA+HPI** | 0 | 0 | 12 (100%) | | |
| **-0.14** | **Control** | 0 | 0 | 12 (100%) | ns | >0.9999 |
| | **HPI Only** | 0 | 0 | 12 (100%) | | |
| | **Sham FUS** | 0 | 0 | 12 (100%) | | |
| | **FUS Only** | 0 | 0 | 12 (100%) | | |
| | **FUS+HPI** | 0 | 0 | 12 (100%) | | |
| | **LA+HPI** | 0 | 0 | 12 (100%) | | |
| **0** | **Control** | 0 | 0 | 12 (100%) | **** | <0.0001 |
| | **HPI Only** | 0 | 0 | 12 (100%) | | |
| | **Sham FUS** | 0 | 0 | 12 (100%) | | |
| | **FUS Only** | 0 | 1 (8.33%) | 11 (91.67%) | | |
| | **FUS+HPI** | 0 | 0 | 12 (100%) | | |
| | **LA+HPI** | 7 (58.33%) | 5 (41.67%) | 0 | | |
| **0.14** | **Control** | 0 | 0 | 12 (100%) | ns | >0.9999 |

version 3

| | | | | | | |
|---|---|---|---|---|---|---|
| | **HPI Only** | 0 | 0 | 12 (100%) | | |
| | **Sham FUS** | 0 | 0 | 12 (100%) | | |
| | **FUS Only** | 0 | 1 (8.33%) | 11 (91.67%) | | |
| | **FUS+HPI** | 0 | 0 | 12 (100%) | | |
| | **LA+HPI** | 0 | 0 | 12 (100%) | | |
| **0.43** | **Control** | 0 | 0 | 12 (100%) | ns | >0.9999 |
| | **HPI Only** | 0 | 0 | 12 (100%) | | |
| | **Sham FUS** | 0 | 0 | 12 (100%) | | |
| | **FUS Only** | 0 | 1 (8.33%) | 11 (91.67%) | | |
| | **FUS+HPI** | 0 | 0 | 12 (100%) | | |
| | **LA+HPI** | 0 | 0 | 12 (100%) | | |
| **1** | **Control** | 0 | 0 | 12 (100%) | ns | >0.9999 |
| | **HPI Only** | 0 | 0 | 12 (100%) | | |
| | **Sham FUS** | 0 | 0 | 12 (100%) | | |
| | **FUS Only** | 0 | 1 (8.33%) | 11 (91.67%) | | |
| | **FUS+HPI** | 0 | 0 | 12 (100%) | | |
| | **LA+HPI** | 0 | 0 | 12 (100%) | | |

version 3

| | | | | | | |
|---|---|---|---|---|---|---|
| **1.43** | **Control** | 0 | 0 | 12 (100%) | ns | >0.9999 |
| | **HPI Only** | 0 | 0 | 12 (100%) | | |
| | **Sham FUS** | 0 | 0 | 12 (100%) | | |
| | **FUS Only** | 0 | 0 | 12 (100%) | | |
| | **FUS+HPI** | 0 | 0 | 12 (100%) | | |
| | **LA+HPI** | 0 | 0 | 12 (100%) | | |
| **2** | **Control** | 0 | 0 | 12 (100%) | ns | >0.9999 |
| | **HPI Only** | 0 | 0 | 12 (100%) | | |
| | **Sham FUS** | 0 | 0 | 12 (100%) | | |
| | **FUS Only** | 0 | 0 | 12 (100%) | | |
| | **FUS+HPI** | 0 | 0 | 12 (100%) | | |
| | **LA+HPI** | 0 | 0 | 12 (100%) | | |
| **2.43** | **Control** | 0 | 0 | 10 (100%) | ns | >0.9999 |
| | **HPI Only** | 0 | 0 | 10 (100%) | | |
| | **Sham FUS** | 0 | 0 | 10 (100%) | | |
| | **FUS Only** | 0 | 0 | 10 (100%) | | |
| | **FUS+HPI** | 0 | 0 | 10 (100%) | | |

version 3

| | | | | | | |
|---|---|---|---|---|---|---|
| | **LA+HPI** | 0 | 0 | 10 (100%) | | |
| **3** | **Control** | 0 | 0 | 10 (100%) | ns | >0.9999 |
| | **HPI Only** | 0 | 0 | 10 (100%) | | |
| | **Sham FUS** | 0 | 0 | 10 (100%) | | |
| | **FUS Only** | 0 | 0 | 10 (100%) | | |
| | **FUS+HPI** | 0 | 0 | 10 (100%) | | |
| | **LA+HPI** | 0 | 0 | 10 (100%) | | |
| **3.43** | **Control** | 0 | 0 | 10 (100%) | ns | >0.9999 |
| | **HPI Only** | 0 | 0 | 10 (100%) | | |
| | **Sham FUS** | 0 | 0 | 10 (100%) | | |
| | **FUS Only** | 0 | 0 | 10 (100%) | | |
| | **FUS+HPI** | 0 | 0 | 10 (100%) | | |
| | **LA+HPI** | 0 | 0 | 10 (100%) | | |
| **4** | **Control** | 0 | 0 | 8 (100%) | ns | >0.9999 |
| | **HPI Only** | 0 | 0 | 8 (100%) | | |
| | **Sham FUS** | 0 | 0 | 10 (100%) | | |
| | **FUS Only** | 0 | 0 | 10 (100%) | | |

version 3

| | | | | | | |
|---|---|---|---|---|---|---|
| | **FUS+HPI** | 0 | 0 | 10 (100%) | | |
| | **LA+HPI** | 0 | 0 | 8 (100%) | | |
| **5** | **Control** | 0 | 0 | 8 (100%) | ns | >0.9999 |
| | **HPI Only** | 0 | 0 | 8 (100%) | | |
| | **Sham FUS** | 0 | 0 | 8 (100%) | | |
| | **FUS Only** | 0 | 0 | 8 (100%) | | |
| | **FUS+HPI** | 0 | 0 | 8 (100%) | | |
| | **LA+HPI** | 0 | 0 | 8 (100%) | | |
| **6** | **Control** | 0 | 0 | 8 (100%) | ns | >0.9999 |
| | **HPI Only** | 0 | 0 | 8 (100%) | | |
| | **Sham FUS** | 0 | 0 | 8 (100%) | | |
| | **FUS Only** | 0 | 0 | 8 (100%) | | |
| | **FUS+HPI** | 0 | 0 | 8 (100%) | | |
| | **LA+HPI** | 0 | 0 | 8 (100%) | | |
| **7** | **Control** | 0 | 0 | 8 (100%) | ns | >0.9999 |
| | **HPI Only** | 0 | 0 | 8 (100%) | | |
| | **Sham FUS** | 0 | 0 | 8 (100%) | | |

version 3

| | | | | | | |
|---|---|---|---|---|---|---|
| | **FUS Only** | 0 | 0 | 8 (100%) | | |
| | **FUS+HPI** | 0 | 0 | 8 (100%) | | |
| | **LA+HPI** | 0 | 0 | 8 (100%) | | |
| **8** | **Control** | 0 | 0 | 6 (100%) | ns | >0.9999 |
| | **HPI Only** | 0 | 0 | 6 (100%) | | |
| | **Sham FUS** | 0 | 0 | 8 (100%) | | |
| | **FUS Only** | 0 | 0 | 8 (100%) | | |
| | **FUS+HPI** | 0 | 0 | 8 (100%) | | |
| | **LA+HPI** | 0 | 0 | 8 (100%) | | |
| **9** | **Control** | 0 | 0 | 6 (100%) | ns | >0.9999 |
| | **HPI Only** | 0 | 0 | 6 (100%) | | |
| | **Sham FUS** | 0 | 0 | 6 (100%) | | |
| | **FUS Only** | 0 | 0 | 6 (100%) | | |
| | **FUS+HPI** | 0 | 0 | 6 (100%) | | |
| | **LA+HPI** | 0 | 0 | 8 (100%) | | |
| **12** | **Control** | 0 | 0 | 6 (100%) | ns | >0.9999 |
| | **HPI Only** | 0 | 0 | 6 (100%) | | |

| | | | | | | |
|---|---|---|---|---|---|---|
| | **Sham FUS** | 0 | 0 | 6 (100%) | | |
| | **FUS Only** | 0 | 0 | 6 (100%) | | |
| | **FUS+HPI** | 0 | 0 | 6 (100%) | | |
| | **LA+HPI** | 0 | 0 | 8 (100%) | | |
| **14** | **Control** | 0 | 0 | 6 (100%) | ns | >0.9999 |
| | **HPI Only** | 0 | 0 | 4 (100%) | | |
| | **Sham FUS** | 0 | 0 | 6 (100%) | | |
| | **FUS Only** | 0 | 0 | 6 (100%) | | |
| | **FUS+HPI** | 0 | 0 | 5 (100%) | | |
| | **LA+HPI** | 0 | 0 | 5 (100%) | | |
| **16** | **Control** | 0 | 0 | 6 (100%) | ns | >0.9999 |
| | **HPI Only** | 0 | 0 | 4 (100%) | | |
| | **Sham FUS** | 0 | 0 | 4 (100%) | | |
| | **FUS Only** | 0 | 0 | 4 (100%) | | |
| | **FUS+HPI** | 0 | 0 | 4 (100%) | | |
| | **LA+HPI** | 0 | 0 | 5 (100%) | | |

version 3

**Supplementary Table 13:** Hindpaw (HP) flexion testing (motor and non-pain sensory response) across all experimental groups, including No Intervention (Control; n = 12), HPI Only (Disease

Control; n = 12), Sham FUS (n = 12), FUS Only (n = 12), FUS+HPI (Intervention; n = 12), and LA+HPI (Positive Control; n = 12) at each time point up to 16 weeks. Data were analyzed using contingency table analysis followed by Fisher's exact test, and behavioral responses are presented as fractional response values. FUS = focused ultrasound; HPI = hindpaw incision; LA = local anesthetic.

| **Time (Weeks)** | **Study Arms** | **No Response N (%)** | **Partial Flexion N (%)** | **Full Flexion N (%)** | **Summary** | **Fisher's exact p-value** |
|---|---|---|---|---|---|---|
| **-0.29** | **Control** | 0 | 0 | 6 (100%) | ns | >0.9999 |
| | **HPI Only** | 0 | 0 | 6 (100%) | | |
| | **Sham FUS** | 0 | 0 | 6 (100%) | | |
| | **FUS Only** | 0 | 0 | 6 (100%) | | |
| | **FUS+HPI** | 0 | 0 | 4 (100%) | | |
| | **LA+HPI** | 0 | 0 | 6 (100%) | | |
| **-0.14** | **Control** | 0 | 0 | 6 (100%) | ns | >0.9999 |
| | **HPI Only** | 0 | 0 | 6 (100%) | | |
| | **Sham FUS** | 0 | 0 | 6 (100%) | | |
| | **FUS Only** | 0 | 0 | 6 (100%) | | |
| | **FUS+HPI** | 0 | 0 | 4 (100%) | | |
| | **LA+HPI** | 0 | 0 | 6 (100%) | | |
| **0** | **Control** | 0 | 0 | 6 (100%) | **** | <0.0001 |
| | **HPI Only** | 0 | 0 | 6 (100%) | | |

version 3

| | | | | | | |
|---|---|---|---|---|---|---|
| | **Sham FUS** | 0 | 0 | 6 (100%) | | |
| | **FUS Only** | 1 (16.67%) | 1 (16.67%) | 4 (66.7%) | | |
| | **FUS+HPI** | 0 | 0 | 4 (100%) | | |
| | **LA+HPI** | 5 (83.33%) | 1 (16.67%) | 0 | | |
| **0.14** | **Control** | 0 | 0 | 6 (100%) | ns | 0.3583 |
| | **HPI Only** | 0 | 0 | 6 (100%) | | |
| | **Sham FUS** | 0 | 0 | 6 (100%) | | |
| | **FUS Only** | 1 (16.67%) | 1 (16.67%) | 4 (66.7%) | | |
| | **FUS+HPI** | 0 | 0 | 4 (100%) | | |
| | **LA+HPI** | 0 | 0 | 6 (100%) | | |
| **0.43** | **Control** | 0 | 0 | 6 (100%) | ns | 0.3583 |
| | **HPI Only** | 0 | 0 | 6 (100%) | | |
| | **Sham FUS** | 0 | 0 | 6 (100%) | | |
| | **FUS Only** | 1 (16.67%) | 1 (16.67%) | 4 (66.7%) | | |
| | **FUS+HPI** | 0 | 0 | 4 (100%) | | |
| | **LA+HPI** | 0 | 0 | 6 (100%) | | |
| **1** | **Control** | 0 | 0 | 6 (100%) | ns | >0.9999 |

version 3

| | | | | | | |
|---|---|---|---|---|---|---|
| | **HPI Only** | 0 | 0 | 6 (100%) | | |
| | **Sham FUS** | 0 | 0 | 6 (100%) | | |
| | **FUS Only** | 0 | 1 (16.67%) | 5 (83.33%) | | |
| | **FUS+HPI** | 0 | 0 | 4 (100%) | | |
| | **LA+HPI** | 0 | 0 | 6 (100%) | | |
| **1.43** | **Control** | 0 | 0 | 6 (100%) | ns | >0.9999 |
| | **HPI Only** | 0 | 0 | 6 (100%) | | |
| | **Sham FUS** | 0 | 0 | 6 (100%) | | |
| | **FUS Only** | 0 | 0 | 6 (100%) | | |
| | **FUS+HPI** | 0 | 0 | 4 (100%) | | |
| | **LA+HPI** | 0 | 0 | 6 (100%) | | |
| **2** | **Control** | 0 | 0 | 6 (100%) | ns | >0.9999 |
| | **HPI Only** | 0 | 0 | 4 (100%) | | |
| | **Sham FUS** | 0 | 0 | 6 (100%) | | |
| | **FUS Only** | 0 | 0 | 6 (100%) | | |
| | **FUS+HPI** | 0 | 0 | 4 (100%) | | |
| | **LA+HPI** | 0 | 0 | 6 (100%) | | |

version 3

| | | | | | | |
|---|---|---|---|---|---|---|
| **2.43** | **Control** | 0 | 0 | 6 (100%) | ns | >0.9999 |
| | **HPI Only** | 0 | 0 | 4 (100%) | | |
| | **Sham FUS** | 0 | 0 | 6 (100%) | | |
| | **FUS Only** | 0 | 0 | 6 (100%) | | |
| | **FUS+HPI** | 0 | 0 | 3 (100%) | | |
| | **LA+HPI** | 0 | 0 | 6 (100%) | | |
| **3** | **Control** | 0 | 0 | 6 (100%) | ns | >0.9999 |
| | **HPI Only** | 0 | 0 | 4 (100%) | | |
| | **Sham FUS** | 0 | 0 | 6 (100%) | | |
| | **FUS Only** | 0 | 0 | 6 (100%) | | |
| | **FUS+HPI** | 0 | 0 | 3 (100%) | | |
| | **LA+HPI** | 0 | 0 | 6 (100%) | | |
| **3.43** | **Control** | 0 | 0 | 6 (100%) | ns | >0.9999 |
| | **HPI Only** | 0 | 0 | 4 (100%) | | |
| | **Sham FUS** | 0 | 0 | 6 (100%) | | |
| | **FUS Only** | 0 | 0 | 6 (100%) | | |
| | **FUS+HPI** | 0 | 0 | 3 (100%) | | |

version 3

| | | | | | | |
|---|---|---|---|---|---|---|
| | **LA+HPI** | 0 | 0 | 6 (100%) | | |
| **4** | **Control** | 0 | 0 | 5 (100%) | ns | >0.9999 |
| | **HPI Only** | 0 | 0 | 3 (100%) | | |
| | **Sham FUS** | 0 | 0 | 6 (100%) | | |
| | **FUS Only** | 0 | 0 | 6 (100%) | | |
| | **FUS+HPI** | 0 | 0 | 3 (100%) | | |
| | **LA+HPI** | 0 | 0 | 5 (100%) | | |
| **5** | **Control** | 0 | 0 | 5 (100%) | ns | >0.9999 |
| | **HPI Only** | 0 | 0 | 5 (100%) | | |
| | **Sham FUS** | 0 | 0 | 5 (100%) | | |
| | **FUS Only** | 0 | 0 | 4 (100%) | | |
| | **FUS+HPI** | 0 | 0 | 3 (100%) | | |
| | **LA+HPI** | 0 | 0 | 5 (100%) | | |
| **6** | **Control** | 0 | 0 | 5 (100%) | ns | >0.9999 |
| | **HPI Only** | 0 | 0 | 5 (100%) | | |
| | **Sham FUS** | 0 | 0 | 5 (100%) | | |
| | **FUS Only** | 0 | 0 | 4 (100%) | | |

version 3

| | | | | | | |
|---|---|---|---|---|---|---|
| | **FUS+HPI** | 0 | 0 | 3 (100%) | | |
| | **LA+HPI** | 0 | 0 | 5 (100%) | | |
| **7** | **Control** | 0 | 0 | 5 (100%) | ns | >0.9999 |
| | **HPI Only** | 0 | 0 | 5 (100%) | | |
| | **Sham FUS** | 0 | 0 | 5 (100%) | | |
| | **FUS Only** | 0 | 0 | 4 (100%) | | |
| | **FUS+HPI** | 0 | 0 | 3 (100%) | | |
| | **LA+HPI** | 0 | 0 | 5 (100%) | | |
| **8** | **Control** | 0 | 0 | 3 (100%) | ns | >0.9999 |
| | **HPI Only** | 0 | 0 | 5 (100%) | | |
| | **Sham FUS** | 0 | 0 | 5 (100%) | | |
| | **FUS Only** | 0 | 0 | 4 (100%) | | |
| | **FUS+HPI** | 0 | 0 | 3 (100%) | | |
| | **LA+HPI** | 0 | 0 | 5 (100%) | | |
| **9** | **Control** | 0 | 0 | 3 (100%) | ns | >0.9999 |
| | **HPI Only** | 0 | 0 | 5 (100%) | | |
| | **Sham FUS** | 0 | 0 | 3 (100%) | | |

version 3

| | | | | | | |
|---|---|---|---|---|---|---|
| | **FUS Only** | 0 | 0 | 4 (100%) | | |
| | **FUS+HPI** | 0 | 0 | 2 (100%) | | |
| | **LA+HPI** | 0 | 0 | 5 (100%) | | |
| **12** | **Control** | 0 | 0 | 3 (100%) | ns | >0.9999 |
| | **HPI Only** | 0 | 0 | 5 (100%) | | |
| | **Sham FUS** | 0 | 0 | 3 (100%) | | |
| | **FUS Only** | 0 | 0 | 4 (100%) | | |
| | **FUS+HPI** | 0 | 0 | 2 (100%) | | |
| | **LA+HPI** | 0 | 0 | 5 (100%) | | |
| **14** | **Control** | 0 | 0 | 3 (100%) | ns | >0.9999 |
| | **HPI Only** | 0 | 0 | 3 (100%) | | |
| | **Sham FUS** | 0 | 0 | 3 (100%) | | |
| | **FUS Only** | 0 | 0 | 4 (100%) | | |
| | **FUS+HPI** | 0 | 0 | 2 (100%) | | |
| | **LA+HPI** | 0 | 0 | 2 (100%) | | |
| **16** | **Control** | 0 | 0 | 3 (100%) | ns | >0.9999 |
| | **HPI Only** | 0 | 0 | 3 (100%) | | |

version 3

| | | | | |
|---|---|---|---|---|
| | **Sham FUS** | 0 | 0 | 1 (100%) |
| | **FUS Only** | 0 | 0 | 3 (100%) |
| | **FUS+HPI** | 0 | 0 | 2 (100%) |
| | **LA+HPI** | 0 | 0 | 2 (100%) |

**Supplementary Table 14:**Hindpaw (HP) flexion testing (motor and non-pain sensory response) across all experimental groups in male rats, including No Intervention (Control; n = 6), HPI Only (Disease Control; n = 6), Sham FUS (n = 6), FUS Only (n = 6), FUS+HPI (Intervention; n = 5), and LA+HPI (Positive Control; n = 6) at each time point up to 16 weeks. Data were analyzed using contingency table analysis followed by Fisher's exact test, and behavioral responses are presented as fractional response values. FUS = focused ultrasound; HPI = hindpaw incision; LA = local anesthetic.

| Time (Weeks) | Study Arms | No Response N (%) | Partial Flexion N (%) | Full Flexion N (%) | Summary | Fisher's exact p-value |
|---|---|---|---|---|---|---|
| **-0.29** | **Control** | 0 | 0 | 6 (100%) | ns | >0.9999 |
| | **HPI Only** | 0 | 0 | 6 (100%) | | |
| | **Sham FUS** | 0 | 0 | 6 (100%) | | |
| | **FUS Only** | 0 | 0 | 6 (100%) | | |
| | **FUS+HPI** | 0 | 0 | 8 (100%) | | |
| | **LA+HPI** | 0 | 0 | 6 (100%) | | |
| **-0.14** | **Control** | 0 | 0 | 6 (100%) | ns | >0.9999 |
| | **HPI Only** | 0 | 0 | 6 (100%) | | |

version 3

| | | | | | | |
|---|---|---|---|---|---|---|
| | **Sham FUS** | 0 | 0 | 6 (100%) | | |
| | **FUS Only** | 0 | 0 | 6 (100%) | | |
| | **FUS+HPI** | 0 | 0 | 8 (100%) | | |
| | **LA+HPI** | 0 | 0 | 6 (100%) | | |
| **0** | **Control** | 0 | 0 | 6 (100%) | **** | <0.0001 |
| | **HPI Only** | 0 | 0 | 6 (100%) | | |
| | **Sham FUS** | 0 | 0 | 6 (100%) | | |
| | **FUS Only** | 0 | 0 | 6 (100%) | | |
| | **FUS+HPI** | 0 | 0 | 8 (100%) | | |
| | **LA+HPI** | 5 (83.33%) | 1 (16.67%) | 0 | | |
| **0.14** | **Control** | 0 | 0 | 6 (100%) | ns | >0.9999 |
| | **HPI Only** | 0 | 0 | 6 (100%) | | |
| | **Sham FUS** | 0 | 0 | 6 (100%) | | |
| | **FUS Only** | 0 | 0 | 6 (100%) | | |
| | **FUS+HPI** | 0 | 0 | 8 (100%) | | |
| | **LA+HPI** | 0 | 0 | 6 (100%) | | |
| **0.43** | **Control** | 0 | 0 | 6 (100%) | ns | >0.3583 |

version 3

| | | | | | | |
|---|---|---|---|---|---|---|
| | **HPI Only** | 0 | 0 | 6 (100%) | | |
| | **Sham FUS** | 0 | 0 | 6 (100%) | | |
| | **FUS Only** | 1 (16.67%) | 1 (16.67%) | 4 (66.67%) | | |
| | **FUS+HPI** | 0 | 0 | 4 (100%) | | |
| | **LA+HPI** | 0 | 0 | 6 (100%) | | |
| **1** | **Control** | 0 | 0 | 6 (100%) | ns | >0.9999 |
| | **HPI Only** | 0 | 0 | 6 (100%) | | |
| | **Sham FUS** | 0 | 0 | 6 (100%) | | |
| | **FUS Only** | 0 | 1 (8.33%) | 6 (91.67%) | | |
| | **FUS+HPI** | 0 | 0 | 8 (100%) | | |
| | **LA+HPI** | 0 | 0 | 6 (100%) | | |
| **1.43** | **Control** | 0 | 0 | 6 (100%) | ns | >0.9999 |
| | **HPI Only** | 0 | 0 | 6 (100%) | | |
| | **Sham FUS** | 0 | 0 | 6 (100%) | | |
| | **FUS Only** | 0 | 0 | 6 (100%) | | |
| | **FUS+HPI** | 0 | 0 | 8 (100%) | | |
| | **LA+HPI** | 0 | 0 | 6 (100%) | | |

version 3

| | | | | | | |
|---|---|---|---|---|---|---|
| **2** | **Control** | 0 | 0 | 4 (100%) | ns | >0.9999 |
| | **HPI Only** | 0 | 0 | 6 (100%) | | |
| | **Sham FUS** | 0 | 0 | 6 (100%) | | |
| | **FUS Only** | 0 | 0 | 6 (100%) | | |
| | **FUS+HPI** | 0 | 0 | 8 (100%) | | |
| | **LA+HPI** | 0 | 0 | 4 (100%) | | |
| **2.43** | **Control** | 0 | 0 | 4 (100%) | ns | >0.9999 |
| | **HPI Only** | 0 | 0 | 6 (100%) | | |
| | **Sham FUS** | 0 | 0 | 4 (100%) | | |
| | **FUS Only** | 0 | 0 | 4 (100%) | | |
| | **FUS+HPI** | 0 | 0 | 7 (100%) | | |
| | **LA+HPI** | 0 | 0 | 4 (100%) | | |
| **3** | **Control** | 0 | 0 | 4 (100%) | ns | >0.9999 |
| | **HPI Only** | 0 | 0 | 6 (100%) | | |
| | **Sham FUS** | 0 | 0 | 4 (100%) | | |
| | **FUS Only** | 0 | 0 | 4 (100%) | | |
| | **FUS+HPI** | 0 | 0 | 7 (100%) | | |

version 3

| | | | | | | |
|---|---|---|---|---|---|---|
| | **LA+HPI** | 0 | 0 | 4 (100%) | | |
| **3.43** | **Control** | 0 | 0 | 4 (100%) | ns | >0.9999 |
| | **HPI Only** | 0 | 0 | 6 (100%) | | |
| | **Sham FUS** | 0 | 0 | 4 (100%) | | |
| | **FUS Only** | 0 | 0 | 4 (100%) | | |
| | **FUS+HPI** | 0 | 0 | 7 (100%) | | |
| | **LA+HPI** | 0 | 0 | 4 (100%) | | |
| **4** | **Control** | 0 | 0 | 3 (100%) | ns | >0.9999 |
| | **HPI Only** | 0 | 0 | 5 (100%) | | |
| | **Sham FUS** | 0 | 0 | 4 (100%) | | |
| | **FUS Only** | 0 | 0 | 4 (100%) | | |
| | **FUS+HPI** | 0 | 0 | 7 (100%) | | |
| | **LA+HPI** | 0 | 0 | 3 (100%) | | |
| **5** | **Control** | 0 | 0 | 3 (100%) | ns | >0.9999 |
| | **HPI Only** | 0 | 0 | 5 (100%) | | |
| | **Sham FUS** | 0 | 0 | 3 (100%) | | |
| | **FUS Only** | 0 | 0 | 4 (100%) | | |

version 3

| | | | | | | |
|---|---|---|---|---|---|---|
| | **FUS+HPI** | 0 | 0 | 5 (100%) | | |
| | **LA+HPI** | 0 | 0 | 3 (100%) | | |
| **6** | **Control** | 0 | 0 | 3 (100%) | ns | >0.9999 |
| | **HPI Only** | 0 | 0 | 5 (100%) | | |
| | **Sham FUS** | 0 | 0 | 3 (100%) | | |
| | **FUS Only** | 0 | 0 | 4 (100%) | | |
| | **FUS+HPI** | 0 | 0 | 5 (100%) | | |
| | **LA+HPI** | 0 | 0 | 3 (100%) | | |
| **7** | **Control** | 0 | 0 | 3 (100%) | ns | >0.9999 |
| | **HPI Only** | 0 | 0 | 5 (100%) | | |
| | **Sham FUS** | 0 | 0 | 3 (100%) | | |
| | **FUS Only** | 0 | 0 | 4 (100%) | | |
| | **FUS+HPI** | 0 | 0 | 5 (100%) | | |
| | **LA+HPI** | 0 | 0 | 3 (100%) | | |
| **8** | **Control** | 0 | 0 | 3 (100%) | ns | >0.9999 |
| | **HPI Only** | 0 | 0 | 3 (100%) | | |
| | **Sham FUS** | 0 | 0 | 3 (100%) | | |

version 3

| | | | | | | |
|---|---|---|---|---|---|---|
| | **FUS Only** | 0 | 0 | 4 (100%) | | |
| | **FUS+HPI** | 0 | 0 | 5 (100%) | | |
| | **LA+HPI** | 0 | 0 | 3 (100%) | | |
| **9** | **Control** | 0 | 0 | 3 (100%) | ns | >0.9999 |
| | **HPI Only** | 0 | 0 | 3 (100%) | | |
| | **Sham FUS** | 0 | 0 | 3 (100%) | | |
| | **FUS Only** | 0 | 0 | 2 (100%) | | |
| | **FUS+HPI** | 0 | 0 | 4 (100%) | | |
| | **LA+HPI** | 0 | 0 | 3 (100%) | | |
| **12** | **Control** | 0 | 0 | 3 (100%) | ns | >0.9999 |
| | **HPI Only** | 0 | 0 | 3 (100%) | | |
| | **Sham FUS** | 0 | 0 | 3 (100%) | | |
| | **FUS Only** | 0 | 0 | 2 (100%) | | |
| | **FUS+HPI** | 0 | 0 | 4 (100%) | | |
| | **LA+HPI** | 0 | 0 | 3 (100%) | | |
| **14** | **Control** | 0 | 0 | 3 (100%) | ns | >0.9999 |
| | **HPI Only** | 0 | 0 | 1 (100%) | | |

version 3

| | | | | | | |
|---|---|---|---|---|---|---|
| | **Sham FUS** | 0 | 0 | 3 (100%) | | |
| | **FUS Only** | 0 | 0 | 2 (100%) | | |
| | **FUS+HPI** | 0 | 0 | 3 (100%) | | |
| | **LA+HPI** | 0 | 0 | 3 (100%) | | |
| **16** | **Control** | 0 | 0 | 3 (100%) | ns | >0.9999 |
| | **HPI Only** | 0 | 0 | 1 (100%) | | |
| | **Sham FUS** | 0 | 0 | 3 (100%) | | |
| | **FUS Only** | 0 | 0 | 1 (100%) | | |
| | **FUS+HPI** | 0 | 0 | 2 (100%) | | |
| | **LA+HPI** | 0 | 0 | 3 (100%) | | |

Version 3

**Supplementary Table 15:** Summary of group-wise comparisons for Randall–Selitto testing (hindpaw mechanical withdrawal thresholds) across all experimental groups, including No Intervention (Control; n = 12), HPI Only (Disease Control; n = 12), Sham FUS (n = 12), FUS Only (n = 12), FUS+HPI (Intervention; n = 12), and LA+HPI (Positive Control; n = 12) at each time point up to 16 weeks. Measurements were analyzed using a two-way mixed-effects ANOVA with fixed effects for treatment group and time, followed by Bonferroni-corrected post hoc multiple-comparison testing. Statistical significance is denoted as follows: ns ($p \geq 0.05$), * ($p < 0.05$), ** ($p < 0.01$), *** ($p < 0.001$).

| Time (Weeks) | Study Arms | No Response N (%) | Partial Extension N (%) | Full Extension N (%) | Summary | Fisher's exact p-value |
|---|---|---|---|---|---|---|
| **-0.29** | **Control** | 0 | 0 | 12 (100%) | ns | >0.9999 |
| | **HPI Only** | 0 | 0 | 12 (100%) | | |

| | | | | | | |
|---|---|---|---|---|---|---|
| | **Sham FUS** | 0 | 0 | 12 (100%) | | |
| | **FUS Only** | 0 | 0 | 12 (100%) | | |
| | **FUS+HPI** | 0 | 0 | 12 (100%) | | |
| | **LA+HPI** | 0 | 0 | 12 (100%) | | |
| **-0.14** | **Control** | 0 | 0 | 12 (100%) | ns | >0.9999 |
| | **HPI Only** | 0 | 0 | 12 (100%) | | |
| | **Sham FUS** | 0 | 0 | 12 (100%) | | |
| | **FUS Only** | 0 | 0 | 12 (100%) | | |
| | **FUS+HPI** | 0 | 0 | 12 (100%) | | |
| | **LA+HPI** | 0 | 0 | 12 (100%) | | |
| **0** | **Control** | 0 | 0 | 12 (100%) | **** | <0.0001 |
| | **HPI Only** | 0 | 0 | 12 (100%) | | |
| | **Sham FUS** | 0 | 0 | 12 (100%) | | |
| | **FUS Only** | 1 (8.33%) | 1 (8.33%) | 10 (83.33%) | | |
| | **FUS+HPI** | 1 (8.33%) | 0 | 11 (91.67%) | | |
| | **LA+HPI** | 11 (91.67%) | 1 (8.33%) | 0 | | |
| **0.14** | **Control** | 0 | 0 | 12 (100%) | ns | 0.4205 |

version 3

| | | | | | | |
|---|---|---|---|---|---|---|
| | **HPI Only** | 0 | 0 | 12 (100%) | | |
| | **Sham FUS** | 0 | 0 | 12 (100%) | | |
| | **FUS Only** | 1 (8.33%) | 1 (8.33%) | 10 (83.33%) | | |
| | **FUS+HPI** | 1 (8.33%) | 0 | 11 (91.67%) | | |
| | **LA+HPI** | 0 | 0 | 12 (100%) | | |
| **0.43** | **Control** | 0 | 0 | 12 (100%) | ns | 0.1549 |
| | **HPI Only** | 0 | 0 | 12 (100%) | | |
| | **Sham FUS** | 0 | 0 | 12 (100%) | | |
| | **FUS Only** | 1 (8.33%) | 1 (8.33%) | 10 (83.33%) | | |
| | **FUS+HPI** | 0 | 0 | 12 (100%) | | |
| | **LA+HPI** | 0 | 0 | 12 (100%) | | |
| **1** | **Control** | 0 | 0 | 12 (100%) | ns | >0.9999 |
| | **HPI Only** | 0 | 0 | 12 (100%) | | |
| | **Sham FUS** | 0 | 0 | 12 (100%) | | |
| | **FUS Only** | 1 (8.33%) | 0 | 11 (91.67%) | | |
| | **FUS+HPI** | 0 | 0 | 12 (100%) | | |
| | **LA+HPI** | 0 | 0 | 12 (100%) | | |

version 3

| | | | | | | |
|---|---|---|---|---|---|---|
| **1.43** | **Control** | 0 | 0 | 12 (100%) | ns | >0.9999 |
| | **HPI Only** | 0 | 0 | 12 (100%) | | |
| | **Sham FUS** | 0 | 0 | 12 (100%) | | |
| | **FUS Only** | 1 (8.33%) | 0 | 11 (91.67%) | | |
| | **FUS+HPI** | 0 | 0 | 12 (100%) | | |
| | **LA+HPI** | 0 | 0 | 12 (100%) | | |
| **2** | **Control** | 0 | 0 | 10 (100%) | ns | >0.9999 |
| | **HPI Only** | 0 | 0 | 10 (100%) | | |
| | **Sham FUS** | 0 | 0 | 12 (100%) | | |
| | **FUS Only** | 0 | 0 | 12 (100%) | | |
| | **FUS+HPI** | 0 | 1 (8.33%) | 11 (91.67%) | | |
| | **LA+HPI** | 0 | 0 | 10 (100%) | | |
| **2.43** | **Control** | 0 | 0 | 10 (100%) | ns | >0.9999 |
| | **HPI Only** | 0 | 0 | 10 (100%) | | |
| | **Sham FUS** | 0 | 0 | 10 (100%) | | |
| | **FUS Only** | 0 | 0 | 10 (100%) | | |
| | **FUS+HPI** | 0 | 0 | 10 (100%) | | |

version 3

| | | | | | | |
|---|---|---|---|---|---|---|
| | **LA+HPI** | 0 | 0 | 10 (100%) | | |
| **3** | **Control** | 0 | 0 | 10 (100%) | ns | >0.9999 |
| | **HPI Only** | 0 | 0 | 10 (100%) | | |
| | **Sham FUS** | 0 | 0 | 10 (100%) | | |
| | **FUS Only** | 0 | 0 | 10 (100%) | | |
| | **FUS+HPI** | 0 | 0 | 10 (100%) | | |
| | **LA+HPI** | 0 | 0 | 10 (100%) | | |
| **3.43** | **Control** | 0 | 0 | 10 (100%) | ns | >0.9999 |
| | **HPI Only** | 0 | 0 | 10 (100%) | | |
| | **Sham FUS** | 0 | 0 | 10 (100%) | | |
| | **FUS Only** | 0 | 0 | 10 (100%) | | |
| | **FUS+HPI** | 0 | 0 | 10 (100%) | | |
| | **LA+HPI** | 0 | 0 | 10 (100%) | | |
| **4** | **Control** | 0 | 0 | 8 (100%) | ns | >0.9999 |
| | **HPI Only** | 0 | 0 | 8 (100%) | | |
| | **Sham FUS** | 0 | 0 | 10 (100%) | | |
| | **FUS Only** | 0 | 0 | 10 (100%) | | |

version3

| | | | | | | |
|---|---|---|---|---|---|---|
| | **FUS+HPI** | 0 | 0 | 10 (100%) | | |
| | **LA+HPI** | 0 | 0 | 8 (100%) | | |
| **5** | **Control** | 0 | 0 | 8 (100%) | ns | >0.9999 |
| | **HPI Only** | 0 | 0 | 8 (100%) | | |
| | **Sham FUS** | 0 | 0 | 8 (100%) | | |
| | **FUS Only** | 0 | 0 | 8 (100%) | | |
| | **FUS+HPI** | 0 | 0 | 8 (100%) | | |
| | **LA+HPI** | 0 | 0 | 8 (100%) | | |
| **6** | **Control** | 0 | 0 | 8 (100%) | ns | >0.9999 |
| | **HPI Only** | 0 | 0 | 8 (100%) | | |
| | **Sham FUS** | 0 | 0 | 8 (100%) | | |
| | **FUS Only** | 0 | 0 | 8 (100%) | | |
| | **FUS+HPI** | 0 | 0 | 8 (100%) | | |
| | **LA+HPI** | 0 | 0 | 8 (100%) | | |
| **7** | **Control** | 0 | 0 | 8 (100%) | ns | >0.9999 |
| | **HPI Only** | 0 | 0 | 8 (100%) | | |
| | **Sham FUS** | 0 | 0 | 8 (100%) | | |

version3

| | | | | | | |
|---|---|---|---|---|---|---|
| | **FUS Only** | 0 | 0 | 8 (100%) | | |
| | **FUS+HPI** | 0 | 0 | 8 (100%) | | |
| | **LA+HPI** | 0 | 0 | 8 (100%) | | |
| **8** | **Control** | 0 | 0 | 6 (100%) | ns | >0.9999 |
| | **HPI Only** | 0 | 0 | 6 (100%) | | |
| | **Sham FUS** | 0 | 0 | 8 (100%) | | |
| | **FUS Only** | 0 | 0 | 8 (100%) | | |
| | **FUS+HPI** | 0 | 0 | 8 (100%) | | |
| | **LA+HPI** | 0 | 0 | 8 (100%) | | |
| **9** | **Control** | 0 | 0 | 6 (100%) | ns | >0.9999 |
| | **HPI Only** | 0 | 0 | 6 (100%) | | |
| | **Sham FUS** | 0 | 0 | 6 (100%) | | |
| | **FUS Only** | 0 | 0 | 6 (100%) | | |
| | **FUS+HPI** | 0 | 0 | 6 (100%) | | |
| | **LA+HPI** | 0 | 0 | 8 (100%) | | |
| **12** | **Control** | 0 | 0 | 6 (100%) | ns | >0.9999 |
| | **HPI Only** | 0 | 0 | 6 (100%) | | |

version3

| | | | | | | |
|---|---|---|---|---|---|---|
| | **Sham FUS** | 0 | 0 | 6 (100%) | | |
| | **FUS Only** | 0 | 0 | 6 (100%) | | |
| | **FUS+HPI** | 0 | 0 | 6 (100%) | | |
| | **LA+HPI** | 0 | 0 | 8 (100%) | | |
| **14** | **Control** | 0 | 0 | 6 (100%) | ns | >0.9999 |
| | **HPI Only** | 0 | 0 | 4 (100%) | | |
| | **Sham FUS** | 0 | 0 | 6 (100%) | | |
| | **FUS Only** | 0 | 0 | 6 (100%) | | |
| | **FUS+HPI** | 0 | 0 | 5 (100%) | | |
| | **LA+HPI** | 0 | 0 | 5 (100%) | | |
| **16** | **Control** | 0 | 0 | 6 (100%) | ns | >0.9999 |
| | **HPI Only** | 0 | 0 | 4 (100%) | | |
| | **Sham FUS** | 0 | 0 | 4 (100%) | | |
| | **FUS Only** | 0 | 0 | 4 (100%) | | |
| | **FUS+HPI** | 0 | 0 | 4 (100%) | | |
| | **LA+HPI** | 0 | 0 | 5 (100%) | | |

version 3

**Supplementary Table 16:** Hindpaw (HP) extension testing (motor response/startle reflex assessment) across all experimental groups, including No Intervention (Control; n = 12), HPI

Only (Disease Control; n = 12), Sham FUS (n = 12), FUS Only (n = 12), FUS+HPI (Intervention; n = 12), and LA+HPI (Positive Control; n = 12) at each time point up to 16 weeks. Data were analyzed using contingency table analysis followed by Fisher's exact test, and behavioral responses are presented as fractional response values. FUS = focused ultrasound; HPI = hindpaw incision; LA = local anesthetic.

| Time (Weeks) | Study Arms | No Response N (%) | Partial Extension N (%) | Full Extension N (%) | Summary | Fisher's exact p-value |
|---|---|---|---|---|---|---|
| -0.29 | Control | 0 | 0 | 6 (100%) | ns | >0.9999 |
| | HPI Only | 0 | 0 | 6 (100%) | | |
| | Sham FUS | 0 | 0 | 6 (100%) | | |
| | FUS Only | 0 | 0 | 6 (100%) | | |
| | FUS+HPI | 0 | 0 | 4 (100%) | | |
| | LA+HPI | 0 | 0 | 6 (100%) | | |
| -0.14 | Control | 0 | 0 | 6 (100%) | ns | >0.9999 |
| | HPI Only | 0 | 0 | 6 (100%) | | |
| | Sham FUS | 0 | 0 | 6 (100%) | | |
| | FUS Only | 0 | 0 | 6 (100%) | | |
| | FUS+HPI | 0 | 0 | 4 (100%) | | |
| | LA+HPI | 0 | 0 | 6 (100%) | | |
| 0 | Control | 0 | 0 | 6 (100%) | **** | <0.0001 |
| | HPI Only | 0 | 0 | 6 (100%) | | |

version 3

| | | | | | | |
|---|---|---|---|---|---|---|
| | **Sham FUS** | 0 | 0 | 6 (100%) | | |
| | **FUS Only** | 0 | 1 (16.67%) | 5 (83.33%) | | |
| | **FUS+HPI** | 0 | 0 | 4 (100%) | | |
| | **LA+HPI** | 2 (33.33%) | 4 (66.67%) | 0 | | |
| **0.14** | **Control** | 0 | 0 | 6 (100%) | ns | >0.9999 |
| | **HPI Only** | 0 | 0 | 6 (100%) | | |
| | **Sham FUS** | 0 | 0 | 6 (100%) | | |
| | **FUS Only** | 0 | 1 (16.67%) | 5 (83.33%) | | |
| | **FUS+HPI** | 0 | 0 | 4 (100%) | | |
| | **LA+HPI** | 0 | 0 | 6 (100%) | | |
| **0.43** | **Control** | 0 | 0 | 6 (100%) | ns | >0.9999 |
| | **HPI Only** | 0 | 0 | 6 (100%) | | |
| | **Sham FUS** | 0 | 0 | 6 (100%) | | |
| | **FUS Only** | 0 | 1 (16.67%) | 5 (83.33%) | | |
| | **FUS+HPI** | 0 | 0 | 4 (100%) | | |
| | **LA+HPI** | 0 | 0 | 6 (100%) | | |
| **1** | **Control** | 0 | 0 | 6 (100%) | ns | >0.9999 |

version 3

| | | | | | | |
|---|---|---|---|---|---|---|
| | **HPI Only** | 0 | 0 | 6 (100%) | | |
| | **Sham FUS** | 0 | 0 | 6 (100%) | | |
| | **FUS Only** | 0 | 1 (16.67%) | 5 (83.33%) | | |
| | **FUS+HPI** | 0 | 0 | 4 (100%) | | |
| | **LA+HPI** | 0 | 0 | 6 (100%) | | |
| **1.43** | **Control** | 0 | 0 | 6 (100%) | ns | >0.9999 |
| | **HPI Only** | 0 | 0 | 6 (100%) | | |
| | **Sham FUS** | 0 | 0 | 6 (100%) | | |
| | **FUS Only** | 0 | 0 | 6 (100%) | | |
| | **FUS+HPI** | 0 | 0 | 4 (100%) | | |
| | **LA+HPI** | 0 | 0 | 6 (100%) | | |
| **2** | **Control** | 0 | 0 | 6 (100%) | ns | >0.9999 |
| | **HPI Only** | 0 | 0 | 4 (100%) | | |
| | **Sham FUS** | 0 | 0 | 6 (100%) | | |
| | **FUS Only** | 0 | 0 | 6 (100%) | | |
| | **FUS+HPI** | 0 | 0 | 4 (100%) | | |
| | **LA+HPI** | 0 | 0 | 6 (100%) | | |

version 3

| | | | | | | |
|---|---|---|---|---|---|---|
| **2.43** | **Control** | 0 | 0 | 6 (100%) | ns | >0.9999 |
| | **HPI Only** | 0 | 0 | 4 (100%) | | |
| | **Sham FUS** | 0 | 0 | 6 (100%) | | |
| | **FUS Only** | 0 | 0 | 6 (100%) | | |
| | **FUS+HPI** | 0 | 0 | 3 (100%) | | |
| | **LA+HPI** | 0 | 0 | 6 (100%) | | |
| **3** | **Control** | 0 | 0 | 6 (100%) | ns | >0.9999 |
| | **HPI Only** | 0 | 0 | 4 (100%) | | |
| | **Sham FUS** | 0 | 0 | 6 (100%) | | |
| | **FUS Only** | 0 | 0 | 6 (100%) | | |
| | **FUS+HPI** | 0 | 0 | 3 (100%) | | |
| | **LA+HPI** | 0 | 0 | 6 (100%) | | |
| **3.43** | **Control** | 0 | 0 | 6 (100%) | ns | >0.9999 |
| | **HPI Only** | 0 | 0 | 4 (100%) | | |
| | **Sham FUS** | 0 | 0 | 6 (100%) | | |
| | **FUS Only** | 0 | 0 | 6 (100%) | | |
| | **FUS+HPI** | 0 | 0 | 3 (100%) | | |

version 3

| | | | | | | |
|---|---|---|---|---|---|---|
| | **LA+HPI** | 0 | 0 | 6 (100%) | | |
| **4** | **Control** | 0 | 0 | 5 (100%) | ns | >0.9999 |
| | **HPI Only** | 0 | 0 | 3 (100%) | | |
| | **Sham FUS** | 0 | 0 | 6 (100%) | | |
| | **FUS Only** | 0 | 0 | 6 (100%) | | |
| | **FUS+HPI** | 0 | 0 | 3 (100%) | | |
| | **LA+HPI** | 0 | 0 | 5 (100%) | | |
| **5** | **Control** | 0 | 0 | 5 (100%) | ns | >0.9999 |
| | **HPI Only** | 0 | 0 | 5 (100%) | | |
| | **Sham FUS** | 0 | 0 | 5 (100%) | | |
| | **FUS Only** | 0 | 0 | 4 (100%) | | |
| | **FUS+HPI** | 0 | 0 | 3 (100%) | | |
| | **LA+HPI** | 0 | 0 | 5 (100%) | | |
| **6** | **Control** | 0 | 0 | 5 (100%) | ns | >0.9999 |
| | **HPI Only** | 0 | 0 | 5 (100%) | | |
| | **Sham FUS** | 0 | 0 | 5 (100%) | | |
| | **FUS Only** | 0 | 0 | 4 (100%) | | |

version3

| | | | | | | |
|---|---|---|---|---|---|---|
| | **FUS+HPI** | 0 | 0 | 3 (100%) | | |
| | **LA+HPI** | 0 | 0 | 5 (100%) | | |
| **7** | **Control** | 0 | 0 | 5 (100%) | ns | >0.9999 |
| | **HPI Only** | 0 | 0 | 5 (100%) | | |
| | **Sham FUS** | 0 | 0 | 5 (100%) | | |
| | **FUS Only** | 0 | 0 | 4 (100%) | | |
| | **FUS+HPI** | 0 | 0 | 3 (100%) | | |
| | **LA+HPI** | 0 | 0 | 5 (100%) | | |
| **8** | **Control** | 0 | 0 | 3 (100%) | ns | >0.9999 |
| | **HPI Only** | 0 | 0 | 5 (100%) | | |
| | **Sham FUS** | 0 | 0 | 5 (100%) | | |
| | **FUS Only** | 0 | 0 | 4 (100%) | | |
| | **FUS+HPI** | 0 | 0 | 3 (100%) | | |
| | **LA+HPI** | 0 | 0 | 5 (100%) | | |
| **9** | **Control** | 0 | 0 | 3 (100%) | ns | >0.9999 |
| | **HPI Only** | 0 | 0 | 5 (100%) | | |
| | **Sham FUS** | 0 | 0 | 3 (100%) | | |

version3

| | | | | | | |
|---|---|---|---|---|---|---|
| | **FUS Only** | 0 | 0 | 4 (100%) | | |
| | **FUS+HPI** | 0 | 0 | 2 (100%) | | |
| | **LA+HPI** | 0 | 0 | 5 (100%) | | |
| **12** | **Control** | 0 | 0 | 3 (100%) | ns | >0.9999 |
| | **HPI Only** | 0 | 0 | 5 (100%) | | |
| | **Sham FUS** | 0 | 0 | 3 (100%) | | |
| | **FUS Only** | 0 | 0 | 4 (100%) | | |
| | **FUS+HPI** | 0 | 0 | 2 (100%) | | |
| | **LA+HPI** | 0 | 0 | 5 (100%) | | |
| **14** | **Control** | 0 | 0 | 3 (100%) | ns | >0.9999 |
| | **HPI Only** | 0 | 0 | 3 (100%) | | |
| | **Sham FUS** | 0 | 0 | 3 (100%) | | |
| | **FUS Only** | 0 | 0 | 4 (100%) | | |
| | **FUS+HPI** | 0 | 0 | 2 (100%) | | |
| | **LA+HPI** | 0 | 0 | 2 (100%) | | |
| **16** | **Control** | 0 | 0 | 3 (100%) | ns | >0.9999 |

version 3

**Supplementary Table 17:** Hindpaw (HP) extension testing (motor response/startle reflex assessment) across all experimental groups in male rats, including No Intervention (Control; n = 6), HPI Only (Disease Control; n = 6), Sham FUS (n = 6), FUS Only (n = 6), FUS+HPI

(Intervention; n = 5), and LA+HPI (Positive Control; n = 6) at each time point up to 16 weeks. Data were analyzed using contingency table analysis followed by Fisher's exact test, and behavioral responses are presented as fractional response values. FUS = focused ultrasound; HPI = hindpaw incision; LA = local anesthetic.

| Time (Weeks) | Study Arms | No Response N (%) | Partial Extension N (%) | Full Extension N (%) | Summary | Fisher's exact p-value |
|---|---|---|---|---|---|---|
| **-0.29** | **Control** | 0 | 0 | 6 (100%) | ns | >0.9999 |
| | **HPI Only** | 0 | 0 | 6 (100%) | | |
| | **Sham FUS** | 0 | 0 | 6 (100%) | | |
| | **FUS Only** | 0 | 0 | 6 (100%) | | |
| | **FUS+HPI** | 0 | 0 | 8 (100%) | | |
| | **LA+HPI** | 0 | 0 | 6 (100%) | | |
| **-0.14** | **Control** | 0 | 0 | 6 (100%) | ns | >0.9999 |
| | **HPI Only** | 0 | 0 | 6 (100%) | | |
| | **Sham FUS** | 0 | 0 | 6 (100%) | | |
| | **FUS Only** | 0 | 0 | 6 (100%) | | |
| | **FUS+HPI** | 0 | 0 | 8 (100%) | | |
| | **LA+HPI** | 0 | 0 | 6 (100%) | | |
| **0** | **Control** | 0 | 0 | 6 (100%) | **** | <0.0001 |
| | **HPI Only** | 0 | 0 | 6 (100%) | | |

version 3

| | | | | | | |
|---|---|---|---|---|---|---|
| | **Sham FUS** | 0 | 0 | 6 (100%) | | |
| | **FUS Only** | 0 | 0 | 6 (100%) | | |
| | **FUS+HPI** | 0 | 1 (12.5%) | 7 (87.5%) | | |
| | **LA+HPI** | 6 (100%) | 0 | 0 | | |
| **0.14** | **Control** | 0 | 0 | 6 (100%) | ns | >0.9999 |
| | **HPI Only** | 0 | 0 | 6 (100%) | | |
| | **Sham FUS** | 0 | 0 | 6 (100%) | | |
| | **FUS Only** | 0 | 0 | 6 (100%) | | |
| | **FUS+HPI** | 1 (12.5%) | 0 | 7 (87.5%) | | |
| | **LA+HPI** | 0 | 0 | 6 (100%) | | |
| **0.43** | **Control** | 0 | 0 | 6 (100%) | ns | >0.9999 |
| | **HPI Only** | 0 | 0 | 6 (100%) | | |
| | **Sham FUS** | 0 | 0 | 6 (100%) | | |
| | **FUS Only** | 0 | 0 | 8 (100%) | | |
| | **FUS+HPI** | 0 | 0 | 8 (100%) | | |
| | **LA+HPI** | 0 | 0 | 6 (100%) | | |
| **1** | **Control** | 0 | 0 | 6 (100%) | ns | >0.9999 |

version 3

| | | | | | | |
|---|---|---|---|---|---|---|
| | **HPI Only** | 0 | 0 | 6 (100%) | | |
| | **Sham FUS** | 0 | 0 | 6 (100%) | | |
| | **FUS Only** | 0 | 0 | 6 (100%) | | |
| | **FUS+HPI** | 1 (12.5%) | 0 | 7 (87.5%) | | |
| | **LA+HPI** | 0 | 0 | 6 (100%) | | |
| **1.43** | **Control** | 0 | 0 | 6 (100%) | ns | >0.9999 |
| | **HPI Only** | 0 | 0 | 6 (100%) | | |
| | **Sham FUS** | 0 | 0 | 6 (100%) | | |
| | **FUS Only** | 0 | 0 | 6 (100%) | | |
| | **FUS+HPI** | 1 (12.5%) | 0 | 7 (87.5%) | | |
| | **LA+HPI** | 0 | 0 | 6 (100%) | | |
| **2** | **Control** | 0 | 0 | 4 (100%) | ns | >0.9999 |
| | **HPI Only** | 0 | 0 | 6 (100%) | | |
| | **Sham FUS** | 0 | 0 | 6 (100%) | | |
| | **FUS Only** | 0 | 0 | 6 (100%) | | |
| | **FUS+HPI** | 0 | 1 (12.5%) | 7 (87.5%) | | |
| | **LA+HPI** | 0 | 0 | 4 (100%) | | |

version 3

| | | | | | | |
|---|---|---|---|---|---|---|
| **2.43** | **Control** | 0 | 0 | 4 (100%) | ns | >0.9999 |
| | **HPI Only** | 0 | 0 | 6 (100%) | | |
| | **Sham FUS** | 0 | 0 | 4 (100%) | | |
| | **FUS Only** | 0 | 0 | 4 (100%) | | |
| | **FUS+HPI** | 0 | 0 | 7 (100%) | | |
| | **LA+HPI** | 0 | 0 | 4 (100%) | | |
| **3** | **Control** | 0 | 0 | 4 (100%) | ns | >0.9999 |
| | **HPI Only** | 0 | 0 | 6 (100%) | | |
| | **Sham FUS** | 0 | 0 | 4 (100%) | | |
| | **FUS Only** | 0 | 0 | 4 (100%) | | |
| | **FUS+HPI** | 0 | 0 | 7 (100%) | | |
| | **LA+HPI** | 0 | 0 | 4 (100%) | | |
| **3.43** | **Control** | 0 | 0 | 4 (100%) | ns | >0.9999 |
| | **HPI Only** | 0 | 0 | 6 (100%) | | |
| | **Sham FUS** | 0 | 0 | 4 (100%) | | |
| | **FUS Only** | 0 | 0 | 4 (100%) | | |
| | **FUS+HPI** | 0 | 0 | 7 (100%) | | |

version 3

| | | | | | | |
|---|---|---|---|---|---|---|
| | **LA+HPI** | 0 | 0 | 4 (100%) | | |
| **4** | **Control** | 0 | 0 | 3 (100%) | ns | >0.9999 |
| | **HPI Only** | 0 | 0 | 5 (100%) | | |
| | **Sham FUS** | 0 | 0 | 4 (100%) | | |
| | **FUS Only** | 0 | 0 | 4 (100%) | | |
| | **FUS+HPI** | 0 | 0 | 7 (100%) | | |
| | **LA+HPI** | 0 | 0 | 3 (100%) | | |
| **5** | **Control** | 0 | 0 | 3 (100%) | ns | >0.9999 |
| | **HPI Only** | 0 | 0 | 5 (100%) | | |
| | **Sham FUS** | 0 | 0 | 3 (100%) | | |
| | **FUS Only** | 0 | 0 | 4 (100%) | | |
| | **FUS+HPI** | 0 | 0 | 5 (100%) | | |
| | **LA+HPI** | 0 | 0 | 3 (100%) | | |
| **6** | **Control** | 0 | 0 | 3 (100%) | ns | >0.9999 |
| | **HPI Only** | 0 | 0 | 5 (100%) | | |
| | **Sham FUS** | 0 | 0 | 3 (100%) | | |
| | **FUS Only** | 0 | 0 | 4 (100%) | | |

version3

| | | | | | | |
|---|---|---|---|---|---|---|
| | **FUS+HPI** | 0 | 0 | 5 (100%) | | |
| | **LA+HPI** | 0 | 0 | 3 (100%) | | |
| **7** | **Control** | 0 | 0 | 3 (100%) | ns | >0.9999 |
| | **HPI Only** | 0 | 0 | 5 (100%) | | |
| | **Sham FUS** | 0 | 0 | 3 (100%) | | |
| | **FUS Only** | 0 | 0 | 4 (100%) | | |
| | **FUS+HPI** | 0 | 0 | 5 (100%) | | |
| | **LA+HPI** | 0 | 0 | 3 (100%) | | |
| **8** | **Control** | 0 | 0 | 3 (100%) | ns | >0.9999 |
| | **HPI Only** | 0 | 0 | 3 (100%) | | |
| | **Sham FUS** | 0 | 0 | 3 (100%) | | |
| | **FUS Only** | 0 | 0 | 4 (100%) | | |
| | **FUS+HPI** | 0 | 0 | 5 (100%) | | |
| | **LA+HPI** | 0 | 0 | 3 (100%) | | |
| **9** | **Control** | 0 | 0 | 3 (100%) | ns | >0.9999 |
| | **HPI Only** | 0 | 0 | 3 (100%) | | |
| | **Sham FUS** | 0 | 0 | 3 (100%) | | |

version3

| | | | | | | |
|---|---|---|---|---|---|---|
| | **FUS Only** | 0 | 0 | 2 (100%) | | |
| | **FUS+HPI** | 0 | 0 | 4 (100%) | | |
| | **LA+HPI** | 0 | 0 | 3 (100%) | | |
| **12** | **Control** | 0 | 0 | 3 (100%) | ns | >0.9999 |
| | **HPI Only** | 0 | 0 | 3 (100%) | | |
| | **Sham FUS** | 0 | 0 | 3 (100%) | | |
| | **FUS Only** | 0 | 0 | 2 (100%) | | |
| | **FUS+HPI** | 0 | 0 | 4 (100%) | | |
| | **LA+HPI** | 0 | 0 | 3 (100%) | | |
| **14** | **Control** | 0 | 0 | 3 (100%) | ns | >0.9999 |
| | **HPI Only** | 0 | 0 | 1 (100%) | | |
| | **Sham FUS** | 0 | 0 | 3 (100%) | | |
| | **FUS Only** | 0 | 0 | 2 (100%) | | |
| | **FUS+HPI** | 0 | 0 | 3 (100%) | | |
| | **LA+HPI** | 0 | 0 | 3 (100%) | | |
| **16** | **Control** | 0 | 0 | 3 (100%) | ns | >0.9999 |
| | **HPI Only** | 0 | 0 | 1 (100%) | | |

version 3

| | | | | |
|---|---|---|---|---|
| | **Sham FUS** | 0 | 0 | 3 (100%) |
| | **FUS Only** | 0 | 0 | 1 (100%) |
| | **FUS+HPI** | 0 | 0 | 2 (100%) |
| | **LA+HPI** | 0 | 0 | 3 (100%) |

**Supplementary Table 18:** Hindpaw (HP) extension testing (motor response/startle reflex assessment) across all experimental groups in female rats, including No Intervention (Control; n = 6), HPI Only (Disease Control; n = 6), Sham FUS (n = 6), FUS Only (n = 6), FUS+HPI (Intervention; n = 7), and LA+HPI (Positive Control; n = 6) at each time point up to 16 weeks. Data were analyzed using contingency table analysis followed by Fisher's exact test, and behavioral responses are presented as fractional response values. FUS = focused ultrasound; HPI = hindpaw incision; LA = local anesthetic.

version 3
